# MISO Regulates Mitochondrial Dynamics and mtDNA Homeostasis by Establishing Membrane Subdomains

**DOI:** 10.1101/2025.01.05.631328

**Authors:** Yue Zhang, Yuchen Xia, Xinhui Wang, Yueqin Xia, Shang Wu, Jianshuang Li, Xuan Guo, Qinghua Zhou, Li He

## Abstract

Mitochondrial dynamics and mtDNA homeostasis are essential for numerous biological processes and have recently been linked through distinct subdomains termed small MTFP1-enriched mitochondria (SMEM). However, the molecular mechanisms governing this connection remain unclear. Here, we identified MISO (**M**itochondrial **I**nner membrane **S**ubdomain **O**rganizer), a novel protein that regulates mitochondrial dynamics and drives SMEM formation in response to inner mitochondrial membrane (IMM) stresses. We demonstrate that MISO regulates mitochondrial dynamics in both *Drosophila* stem cells *in vivo* and mammalian cells *in vitro*. Mechanistic study showed that MISO promotes mitochondrial fission while inhibiting fusion. Notably, MISO induces SMEM formation and facilitates their peripheral fission, which regulates the lysosomal degradation of mtDNA. Meanwhile, MISO knock-out completely abolishes SMEM generation, indicating that MISO is both necessary and sufficient for SMEM biogenesis. Furthermore, SMEM functionality requires MISO-dependent recruitment of MTFP1 and subsequent engagement of downstream effectors, including the FIS1-DRP1 fission machinery. IMM stresses, including damages in mtDNA, OXPHOS complexes, and cristae, stabilize the normally short-lived MISO protein, thereby triggering SMEM assembly. Furthermore, MISO-orchestrated SMEM formation depends on its C-terminal domain, likely mediated by oligomerization. Together, our work elucidates a molecular mechanism by which IMM stresses affect mitochondrial dynamics and mtDNA homeostasis.

## Introduction

Mitochondria, the primary metabolic and signaling hubs of eukaryotic cells, play pivotal roles in a wide array of essential cellular processes. Over 40% of mitochondrial proteins have been linked to human diseases, underscoring their critical importance in human health and pathology. Recent advances in mitochondrial biology have revealed remarkable spatial heterogeneity within mitochondrial networks, with distinct subdomains exhibiting specialized functions. Among the most well-studied subdomains are endoplasmic reticulum (ER)-mitochondria contact sites, which coordinate lipid exchange, calcium signaling, mitochondrial division, and mitochondrial DNA (mtDNA) replication^1^. Other specialized membrane regions, defined by contacts between mitochondria and lysosomes, the Golgi apparatus, and lipid droplets, have also been found to regulate mitochondrial dynamics and metabolic states^2–4^. Furthermore, researchers have identified mitochondrial subpopulations dedicated to proline and ornithine metabolism^5^. Specific pathological conditions can also induce unique mitochondrial subpopulations, such as those mediating protein degradation during aging or nutrient stress^6, 7^. And *Toxoplasma gondii* infection has been found to trigger the formation of a specialized outer mitochondrial membrane (OMM) subdomain^8^. Despite these advancements, our understanding of mitochondrial subdomain diversity and functionality remains incomplete. Critical questions persist regarding whether additional functional subdomains exist, what molecule organizes them, and how their formation and activity are controlled.

A previous live-imaging study has uncovered a unique mitochondrial behavior termed peripheral fission, which is specifically regulated by mitochondrial fission protein 1 (FIS1)^9^. This process facilitates the segregation of damaged mitochondrial components into smaller fragments for targeted degradation^9^. Subsequent research revealed that the mitochondrial fission process 1 (MTFP1) protein facilitates the partitioning of mtDNA into small MTFP1-enriched mitochondria (SMEM) and promotes their lysosomal degradation through this peripheral fission mechanism^10^. However, MTFP1 is dispensable for the formation of these specialized subdomains, which are marked by prohibitin 2 (PHB2) enrichment^10^. This evidence suggests that the core regulator of SMEM biogenesis remains unidentified. Furthermore, multiple forms of mitochondrial stress, including mtDNA damage (induced by ethidium bromide treatment) and cristae disruption (via MIC60 or ATP5A knockdown), trigger SMEM formation^10^. But the molecular mechanism governing this stress-responsive process is still unknown. Our study addresses these fundamental gaps by identifying MISO as the key regulatory protein that couples mitochondrial dynamics to mtDNA homeostasis, especially under inner mitochondrial membrane (IMM) stress conditions.

## Results

### CG30159/MISO forms mitochondrial subdomains and promotes mitochondrial fragmentation in *Drosophila* intestinal stem cells

In a prior genetic screen designed to identify genes enriched in *Drosophila* intestinal stem cells (ISCs), we identified the *CG30159* gene. Both loss-of-function (LOF) and gain-of-function (GOF) of this gene significantly affected ISC proliferation^11^. Bioinformatic analysis predicts that *CG30159* encodes a putative secreted protein, given the presence of a signal peptide-like sequence at its N-terminus. Despite this prediction, the precise molecular function of CG30159 remains unknown. Its human homologue, C3orf33, has been identified in a genetic screen as a suppressor of ERK signaling; however, the mechanisms underlying this inhibitory effect are also unclear^12^.

To elucidate the biological function of *CG30159*, we first inserted a *Gal4* gene downstream of the start codon (ATG) of *CG30159* (**Fig. 1a**). Using this Gal4 transcriptionally controlled by *CG30159* promoter, we found that CG30159 is highly expressed in the *esg+* stem cells of the *Drosophila* midgut, including both intestinal stem cells (ISCs, marked by Delta staining) and enteroblasts (marked by Su(H)Gbe-LacZ staining) (**Figure 1b, Extended Data Fig. 1a-d**). To explore the subcellular localization of CG30159, we introduced a C-terminal HA-tagged version of CG30159 (CG30159-HA) into ISCs and observed that CG30159-HA formed multiple punctate structures resembling intracellular vesicles (**Fig. 1c**). To determine the precise subcellular localization of CG30159, we performed immuno-electron microscopy on fly midgut expressing CG30159-HA in ISCs, using gold particle-conjugated antibodies against the HA tag. Strikingly, all nano-gold particles were found within the mitochondria, indicating that CG30159 is a mitochondrial protein (**Fig. 1d**). This finding is supported by two independent proteomic studies using APEX proximity-labeling in *Drosophila* and mammalian cells, which identified CG30159 and its human ortholog C3orf33 as proteins localized to the mitochondrial matrix or IMM^13, 14^.

**Fig. 1.**
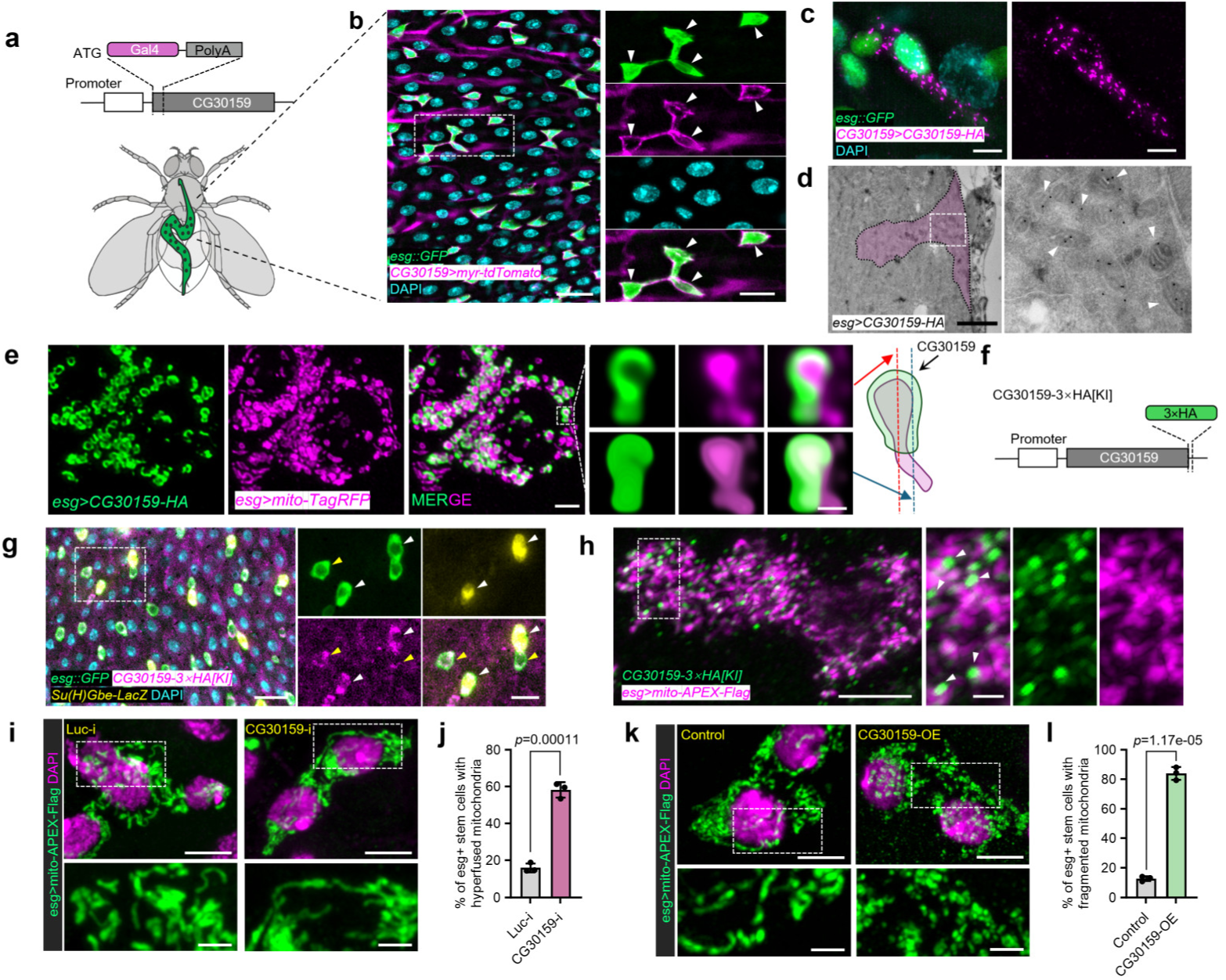
*Drosophila* MISO/CG30159 forms mitochondrial subdomains and regulates mitochondrial dynamics. **a-b.** *CG30159* is expressed in *escargot* (*esg*)-positive stem cells (green) in the adult fly intestine by the Gal4 knock-in element. Scale bars: main panels 20 μm, magnified insets 10 μm. **c.** HA-tagged CG30159 in stem cells exhibits multiple punctate patterns. Scale bars: 5 μm. **d.** Immuno-electron microscopy (EM) analysis of CG30159 (*esg>CG30159-HA*) localization in the *Drosophila* intestine. White arrowheads mark immunogold-labeled CG30159. The red pseudo-color indicates stem cells. **e.** Representative images of CG30159 immunofluorescence staining in *esg*+ stem cells. mito-TagRFP is used as a marker for mitochondria matrix. Scale bar: 2 μm. Magnified insets display the original data of a single subdomain of CG30159 within mitochondria (upper panel) and corresponding 3D surface reconstructions (lower panel). Scale bar: 500 nm. **f.** A diagram of *CG30159* gene structure with 3×HA tag knock-in element. **g.** Representative images showing endogenous CG30159 protein expression in fly intestine. Yellow arrowheads indicate stem cells, and white arrowheads indicate enteroblasts. Scale bars: main panels 20 μm, magnified insets 10 μm. **h.** Representative images depicting the localization of endogenous CG3015 in intestinal stem cells. White arrowheads point to membrane subdomains. Scale bars: main panels 5 μm, magnified insets 1 μm.

Considering that mitochondria in ISCs typically exhibit more filamentous structures, this raises questions regarding the punctate appearance of CG30159. To investigate this, we performed co-labeling of CG30159 and mitochondria in ISCs and analyzed the samples using high-resolution microscopy. Surprisingly, the CG30159 puncta were found to form “bud-like” membrane subdomains encircling the tips of mitochondria (**Fig. 1e, Extended Data Fig. 1e**). This specific localization of CG30159 to mitochondrial subdomains was further confirmed in cultured *Drosophila* S2 cells, where a different tag (3×Flag) was introduced at the N-terminus of CG30159, indicating that the observed localization is not an artifact of the tag fusion (**Extended Data Fig. 1f**). Given that some proteins may form unnatural aggregates when overexpressed, we further generated a knock-in fly with a 3×HA tag at the C-terminus of endogenous CG30159 (**Fig. 1f**). Our data showed that the CG30159-HA[KI] protein is enriched in the midgut stem cells of flies (**Fig. 1g**) and localizes to numerous punctate subdomains on mitochondria at endogenous expression levels (**Fig. 1h**). These endogenously tagged subdomains are similar in location to those induced by overexpression, albeit smaller in size, suggesting that CG30159 naturally accumulates in specific regions of mitochondria.

Given that CG30159 is a mitochondrial protein, we further investigated the effects of modulating its expression levels on mitochondrial morphology in ISCs. Knocking down CG30159 in ISCs resulted in an elongated mitochondrial phenotype (**Fig. 1i-j, Extended Data Fig. 1g**), whereas overexpression of CG30159 induced pronounced mitochondrial fragmentation (**Fig. 1k-l**). These results demonstrate that CG30159 acts as a novel regulator of mitochondrial dynamics. Given its ability to form mitochondrial subdomains (**Fig. 1e, h**, **Extended Data Fig. 1e, f**), we rename CG30159 as the **M**itochondrial **I**nner membrane **S**ubdomain **O**rganizer (MISO) and refer to it as MISO in all subsequent studies.

### Mammalian MISO/C3orf33 is a novel regulator of mitochondrial morphology

MISO is an evolutionarily conserved protein present in major metazoan species (**Fig. 2a**). A typical MISO protein is composed of 240-300 amino acids (AAs) and includes three distinct regions: an N-terminal region (∼20-60 AAs) comprising a transmembrane domain (previously annotated as a signal peptide for secretion), a highly conserved middle region (∼120 AAs) predicted to contain a core beta-barrel structure, and a less conserved C-terminal region (∼70-100 AAs) that contains 3-4 predicted alpha-helices (**Fig. 2a, Extended Data Fig. 2a**). In humans, two isoforms of MISO/C3orf33 have been predicted, with the shorter isoform lacking the first 43 AAs at the N-terminus. Truncation analysis revealed that the N-terminal region containing the transmembrane domain (residues 1-59) of the long isoform is sufficient for mitochondrial targeting (**Extended Data Fig. 2b-c**). However, a close examination showed that the N-terminal fragment (residues 1-59) localizes predominantly to the outer mitochondrial membrane (OMM). Correct localization to the IMM requires both the N-terminal and middle region (residues 1-216), indicating that the middle region is essential for proper IMM localization (**Extended Data Fig. 2d-g**).

**Fig. 2.**
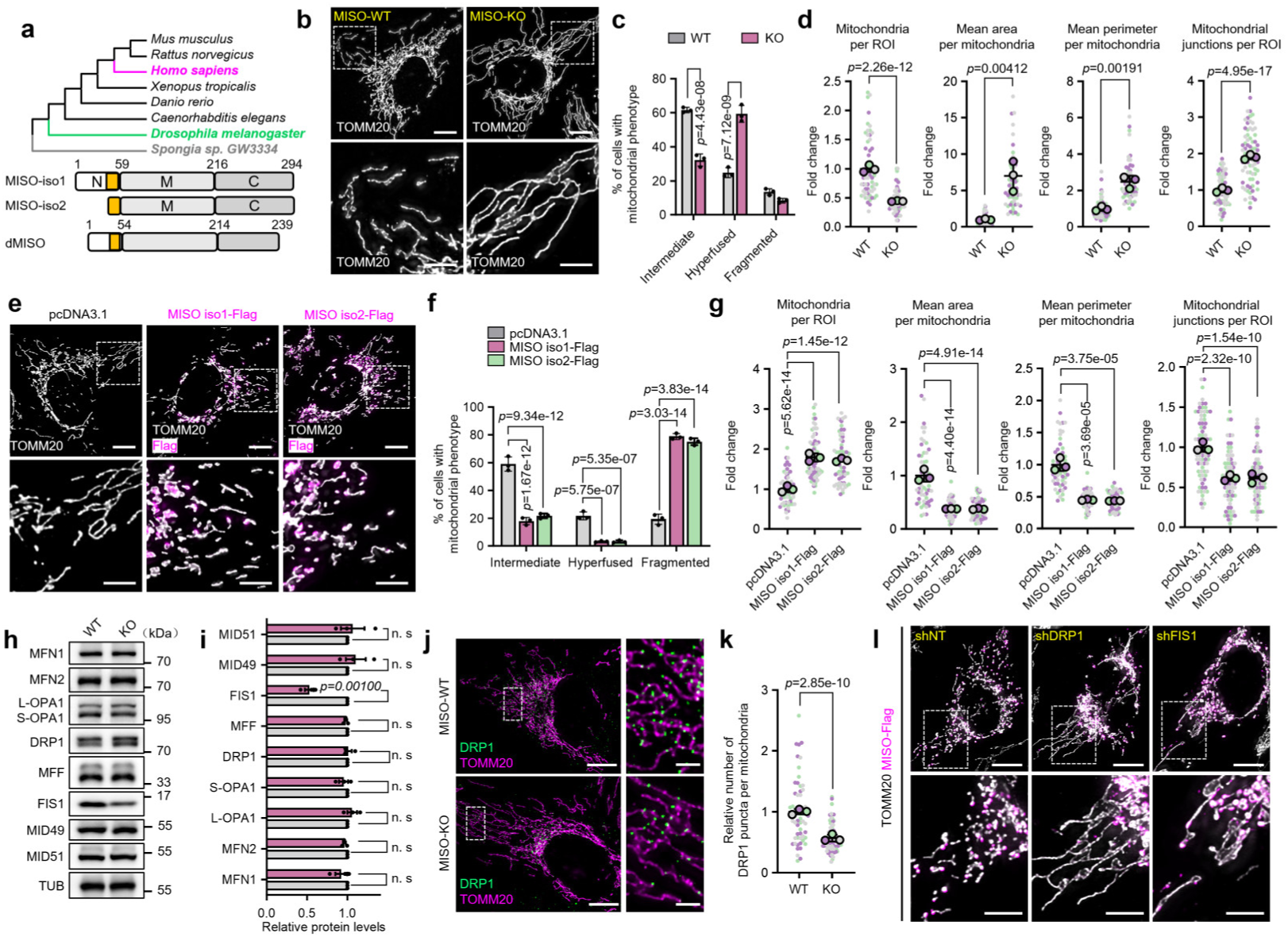
Mammalian MISO/C3orf33 is a novel regulator of mitochondrial morphology. **a.** MISO is evolutionarily conserved across major metazoan species, spanning from sea sponges to humans. The typical transmembrane domains (44-59 amino acids, human) are highlighted in yellow. **b-c.** Representative images (**b**) and corresponding quantification (**c**) of mitochondrial morphology in wild-type (WT) and MISO-knockout (KO) U2OS cells. Scale bars: main panels 10 μm, magnified insets 5 μm. *n* = three experiments. **d.** Quantification of mitochondrial parameters from (**b**). *n* = three experiments. **e-f.** Representative images (**e**) and corresponding quantification (**f**) of mitochondrial morphology in U2OS cells overexpressing an empty vector (pcDNA3.1), MISO isoform1-Flag or MISO isoform2-Flag. Scale bars: main panels 10 μm, magnified insets 5 μm. *n* = three experiments. **g.** Quantification of mitochondrial parameters from (**e**). *n* = three experiments. **h-i.** Representative immunoblots (**h**) and corresponding quantifications (**i**) of proteins involved in mitochondrial fission and fusion from WT and MISO-KO U2OS cells. *n* = three experiments. **j.** Representative images of DRP1 localization in WT and MISO-KO U2OS cells. Scale bars: main panels 10 μm, magnified insets 2 μm. **k.** Quantification of DRP1 puncta on TOMM20-labeled mitochondria. *n* = three experiments. **l.** Representative images of mitochondrial morphology in U2OS cells expressing MISO-Flag and treated with the indicated shRNAs. NT: non-targeted. Scale bars: main panels 10 μm, magnified insets 5 μm. All data are presented as mean ± SD. (**c**) and (**f**): two-way ANOVA with Tukey’s multiple comparisons test; (**d**) and (**k**): two-tailed nested t test; (**j**): nested ordinary one-way ANOVA with Tukey’s multiple comparisons test; (**i**): two-tailed unpaired t test. n. s: not significant.

To investigate the biological function of MISO, we generated MISO knockout (MISO-KO) human osteosarcoma U2OS cells using CRISPR-Cas9 to disrupt both MISO isoforms (**Extended Data Fig. 3a-b**). Similar to the observations in *Drosophila* ISCs, mitochondria in MISO-KO cells exhibited a significantly elongated phenotype (**Fig. 2b-d, Extended Data Fig. 3c-d**). The knockout phenotype is rescued by the expression of either MISO isoform (**Extended Data Fig. 3e-g**). Meanwhile, overexpression of either isoform of MISO in wild-type (WT) cells induced a comparable degree of mitochondrial fragmentation, indicating that both isoforms play similar roles in regulating mitochondrial dynamics (**Fig. 2e-g**). Additionally, both isoforms localized to similar subdomains within the IMM (**Extended Data Fig. 4a-e**), further supporting their functional redundancy. Given these similarities, we primarily used the long isoform of MISO in subsequent experiments and refer to it simply as MISO unless otherwise noted.

Additionally, knocking down MISO using two independent shRNAs targeting both isoforms resulted in similar mitochondrial elongated phenotypes in U2OS cells, as well as in human hepatoma PLC/PRF/5 and Huh7 cells (**Extended Data Fig. 5**), indicating that MISO’s role in mitochondrial regulation is conserved across these human cell types. To examine MISO’s function *in vivo*, we generated a MISO-KO mouse strain using CRISPR-Cas9 combined with germline Cre-dependent excision (**Extended Data Fig. 6a-b**). These systemic MISO-KO mice exhibited no significant developmental or reproductive abnormalities. However, mitochondria in liver tissue, as well as in primary hepatocytes and mouse embryonic fibroblasts (MEFs) from MISO-KO mice, displayed a significantly elongated morphology compared to those from MISO^loxp/loxp^ mice (**Extended Data Fig. 6c-h**).

To investigate whether MISO is required for mitochondrial fission, we tested the effects of overexpressing the mitochondrial fission factor (MFF) and treating cells with the mitochondrial stressor carbonyl cyanide m-chlorophenyl hydrazone (CCCP). Both treatments induced comparable levels of mitochondrial fragmentation in WT and MISO-KO cells, suggesting that MISO is not an indispensable component of mitochondrial fission (**Extended Data Fig. 7a-d**). This modulatory role of MISO may be attributed to its relatively late evolutionary emergence in multicellular organisms.

To investigate how MISO loss affects mitochondrial dynamics, we analyzed the protein levels of key regulators of mitochondrial fission and fusion by Western blotting. Notably, mitochondrial fission protein 1 (FIS1) was significantly reduced in MISO-KO cells (**Fig. 2h-i**). Previous studies have shown that FIS1, together with dynamin-related protein 1 (DRP1), plays a critical role in mediating peripheral fission to generate small daughter mitochondria^9, 10^. We next examined DRP1 recruitment to mitochondria in MISO-deficient cells and found that the number of DRP1 puncta associated with mitochondria was significantly decreased in MISO-KO cells compared to WT controls (**Fig. 2j-k**), suggesting that the elongation phenotype may result from impaired DRP1 recruitment. Furthermore, knockdown of either DRP1 or FIS1 markedly suppressed the mitochondrial fragmentation induced by MISO overexpression (**Fig. 2l, Extended Data Fig. 7e-f**), supporting that MISO promotes mitochondrial fission by regulating FIS1 and DRP1.

### MISO-Enriched mitochondrial subdomains play dual roles in regulating mitochondrial dynamics

The most striking feature of MISO is its ability to form specific subdomains on mitochondria, a phenomenon consistently observed in both *Drosophila* and various mammalian cell types, including U2OS, HeLa, Huh7, A549, PLC, and HEK293T cells (**Fig. 1d**, **Fig. 3a, Extended Data Fig. 8a**). To rule out potential artifacts arising from epitope tagging, we generated a polyclonal antibody against human MISO. Although this antibody failed to detect MISO at endogenous levels, it successfully recognized overexpressed MISO (**Extended Data Fig. 8b**). Immunostaining of overexpressed untagged MISO in U2OS cells revealed mitochondrial subdomain formation similar to that seen with tagged versions (**Extended Data Fig. 8c**), suggesting that subdomain formation is not an artifact of tag fusion. To investigate whether endogenous MISO forms subdomains in mammals, we generated a knock-in mouse with a 3×Flag tag inserted at the C-terminus of mouse MISO (mMISO) (**Extended Data Fig. 9a**). Western blot analysis confirmed that mMISO-Flag is expressed across multiple mouse tissues, with particularly high levels in energy-demanding organs such as the brain, kidney, and brown adipose tissue (**Extended Data Fig. 9b**). To visualize endogenous mMISO localization, we isolated mouse embryonic fibroblasts (MEFs) from these knock-in mice and performed immunostaining using anti-Flag antibodies. Our data showed that mMISO-Flag localizes to mitochondria and forms distinct subdomains closely resembling those observed in *Drosophila* (**Fig. 3b**). These findings confirmed that endogenous MISO forms mitochondrial subdomains in mammalian cells.

**Fig. 3.**
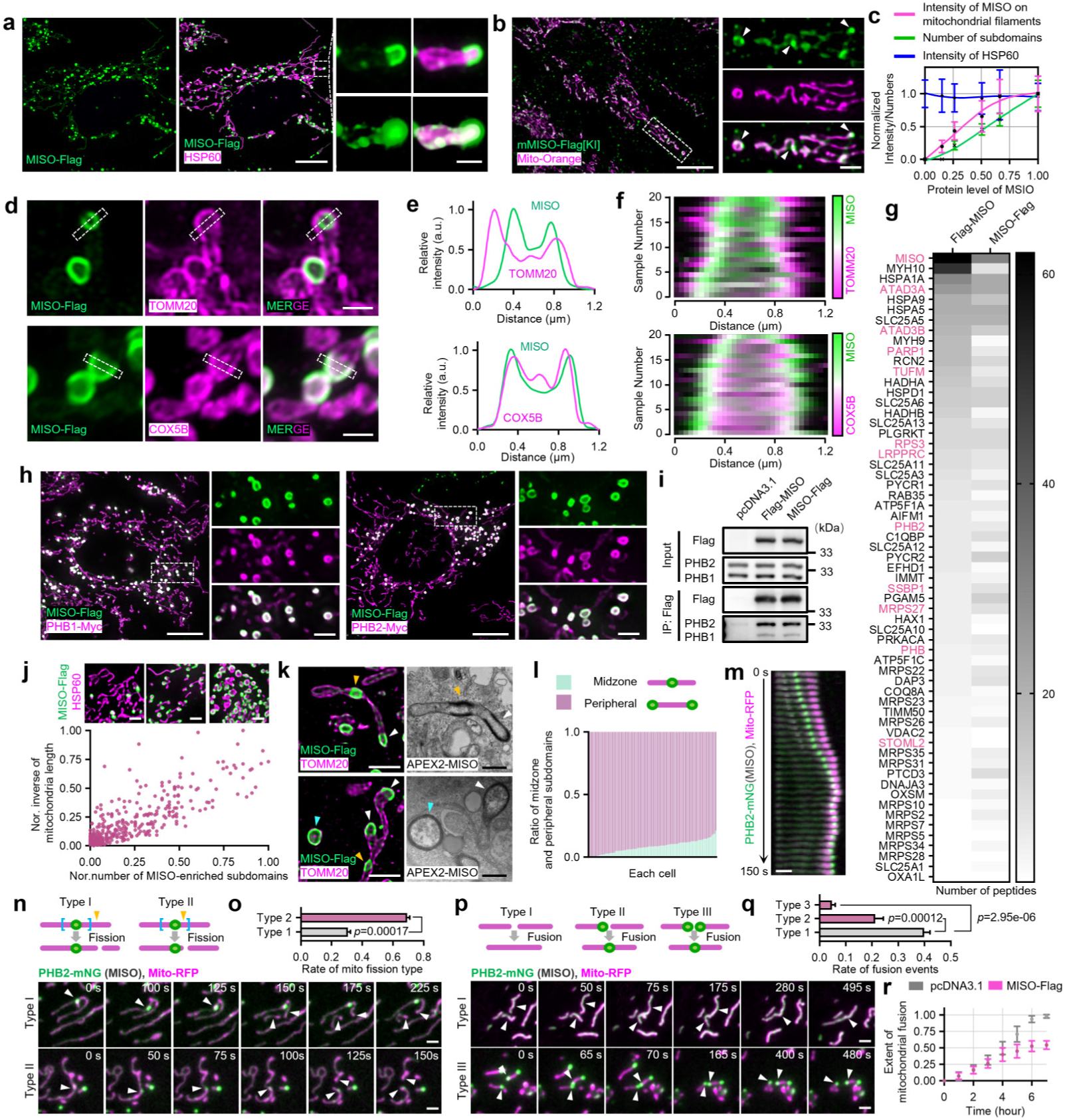
MISO-enriched subdomains exert dual regulatory control over mitochondrial dynamics. **a.** Representative images depicting the localization of MISO in U2OS cells. The insets highlight the raw data pertaining to the mitochondrial subdomains (upper panels) and feature 3D surface reconstructions (lower panels). Scale bars: main panels 10 μm, magnified insets 1 μm. **b.** Representative images depicting the localization of endogenous MISO in MEFs. White arrowheads indicate mitochondrial subdomains. Scale bars: main panels 10 μm, magnified insets 2 μm. **c.** The intensity of MISO on mitochondrial filaments and the number of mitochondrial subdomains are plotted at varying levels of MISO expression. **d.** Representative images confirming MISO-enriched subdomains localization to the inner mitochondrial membrane in U2OS cells expressing MISO-Flag. Immunofluorescence staining was performed to detect the Flag epitope (green) and co-stained with TOMM20 (outer membrane, upper panels, magenta) or COX5B (inner membrane, lower panels, magenta). Scale bars: 1 μm. **e.** Line profiles showing fluorescent signals of each channel across subdomains marked by the dotted rectangles in (**d**). **f.** Fluorescence colocalization analysis of MISO (green) in mitochondrial subdomains with TOMM20 (upper panel, magenta) or COX5B (lower panel, magenta). **g.** Heatmap depicting the major mitochondrial protein of Flag-MISO and MISO-Flag interactomes analyzed via liquid chromatography-mass spectrometry (LC-MS). Color intensity reflects the number of identified peptides. **h.** Fluorescence colocalization analysis of MISO with PHB1-Myc and PHB2-Myc. Scale bars: main panels 10 μm, magnified insets 2 μm. **i.** Immunoblots of the indicated proteins from anti-Flag immunoprecipitation of HEK293T cells overexpressing either an empty vector (pcDNA3.1), Flag-MISO or MISO-Flag. **j.** Representative images and scatter plot illustrating the association between mitochondrial length and the number of MISO-enriched subdomains per ROI in cells expressing MISO-Flag. Scale bars: 2 μm. **k.** Representative fluorescence (left panel) and electron microscopy (right panel) images of midzone and peripheral subdomains in U2OS cells. Arrowheads: white for peripheral subdomains, yellow for midzone subdomains, and blue for isolated subdomains. Scale bar: fluorescence images 2 μm, EM images 500nm. **l.** Histogram showing the distribution of midzone and peripheral MISO-enriched subdomains proportions per cell in cells expressing MISO-Flag. **m.** Kymograph depicting the migration of MISO-enriched subdomains toward the mitochondrial periphery. Scale bar: 2 µm. **n.** Representative time-lapse images of mitochondrial fission events in U2OS cells expressing MISO-Flag and PHB2-mNeonGreen (indicating MISO) (green) along with Mito-RFP (magenta). White arrowheads point to subdomains. Type I fission: Fission event occurring within a 0-1 μm range from the subdomain spot. Type II fission: Fission event occurring within a 1-2 μm range from the subdomain spot. Scale bar: 2 μm. **o.** Quantitative evaluation of the rates of Type I and Type II mitochondrial fission events. *n* = four experiments. **p.** Representative time-lapse images of mitochondrial fusion events in U2OS cells expressing MISO-Flag and PHB2-mNeonGreen (indicating MISO) (green) along with Mito-RFP (magenta). White arrowheads point to MISO-enriched subdomains. Type I fusion: Fusion between two mitochondrial termini without the subdomains. Type II fusion: Fusion between two mitochondrial termini with subdomains present on one terminus. Type III fusion: Fusion between two mitochondrial termini with subdomains present on both termini. Scale bar: 2 μm. **q.** Quantitative assessment of mitochondrial fusion probabilities across three types of encounter events. *n* = three experiments. **r.** Mitochondrial fusion kinetics were quantitatively analyzed by measuring the proportion of overlapping mitochondria relative to total mitochondrial content in at hybrid cells per time point. *n* = three experiments. All data are presented as mean ± SD. (**o**): two-tailed unpaired t test; (**q**): ordinary one-way ANOVA with Tukey’s multiple comparisons test.

We next investigated the quantitative relationship between MISO protein levels and the formation of MISO-enriched mitochondrial subdomains using a Tet-On inducible expression system. Notably, these subdomains became detectable at the earliest stage of induction, even when MISO levels on the bulk mitochondrial network remained barely detectable (**Fig. 3c, Extended Data Fig. 10**). As MISO expression increased, the signal intensity on the general mitochondrial population plateaued, whereas the number of subdomains continued to rise linearly with MISO protein levels (**Fig. 3c, Extended Data Fig. 10)**. This dose-dependent increase in subdomain formation suggests that these structures act as dynamic, functional units that scale with MISO expression, underscoring their potential role in mediating MISO’s biological activities.

Recent studies have shown that the IMM may herniate into the cytosol during mitophagy^15, 16^, prompting us to examine the precise localization of MISO within and outside the subdomain region using STED super-resolution imaging. Our analysis revealed that MISO signals, both within the subdomains and in general mitochondria, colocalized with the IMM marker COX5B and were encapsulated by the OMM marker TOMM20 (**Fig 3. d-f, Extended Data Fig. 11a-c**). However, a small subset of MISO signals in the MISO-enriched subdomains (about 5 %) exhibited exceptions, which we will discuss in the following section. In addition to imaging data, biochemical evidence from proteinase K protection and alkaline sodium carbonate solubilization assays further supports MISO’s identity as an integral IMM protein (**Extended Data Fig. 11d-f**).

To identify proteins presented in the MISO-enriched subdomains, we performed immunoprecipitation of MISO from cell lysates followed by liquid chromatography–tandem mass spectrometry (LC-MS/MS) analysis (**Fig. 3g**). Candidate proteins were selectively validated using immunostaining with available antibodies or cDNA clones (**Extended Data Fig. 12a**). Among the tested proteins, prohibitin 1 and 2 (PHB1 and PHB2) emerged as the most prominent interactors, with endogenous PHB2 showing significant colocalization with MISO-enriched subdomains (**Fig. 3h, Extended Data** Fig. 12b-c). Co-immunoprecipitation experiments confirmed that MISO interacts with both PHB1 and PHB2, with a notably stronger association observed for PHB2 (**Fig. 3i, Extended Data Fig. 12d**). PHB1 and PHB2 are integral IMM proteins that function as heterodimers in mitochondrial membrane organization^17, 18^. Given their enrichment in these subdomains, we investigated whether PHB expression is required for the formation of MISO-enriched subdomains. However, knockdown of either PHB1 or PHB2 did not impair MISO-induced subdomain formation, indicating that PHBs are dispensable for the assembly of these structures (**Extended Data Fig. 13a-b**).

PHBs have been shown to regulate mitochondrial dynamics through multiple mechanisms, including binding and organizing the mitochondrial phospholipid cardiolipin, modulating the processing of the mitochondrial fusion protein optic atrophy 1 (OPA1), and interacting with DRP1^19–21^. Given the enrichment of PHBs within MISO-enriched subdomains, we investigated whether the formation of these subdomains promotes localized mitochondrial fragmentation. We found that the number of MISO-enriched subdomains per cell is inversely correlated with average mitochondrial length (**Fig. 3j, Extended Data Fig. 14**), suggesting a link between subdomain formation and mitochondrial fission. Notably, while these MISO-enriched subdomains can be found at both the midzone and terminals of mitochondria, they exhibit a strong preference for mitochondrial ends (**Fig. 3k-l**). These findings suggest that the localization of MISO-enriched subdomains at mitochondrial termini may facilitate peripheral fission.

To investigate the real-time effects of MISO-enriched subdomains, we initially attempted to tag MISO with GFP for live-cell imaging. However, fusion of GFP to the N-terminus, C-terminus, or internal linker regions of MISO disrupted subdomain formation or resulted in mislocalization to the ER (data not shown). Given the strong association between MISO and PHBs, we adopted an alternative strategy. While PHB2-mNeonGreen (PHB2-mNG) alone uniformly labels the mitochondrial network, it forms distinct subdomains when co-expressed with MISO (**Extended Data Fig. 15a**). This robust recruitment of PHB2-mNG to MISO-enriched subdomains enabled us to visualize their dynamics in live cells. Interestingly, we observed that MISO-enriched subdomains formed at the midzone of mitochondria actively move along the mitochondrial tubules and eventually stabilize at the mitochondrial termini (**Fig. 3m, Extended Data Fig. 15b, Supplementary Video 1**), explaining the preferential enrichment of these subdomains at mitochondrial ends.

We further investigated the spatiotemporal relationship between MISO-enriched subdomains and mitochondrial fission and fusion events. Our data revealed that mitochondria are twice as likely to undergo fission in regions close to MISO-enriched subdomains structures (within 1 micron) compared to regions farther away (1-2 microns) (**Fig. 3n-o, Supplementary Video 2**). Given that these subdomains are predominantly localized at the mitochondrial periphery, this data suggests that the formation of MISO-enriched subdomains facilitates peripheral fission. Indeed, we observed a clear separation of MISO-enriched subdomains from mitochondria, generating small fragments consistent with previously reported peripheral fission events^9, 10^ (**Extended Data Fig. 15c, Supplementary Video 3**). Additionally, we also found that mitochondria with MISO-enriched subdomains at their termini exhibited a significantly reduced likelihood (up to 90%) of fusing with other mitochondria, indicating that these subdomains also inhibit fusion (**Fig. 3p-q, Supplementary Video 4**). We further validated this inhibitory effect using polyethylene glycol (PEG)-mediated cell fusion experiments^22^. Cells overexpressing MISO showed significantly reduced mitochondrial mixing after induced cell fusion (**Fig. 3r, Extended Data Fig. 15d**).

Mechanistically, we found that the IMM fusion protein OPA1 is excluded from MISO-enriched subdomains, while the distribution of OMM protein Mitofusin 1 and 2 (MFN1 and MFN2) remains unaffected (**Extended Data Fig. 16a-b**), suggesting that these subdomains inhibit fusion likely through OPA1 exclusion. Additionally, overexpression of MFN1 or MFN2 reversed the fragmentation phenotype caused by MISO overexpression, suggesting that MISO does not impair OMM fusion capacity (**Extended Data Fig. 16c-d**). Conversely, knockdown of MFN1, MFN2, or OPA1 reversed the elongation phenotype in the MISO-KO cells, demonstrating that MISO’s effects on mitochondrial morphology depend on the canonical fusion machinery (**Extended Data Fig. 16e-g**). Together, these findings establish MISO-enriched subdomains as regulatory platforms that coordinately promote mitochondrial fission and suppress fusion.

Previous studies have shown that mitochondrial fission is frequently associated with membrane contact sites between mitochondria and other organelles, including the ER, Golgi apparatus, and lysosomes^3, 4, 23, 24^. To determine whether MISO-enriched subdomains are spatially associated with specific organelles, we analyzed their co-localization. While these subdomains are predominantly localized within mitochondria, they also exhibit a moderate degree of association with the ER compared to other organelles (**Extended Data Fig. 17**). However, the functional significance of this association remains to be determined through future functional studies.

### MISO-enriched subdomains are SMEMs that regulate mtDNA homeostasis

Our proteomic analysis identified several proteins involved in mitochondrial nucleoid organization that associate with MISO, including poly ADP-ribose polymerase 1 (PARP1), Tu translation elongation factor (TUFM), single-stranded DNA-binding protein 1 (SSBP1), ribosomal protein S3 (RPS3), ATPase family AAA domain-containing protein 3A (ATAD3A), and mitochondrial ribosomal protein S27 (MRPS27) (**Fig. 3g**). Co-localization between MISO and mtDNA and the mtRNA-associated protein FAST kinase domains-containing 2 (FASTKD2) demonstrate a close spatial association between mitochondrial nucleoids and MISO-enriched subdomains (**Fig. 4a-b, Extended Data Fig. 18a**). Notably, nearly all MISO-enriched subdomains contain nucleoids, independent of mtDNA-damaging reagent treatment, suggesting an intrinsic strong affinity (**Extended Data Fig. 18b-c**). Furthermore, immunostaining for mitochondrial transcription factor A (TFAM), a major mtDNA-binding protein, revealed that mtDNA is clearly enriched within MISO-enriched subdomains (**Extended Data Fig. 18d**). Live-cell imaging of MISO-enriched subdomains labeled with PHB1-mCherry also exhibited strong co-variation with nucleoids stained with PicoGreen (**Fig. 4c, Extended Data Fig. 18e, Supplementary Video 5**). An EdU labeling assay further indicates that nucleoids within these subdomains are actively replicating (**Extended Data Fig. 18f**). Additionally, SSBP1, a core mtDNA replication factor^25^, localizes to MISO-enriched subdomains and co-immunoprecipitates with MISO (**Extended Data Fig. 18g-h**), suggesting that MISO may recruit mitochondrial nucleoids to the subdomains through interaction with SSBP1. To test if SSBP1 binding or mtDNA recruitment is required for the formation of MISO-enriched subdomains, we knocked down SSBP1 in cells overexpressing MISO. Depletion of SSBP1 has been shown to cause a marked decrease in mtDNA^26^. However, subdomain formation was unaffected by SSBP1 knock-down, indicating that the assembly of MISO-enriched subdomains does not depend on SSBP1 (**Extended Data Fig. 18i-j**). Given that MISO-enriched subdomains preferentially form at mitochondrial termini, we investigated whether they might affect the spatial distribution of mtDNA. Indeed, MISO overexpression markedly enhanced the probability of mtDNA localization to mitochondrial ends (**Fig. 4d-e**), suggesting that these MISO-enriched membrane subdomains may function as a positioning system that directs mtDNA to mitochondrial termini.

**Fig. 4.**
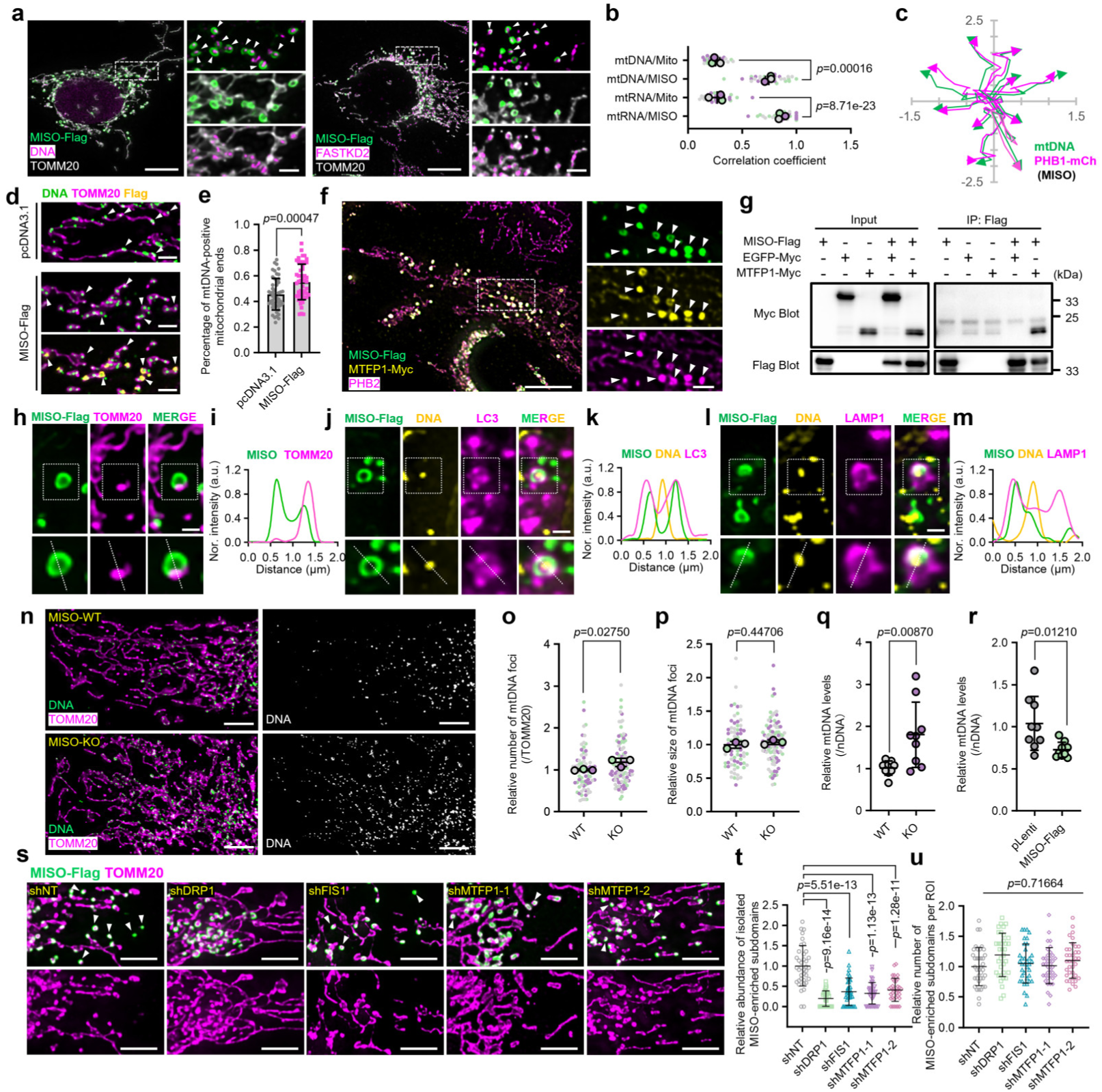
MISO constitutes SMEM that regulate mtDNA homeostasis. **a.** Representative images depicting the association between mtDNA or mtRNA (indicated by FASTKD2 staining) with MISO-enriched subdomains in U2OS cells. Scale bars: main panels 10 μm, magnified insets 2 μm. **b.** Quantification of correlation coefficient between MISO-enriched subdomains and mtDNA or mtRNA in cells shown in (**a**). *n* = three experiments. **c.** Migration paths of mitochondrial DNA (PicoGreen, green) and mCherry-PHB1 (representing MISO, magenta) were tracked using MTrack2 in Fiji software. **d-e.** Representative images (**d**) and corresponding quantifications (**e**) of mtDNA-positive mitochondrial ends in U2OS cells expressing empty vector (pcDNA3.1) or MISO-Flag. White arrows indicate mtDNA-positive mitochondrial ends. Scale bars: 2 μm. **f.** Representative images showing the colocalization of MISO, MTFP1, and PHB2. Scale bars: main panels 10 μm, magnified insets 2 μm. **g.** Co-immunoprecipitation with anti-Flag validate the interaction between MISO and MTFP1. **h-i.** Representative images (**h**) and line profiles (**i**) showing the absence of TOMM20 within MISO-enriched subdomains. Scale bars, 1 μm. **j-k.** Representative images (**j**) and line profiles (**k**) showing the presence of MISO-enriched subdomains within autolysosome structures marked by LC3. Scale bars, 1 μm. **l-m.** Representative images (**l**) and line profiles (**m**) showing the presence of MISO-enriched subdomains within autolysosome structures marked by LAMP1. Scale bars, 1 μm. **n.** Representative images of mtDNA and mitochondria in WT and MISO-KO U2OS cells. Scale bars, 5 μm. **o-p.** Quantification of the number (**o**) and size (**p**) of mtDNA foci per ROI from (**i**). *n* = three experiments. **q.** qPCR analysis of mtDNA copy number in WT and MISO-KO U2OS cells. *n* = nine experiments. **r.** qPCR analysis of mtDNA copy number in U2OS cells expressing an empty vector (pLenti) or MISO-Flag. *n* = nine experiments. **s.** Representative images of MISO-enriched subdomains in U2OS cells treated with the indicated shRNAs. White arrows indicate isolated MISO-enriched subdomains. Scale bars: main panels 10 μm, magnified insets 5 μm. **t-u.** Quantification of the abundance of isolated (**t**) and total (**u**) MISO-enriched subdomains from (**s**). All data are presented as mean ± SD. (**b**), (**o**) and (**p**): two-tailed nested t test; (**e**), (**q**) and (**r**): two-tailed unpaired t test; (**t**) and (**u**): ordinary one-way ANOVA with Tukey’s multiple comparisons test.

Recently, the mitochondrial fusion inhibitor MTFP1 was discovered to regulate mtDNA degradation by participating in the formation of a specialized mitochondrial subdomain termed small MTFP1-enriched mitochondria (SMEM)^10^. Like the MISO-enriched subdomains, SMEM enriched in PHBs eventually separate from the main mitochondria through peripheral fission^10^. These similarities make us wonder if the MISO-enriched subdomains and SMEMs are the same structure. Indeed, we observed clear co-localization of MISO, MTFP1, and PHB2 within the same mitochondrial subdomains (**Fig. 4f**). Immunoprecipitation experiments further confirmed a physical interaction between MISO and MTFP1 (**Fig. 4g**).

Consistent with previous report about SMEM, MISO-enriched subdomains (marked by PHB2-mNG) exhibit elevated mitochondrial ROS levels (detected by MitoSOX) and reduced membrane potential (measured by TMRM) (**Extended Data Fig. 19a**). Supporting these observations, Seahorse assays and TMRM-based measurements revealed that both oxidative phosphorylation (OXPHOS) activity and mitochondrial membrane potential are enhanced in MISO-KO cells but diminished in MISO-overexpressing cells (**Extended Data Fig. 19b-f**). These results suggest that MISO modulates OXPHOS function. Indeed, Western blot analysis of OXPHOS complexes showed an inverse correlation between MISO expression and Complex IV levels in U2OS cells (**Extended Data Fig. 20a-b**). Furthermore, liver tissue from MISO-KO mice displayed increased protein levels and enzymatic activities of the OXPHOS complexes (**Extended Data Fig. 20c-f**).

Previous studies have shown that SMEMs are targeted to autophagic structures, and both PHB2 and MTFP1 have been identified as IMM receptors for the autophagosome-associated protein LC3^10, 27^. In line with this, we observed that MISO-bearing vesicles exhibited signs of OMM rupture, as indicated by the loss of TOMM20 staining (**Fig. 4h-i**). Notably, TOMM20 signal loss was occasionally detected in these subdomains even prior to their separation from the main mitochondrial network via peripheral fission (**Extended Data Fig. 21**), suggesting that, like SMEMs, MISO-enriched subdomains may serve as sites for selective mitochondrial degradation. Indeed, these MISO-positive mitochondrial fragments contained mtDNA and were found to associate with autophagy markers LC3 and lysosome-associated membrane protein 1 (LAMP1) (**Fig. 4j-m**), supporting their trafficking to lysosomal organelles. Together, these findings demonstrate that MISO-enriched subdomains share identical molecular markers, morphological characteristics, and trafficking fates with the previously identified SMEMs, indicating that they represent the same cellular structure. Hence, we will refer to MISO-enriched microdomains as SMEMs in subsequent studies.

As SMEMs have been shown to regulate mtDNA homeostasis, we investigated whether MISO is required for this function. Indeed, MISO-KO cells exhibited increased nucleoid staining compared to WT cells, with minimal change in nucleoid size (**Fig. 4n-p**). Quantitative PCR analysis confirmed a higher relative mtDNA copy number (mtDNA CN) normalized to nuclear DNA in MISO-KO cells (**Fig. 4q**). Elevated mtDNA CN was also observed in primary hepatocytes and MEFs, as well as in multiple tissues from MISO-KO mice, compared to controls *mMISO^flox/flox^* mice (**Extended Data Fig. 22a-c**). Conversely, MISO overexpression in U2OS cells resulted in reduced mtDNA CN (**Fig. 4r**). These results collectively demonstrate that MISO plays a role in regulating mtDNA homeostasis both *in vitro* and *in vivo*.

We next investigated the functional relationship between MISO and MTFP1. Consistent with previous findings showing that MTFP1 is dispensable for SMEM formation^10^, knocking down MTFP1 does not affect the generation of SMEM triggered by MISO overexpression (**Extended Data Fig. 23a-b**). However, depletion of MTFP1 reduced the number of segmented SMEMs to a similar extent as knockdown of the core FIS1-DRP1 fission machinery under MISO overexpression (**Fig. 4s-u**), indicating that MTFP1 is essential for the peripheral fission of SMEMs following subdomain formation. These results position MTFP1 as a key downstream effector in this pathway. Indeed, MTFP1 knockdown markedly suppressed the mitochondrial fragmentation phenotype induced by MISO overexpression (**Extended Data Fig. 23c-d**), while MTFP1 overexpression rescued the mitochondrial elongation observed in MISO-KO cells (**Extended Data Fig. 23e-g**). Together, these findings establish MISO as an upstream organizer of SMEM biogenesis, with MTFP1 acting as a critical downstream effector mediating peripheral fission.

### IMM stress induces SMEM formation by stabilizing the MISO protein

Previous studies have shown that treatment with ethidium bromide (EB)— which disrupts mtDNA replication and promotes nucleoid loss—induces the formation of SMEM^10, 28^. As demonstrated above, MISO levels directly correlate with the number of MISO-enriched subdomains, now identified as SMEM. This led us to hypothesize that MISO may be the missing factor linking IMM stresses to SMEM formation. Indeed, our results show that MISO knockout completely blocks EB-induced SMEM formation, as evidenced by both PHB2 immunostaining and the absence of IMM alterations observed by transmission electron microscopy (TEM), suggesting that MISO is required for SMEM biogenesis (**Fig. 5a-e**). Notably, MISO knockout did not alter the expression levels of MTFP1 or PHBs, suggesting that the loss of SMEMs is likely due to the absence of MISO rather than secondary changes in other SMEM-associated proteins (**Extended Data** Fig. 24a-b). Furthermore, using MEF cells derived from mMISO-Flag knock-in mice, we found that EB treatment significantly increased mMISO protein levels. The levels of PHBs and MTFP1 remained unchanged upon EB treatment (**Fig. 5f-g, Extended Data** Fig. 24a-b), further supporting a unique and central role for MISO in driving SMEM formation in response to mitochondrial stress.

**Fig. 5.**
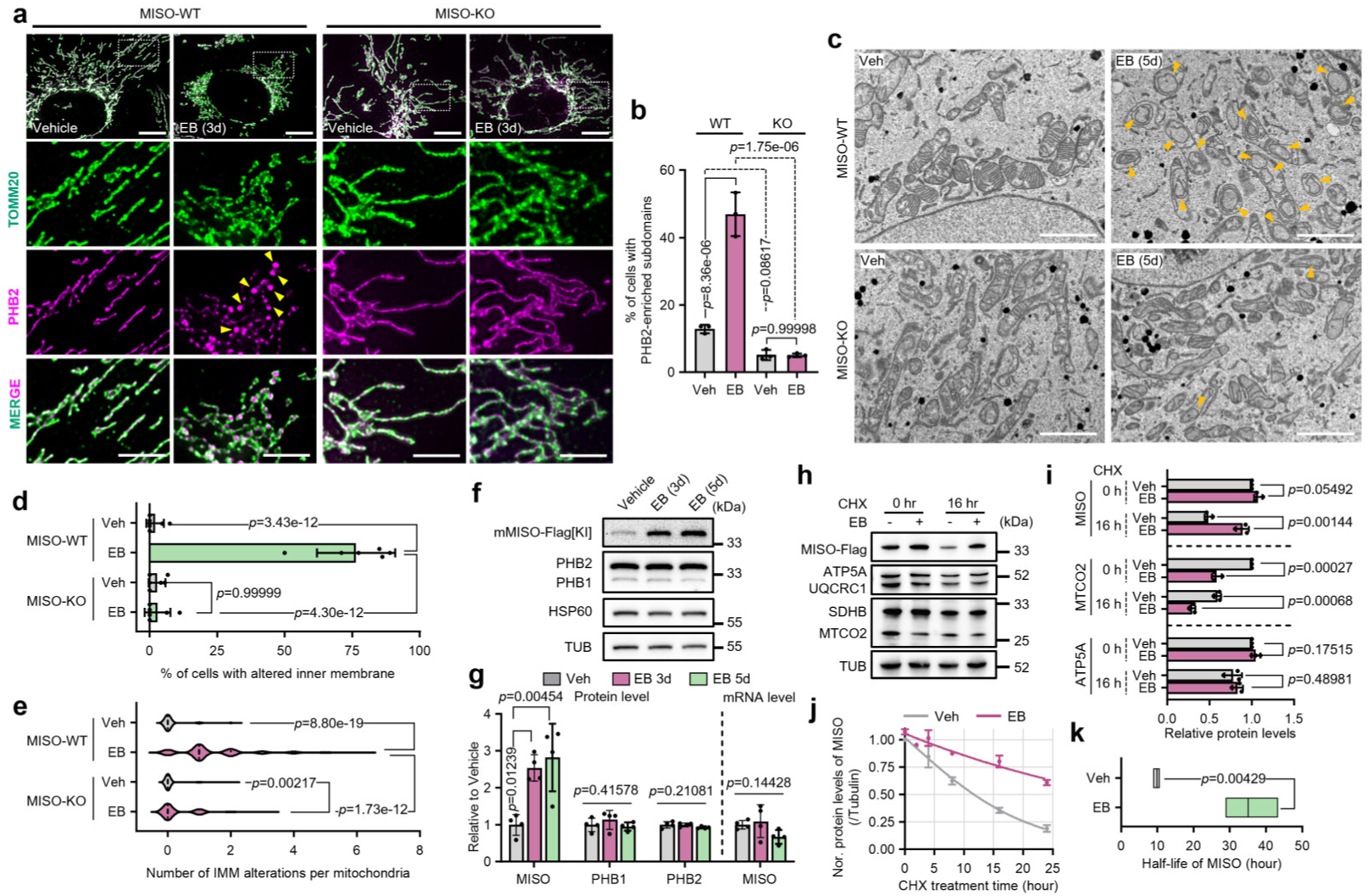
IMM stress leads to SMEM formation through the stabilization of the MISO protein. **a-b.** Representative images (**c**) and corresponding quantifications (**d**) of PHB2-enriched microdomains (yellow arrows) in WT and MISO-KO U2OS cells incubated in the presence or absence of EB. Scale bars, 10 μm and 5 μm. *n* = three experiments. **c-e.** Representative TEM images (**g**) and corresponding quantification (**d-e**) of WT and MISO-KO U2OS cells incubated in the presence or absence of EB. Yellow arrows indicate altered IMM subdomains. Scale bars: 2 μm. **f.** Immunoblots of indicated proteins from 3×Flag tag knock-in MEFs incubated in the presence or absence of EB. **g.** Quantification of indicated proteins and mRNA levels from (**o**). *n* = four experiments. **h.** Cycloheximide (CHX) chase analysis of MISO protein stability in U2OS cells stably expressing MISO and treated with or without EB. **i.** Quantification of indicated protein levels from (**e**). *n* = three experiments. **j.** Relative MISO protein levels at varying time points during cycloheximide (CHX) treatment in the presence or absence of EB. **k.** Half-life of MISO protein in the presence or absence of EB. All data are presented as mean ± SD. (**b**): two-way ANOVA with Tukey’s multiple comparisons test; (**d**) and (**e**): non-parametric Mann-Whitney test; (**g**): ordinary one-way ANOVA with Tukey’s multiple comparisons test; (**i**) and (**k**): two-tailed unpaired t test.

We next investigated how EB regulates MISO protein levels. MISO mRNA levels remained unchanged during EB treatment, suggesting that the observed increase in MISO is likely mediated through post-transcriptional mechanisms (**Fig. 5g**). Mitochondrial stress has been shown to stabilize certain proteins, such as mitochondrial-localized ATF5 transcription factors, that are normally degraded under basal conditions^29^. This prompted us to hypothesize that MISO might also be stabilized upon EB-induced stress. To isolate post-translational regulation from transcriptional effects, we overexpressed MISO in U2OS cells using a constitutive promoter and assessed protein stability by cycloheximide chase assay under both control and EB-treated conditions. Indeed, EB treatment significantly enhanced MISO protein stability (**Fig. 5h-j, Extended Data Fig. 25a-b**), extending its half-life from approximately 9 hours to over 30 hours (**Fig. 5k**). Notably, we also observed a marked reduction in Complex III and IV protein levels in EB-treated samples. Given that mtDNA encodes essential subunits of OXPHOS complexes I, III, IV, and V, we reasoned that MISO stabilization might be a general response to OXPHOS dysfunction. Taking advantage of the *Drosophila* MISO-KI line and previously validated OXPHOS RNAi lines (**Extended Data table 2**), we knocked down individual OXPHOS complexes and found that impairment of Complex III, IV, or V—but not Complex I or II—led to a significant increase in MISO levels (**Extended Data Fig. 26**). The fly adipose tissue was used in this study due to its highly organized and uniform cellular architecture, which facilitates the clear visualization of cellular changes. In contrast, acute inhibition of OXPHOS using rotenone (Complex I), sodium azide (NaN₃, Complex IV), oligomycin (Complex V), or the mitochondrial uncoupler CCCP resulted in reduced MISO levels (**Extended Data Fig. 27a-b**), and failed to induce SMEM formation (**Extended Data Fig. 27c-d**). These findings are consistent with previous reports showing that treatments with CCCP (which induces canonical mitophagy) or mito-paraquat (which elevates mitochondrial reactive oxygen species) fail to induce SMEM formation^10^, suggesting that SMEM biogenesis is a response to specific types of IMM damage. Furthermore, induction of mitophagy by high-dose CCCP in combination with Parkin overexpression resulted in comparable levels of mitochondrial clearance in both WT and MISO-KO cells (**Extended Data Fig. 27e-f**), supporting that MISO-dependent SMEM formation represents a distinct and parallel pathway to canonical Parkin-mediated mitophagy.

In addition to EB treatment, previous studies have shown that SMEM formation can be triggered by depletion of key cristae-organizing proteins, such as MIC60 and ATP5A^10^. Here, we demonstrated that MISO knock-out completely blocked the SMEM formation induced by MIC60 or ATP5A knockdown (**Extended Data Fig. 28a-c**). Moreover, similar to EB-induced stress, MISO protein—when expressed under a constitutive promoter—is stabilized upon MIC60 or ATP5A depletion (**Extended Data Fig. 28d-f**). Together, these findings establish MISO as a central organizer of SMEM that respond specifically to distinct forms of IMM stress, including mtDNA damage, OXPHOS dysfunction, and cristae disorganization, highlighting the critical role of MISO in orchestrating mitochondrial quality control pathways under diverse stress conditions.

### MISO drives SMEM formation by promoting intermembrane contacts

TEM experiment revealed that MISO overexpression led to a significant reduction in mitochondrial cristae, with the IMM forming densely packed, multilayered structures (**Fig. 6a-b, Extended Data Fig. 29a**). To further investigate this structural remodeling, we stained the IMM using 10-N-nonyl acridine orange (NAO), a fluorescent probe that specifically binds cardiolipin—a lipid enriched in the IMM—in live cells^30, 31^. The NAO signal was highly condensed within MISO-induced SMEM domains (marked by PHB1-mCherry) (**Fig. 6c-d, Extended Data Fig. 29b**), consistent with IMM aggregation. Given that canonical cristae architecture is maintained by the mitochondrial contact site and cristae organizing system (MICOS) and ATP synthase complexes^32, 33^, we examined the localization of key cristae-organizing proteins under MISO overexpression. Notably, both ATP5A and MIC60 were excluded from SMEM subdomains, despite their overall protein levels remaining unchanged (**Fig. 6e, Extended Data Fig. 29c-d**). Collectively, these results suggest that SMEM likely originate from localized alterations of IMM, driven by the exclusion of key cristae-organizing proteins.

**Fig. 6.**
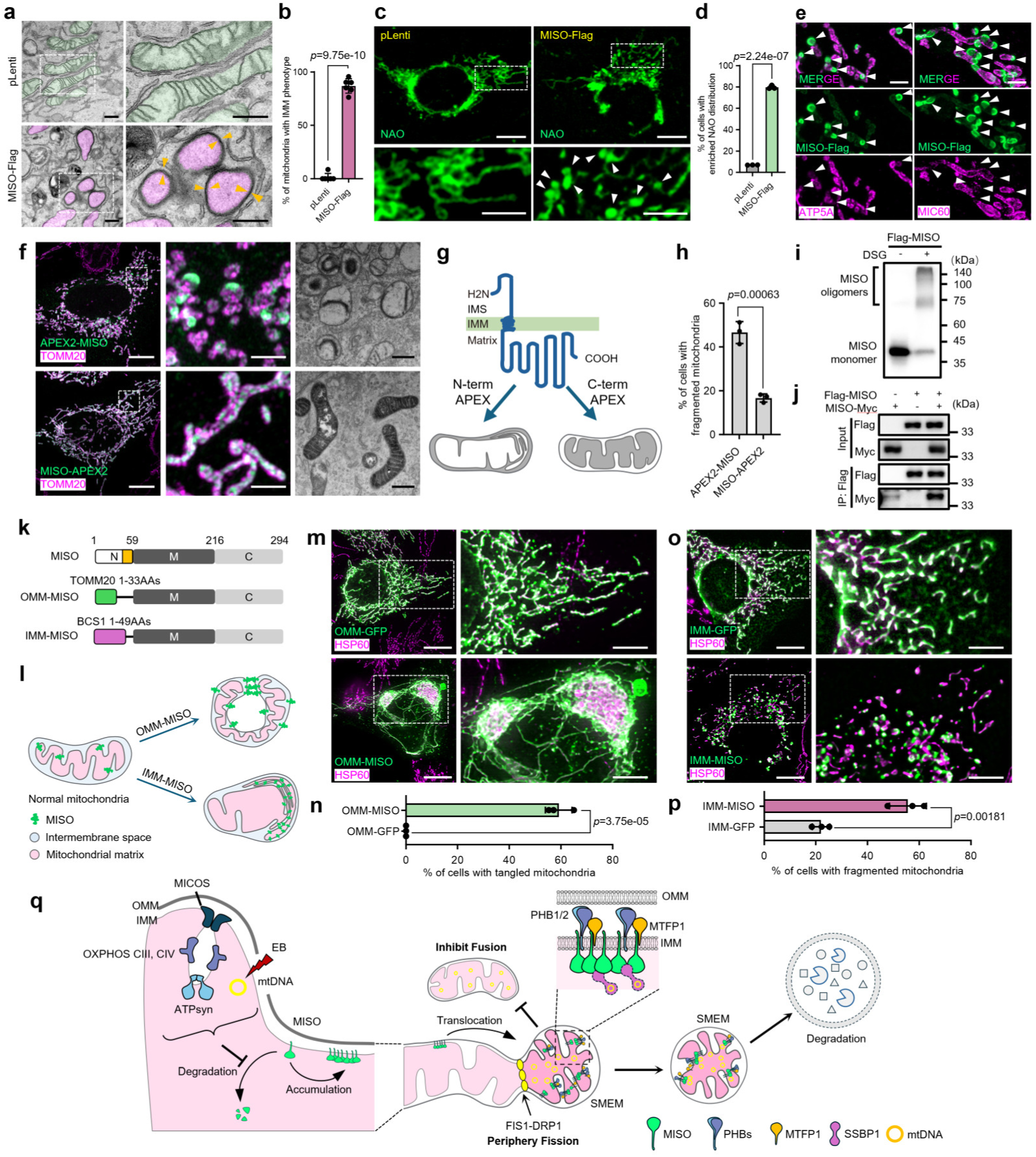
MISO promotes intermembrane interactions to drive SMEM formation. **a-b.** Representative TEM images (**a**) corresponding quantification (**b**) of U2OS cells expressing an empty vector (pLenti) or MISO-Flag. Yellow arrows indicate altered IMM. Scale bars: 500 nm. *n* = three experiments. **c-d.** Representative images (**c**) and corresponding quantification (**d**) of U2OS cells expressing an empty vector (pLenti) or MISO-Flag, incubated with nonyl acridine orange (NAO). Scale bars: main panels 10 μm, magnified insets 5 μm. *n* = three experiments. **e.** Representative images showing the absence of ATP5A and MIC60 within MISO-enriched subdomains. Scale bars: 1 μm. **f.** Representative fluorescence and electron microscopy images of U2OS cells expressing APEX2-MISO or MISO-APEX2. Scale bars: main panels 10 μm, magnified insets 2 μm for fluorescence images; 500 nm for TEM images. **g.** A Model illustrating the topology of MISO within the inner mitochondrial membrane (IMM). **h.** Quantification of mitochondrial morphology from (**e**). *n* = three experiments. **i.** Crosslinking analysis of MISO oligomerization by disuccinimidyl glutarate (DSG) in U2OS cells expressing MISO-Flag. **j.** Co-immunoprecipitation with anti-Flag to validate the self-interaction of MISO. **k.** Scheme of constructions of engineered MISO. MISO was segmented into N (1-59 AAs) and C (60-294 AAs) structural domain. The N terminus was replaced by a peptide localized to the outer mitochondrial membrane (TOMM20, 1-33 AAs) or inner mitochondrial membrane (BCS1, 1-49 AAs), which were fused to C regions. **l.** Model illustrating the dynamics of mitochondrial morphology by OMM-MISO or IMM-MISO. **m-n.** Representative images (**m**) and corresponding quantification (**n**) of mitochondrial morphology in U2OS cells expressing OMM-GFP or OMM-MISO. Scale bars: main panels 10 μm, magnified insets 5 μm. *n* = three experiments. **o-p.** Representative images (**o**) and corresponding quantification (**p**) of mitochondrial morphology in U2OS cells expressing IMM-GFP or IMM-MISO. Scale bars: main panels 10 μm, magnified insets 5 μm. *n* = three experiments. **q.** Proposed Model for the Formation and Function of MISO-Organized SMEM. Under normal conditions, MISO protein is rapidly degraded in mitochondria, maintaining low steady-state levels. However, upon mtDNA damage, OXPHOS complex impairment, or cristae structural abnormalities, MISO degradation is inhibited, leading to its accumulation. The accumulated MISO oligomerizes via its C-terminus within the mitochondrial matrix, promoting interactions between the inner mitochondrial membranes (IMM) to form specialized mitochondrial subdomains termed SMEMs. These SMEM restrict mitochondrial fusion potentially by excluding OPA1 from the subdomain. Additionally, SMEM recruit mtDNA, likely through MISO’s interaction with SSBP1, and facilitate their transport to mitochondrial termini. MISO also recruits other SMEM proteins, including PHB1/2 and MTFP1, into SMEM. These components promote peripheral mitochondrial fission and subsequently interact with LC3 for lysosomal degradation. Collectively, this model underscores the dual role of MISO-organized SMEM in coordinating mitochondrial dynamics and mtDNA quality control in response to specific IMM damage. All data are presented as mean ± SD. (**b**), (**d**), (**h**), (**n**) and (**p**): two-tailed unpaired t test.

To further elucidate how MISO drives structural remodeling of the IMM, we mapped its topology using TEM combined with ascorbate peroxidase (APEX) labeling^13^. APEX catalyzes the polymerization of diaminobenzidine (DAB) into a high electron-density precipitate that can be observed under TEM. By fusing APEX to either the N-or C-terminus of MISO, we found that the N-terminus of MISO resides in the mitochondrial intermembrane space, while the C-terminus is oriented toward the matrix (**Fig. 6f-g**). This topology is further supported by proteinase protection assays, which showed that a C-terminal Flag tag on MISO is shielded from proteases when the IMM remains intact (**Extended Data Fig. 11e**). Notably, when the relatively large APEX tag (28 kDa) was fused to the C-terminus, the resulting MISO-APEX construct failed to induce SMEM formation. This fusion of APEX also abolished MISO-induced mitochondrial fragmentation (**Fig. 6f-h**). In contrast, fusing small epitope tags such as Flag or Myc to the C-terminus did not impair MISO’s functionality, suggesting that the C-terminus may be critical for forming a densely packed complex, and steric hindrance from large fusions likely disrupts its proper assembly or activity.

A recent study has suggested that MISO belongs to the prohibitin-stomatin protein family^34^, which belongs to the evolutionarily conserved SPFH (Stomatin, Prohibitin, Flotillin, HflK/C) superfamily present in both prokaryotic and eukaryotic cells^35^. A hallmark of SPFH family proteins is their ability to self-oligomerize into large membrane-spanning complexes^36^. To investigate whether MISO exhibits similar oligomerization behavior, we performed disuccinimidyl glutarate (DSG) crosslinking experiments. Upon DSG treatment, distinct dimeric and trimeric MISO species were detected (**Fig. 6i**). This oligomerization is conserved across both MISO isoforms and remains unaffected by knockdown of major MISO-interacting proteins, including PHBs, MTFP1, and SSBP1 (**Extended Data Fig. 30a-b**), suggesting that oligomerization is likely to be an intrinsic property of MISO. Furthermore, co-immunoprecipitation assays confirmed that MISO interacts with itself (**Fig. 6j**). To identify the domain responsible for oligomerization, we tested MISO truncations—containing only the N-terminal, middle, or C-terminal regions—but none produced a clear crosslinked band (**Extended Data Fig. 30c**). The absence of detectable oligomers may reflect either a genuine failure of these truncations to oligomerize or, alternatively, a spatial separation between the interaction interface and the crosslinking-competent site.

An alternative approach to probe the biochemical functions of different MISO domains would be to reconstitute full-length MISO in liposomes and assess the behavior of truncated variants in vitro. However, attempts to obtain high-quality soluble full-length MISO protein from either prokaryotic or eukaryotic expression systems were unsuccessful (data not shown). As an alternative, we tested the functional capacity of MISO domains in a cellular context. We engineered an artificial construct, OMM-MISO, by replacing the N-terminal mitochondrial targeting sequence and transmembrane domain (residues 1-59) of MISO with the outer mitochondrial membrane (OMM) localization signal from TOMM20. This chimeric protein directs the matrix portion of MISO (residues 60-294, encompassing the middle and C-terminal domains) to the cytosolic side of the OMM (**Fig. 6k-l**). Strikingly, expression of OMM-MISO in cells induced extensive mitochondrial aggregation, leading to the formation of large, tangled mitochondrial networks in both U2OS and HEK293T cells (**Fig. 6m-n, Extended Data Fig. 30d-e**). This phenotype supports the hypothesis that the matrix-exposed region of MISO mediates inter-membrane IMM interactions. In addition, OMM-MISO with the middle region only (OMM-MISO-M, residues 60-216) failed to trigger mitochondrial aggregation (**Extended Data Fig. 30f-g),** again, suggesting that the C-terminus is essential for the IMM alternation property of MISO.

To further test whether the MISO’s matrix portion is sufficient for its biological function, we fused the MISO matrix portion (residues 60-294) with the IMM targeting sequence from BCS1^37^, a mitochondrial IMM protein with a topology analogous to MISO (**Fig. 6k-l**). Remarkably, IMM-MISO recapitulated key features of wild-type MISO: it induced the formation of SMEM-like subdomains and triggered mitochondrial fragmentation (**Fig. 6o-p, Extended Data Fig. 30h-j**), indicating that MISO’s function is mediated by its matrix-localized region. Interestingly, a truncated version of IMM-MISO containing only the middle domain (IMM-MISO-M, residues 60-216) failed to form SMEM (**Extended Data Fig. 30h**), yet still induced mitochondrial fragmentation like the IMM-MISO (residues 60-294). This uncoupling of phenotypes suggests that mitochondrial fragmentation and SMEM formation are separable functions of MISO, with the C-terminus playing a more critical role in driving structural reorganization of the IMM. Finally, constructs containing only the C-terminal domain (residues 216-294) fused to either OMM or IMM targeting signals (OMM-MISO-C and IMM-MISO-C) failed to localize properly to mitochondria (**Extended Data Fig. 30f, i**), which may be due to misfolding resulting from the absence of the middle domain. Collectively, these findings suggest that MISO may enhance the interactions between the inner mitochondrial membrane through its C-terminal domain, leading to cristae alternation and the formation of distinct SMEM subdomains.

## Discussion

Previous studies have established a strong link between mitochondrial dynamics and mtDNA homeostasis. Mutations in proteins involved in mitochondrial fusion and fission have been shown to alter the size and number of mitochondrial nucleoids^38^. Fission events have been associated with mtDNA replication and segregation, as the positioning and duplication of nucleoids are regulated by ER–mitochondria contact sites that mediate mitochondrial fission^23, 24^. Mitochondrial fusion also contributes to mtDNA maintenance: mutations in fusion machinery consistently result in reduced mtDNA copy numbers^38^. Conversely, inhibition of fission has been shown to increase mtDNA levels and promote the retention of mutant mtDNA^38, 39^. Several molecules have been proposed to link mitochondrial morphology with mtDNA nucleoid organization, including cardiolipin, which binds nucleoids and regulates mitochondrial IMM fusion^40, 41^, and the MICOS complex, which regulates both mitochondrial dynamics and mtDNA organization^42^. Despite these progresses, the exact molecular and structural mechanisms that integrate mitochondrial dynamics with mtDNA maintenance across different physiological and pathological contexts remain poorly understood.

Our study reveals a novel molecular link between mitochondrial dynamics and mtDNA maintenance by characterizing the MISO’s function in orchestrating mitochondrial subdomains, SMEM, particularly under IMM stress conditions. Mechanistically, we identify MISO as a short-lived IMM protein that contributes to basal mitochondrial morphology regulation. Specific mitochondrial stressors—including EB-induced mtDNA damage, disruption of the cristae-organizing complex MICOS, and mutations in OXPHOS complexes III, IV, and V (ATP synthase)—inhibit MISO degradation. This leads to MISO accumulation in the IMM, driving robust SMEM formation. MISO alters IMM architecture by displacing key cristae regulators, including MICOS and ATP synthase complexes, a process probably mediated by its self-polymerization and dependent on its C-terminal domain. Mitochondrial nucleoids are recruited into SMEM, likely through MISO-SSBP1 interactions. Given SMEM’s preferential localization at mitochondrial termini, their formation facilitates the peripheral redistribution of nucleoids. Furthermore, SMEM recruit the downstream effector MTFP1 via MISO binding, promoting peripheral mitochondrial fission through the FIS1-DRP1 pathway. Once detached from the mitochondrial network, SMEM are subsequently degraded via lysosomal mechanisms. Collectively, our findings establish the MISO-organized SMEM as stress-responsive signaling hubs that spatially compartmentalize regulatory molecules to coordinate mitochondrial morphology and mtDNA homeostasis (**Fig. 6q**).

Despite the findings presented above, several compelling questions regarding MISO-organized SMEMs remain outside the scope of this study, yet clearly warrant future investigation. First, the unique biochemical properties of MISO merit further exploration. Unlike typical mitochondrial proteins, which possess an N-terminal mitochondrial targeting sequence (MTS), MISO appears to require the coordinated action of multiple domains—including both its transmembrane and central regions—for proper subcellular targeting (**Extended Data Fig. 2**). Moreover, this targeting process is further modulated by the presence of its C-terminus (**Extended Data Fig. 30**). These observations underscore the complexity of MISO’s targeting mechanism and highlight the need for more extensive investigation to elucidate its biochemical behavior, which will be essential for designing mutants to dissect the functional roles of its various domains. Meanwhile, it remains unclear how MISO causes the reorganization of IMM. A recent study employed cryo-electron tomography to reveal the in situ supramolecular architecture of PHB complexes^18^. Examining the structure of MISO-containing SMEMs using similar techniques will provide detailed insights into how MISO assembles and organizes this specialized membrane domain. Meanwhile, it is important to note that MISO belongs to the SPFH protein family, which is well known for organizing cholesterol-rich lipid rafts on the cell membrane^36^. The mitochondrial IMM has traditionally been considered devoid of lipid raft-like domains due to its low cholesterol content. However, our findings demonstrate that MISO promotes the formation of SMEM subdomains highly enriched in cardiolipin (**Fig. 6c, Extended Data Fig. 29b**). This raises an intriguing possibility: different SPFH family proteins may organize distinct types of “raft-like” microdomains by preferentially interacting with specific lipids. Testing this hypothesis may broaden our understanding of functional compartmentalization across diverse cellular membranes.

Another compelling question is how specific IMM stressors stabilize the MISO protein. Mitochondrial unfolded protein response (UPR^mt^), caused by loss of membrane potential or increase of mitochondrial ROS, has been shown to stabilize ATF5 due to impaired import system^43^. However, this mechanism appears not to apply to MISO, as both this study and previous literature have shown that SMEM formation is not induced by classical stressors such as CCCP, oligomycin, or mito-paraquat^10^—agents widely used to trigger canonical UPR^mt^. This result indicates MISO may be regulated by a distinct protein quality control mechanism, different from classic UPR^mt^. Therefore, elucidating how MISO is degraded under basal conditions and how this degradation is inhibited by certain IMM stressors could provide a new avenue for understanding mitochondrial protein homeostasis.

Finally, MISO knockout in both Drosophila and mice is viable and fertile, suggesting that MISO is not essential for normal development. However, its evolutionarily conserved role in responding to mitochondrial stress in both flies and mammals indicates that it may play important physiological roles under pathological conditions. Indeed, recent high-throughput studies have linked MISO to various diseases such as cancers^44^ and brain-related disorders, including common epilepsies and alcohol dependence^45, 46^ —findings that are consistent with its comparatively high expression in the mouse central nervous system (CNS). Therefore, further investigation under pathological or sensitized conditions may provide deeper insights into the functional significance of MISO-organized SMEM.

## Author Contributions

Conceptualization, experiments design by L.H., Y.Z., and Q.Z.; experiments performance, data collection and analysis by Y.Z., Y.X., X.W., Y.X., S.W., J.L., X.G., Q.Z., and L.H. All authors have read and agreed to the published version of the manuscript.

## Data Availability Statement

All data generated or analyzed in this study are included in the main text or the Supplementary materials. The raw data generated in this study are provided in the Source Data file in this paper. The protein mass spectrometry data has been deposited on the ProteomeXchange PRIDE database with the identifier: PXD057461.

## Acknowledgments

funding for this work is from the National Natural Science Foundation of China (No. 32470754, 32070750) and Opening Foundation of National Engineering Research Center of Genetic Medicine, China (NERCGM-OF-20250201) to L.H.; National Natural Science Foundation of China (No. 32430049, 82370596), National Key R&D Program of China (No. 2021YFA08049000), The Guangdong Basic and Applied Basic Research Foundation (No. 2023B1515120089) to Q. Z.; National Natural Science Foundation of China (No. 82370576, 32170772) to J. L.

We thank Dr. Norbert Perrimon and Dr. Richard Binari at Harvard Medical School for support of the initial genetic screen in *Drosophila*. We thank the Cryo-EM Center in University of Science and Technology of China and the Core facility of Harvard Medical School for preparation of ultrathin sections and TEM operation. JL is supported by the Youth S&T Talent Support Programme of Guangdong Provincial Association for Science and Technology. JL and QZ are supported by the K.C. Wong Education Foundation.

