## Supplementary Information for "MISO Regulates Mitochondrial Dynamics and mtDNA Homeostasis by Establishing Membrane Subdomains"

### Materials and Methods

#### Antibodies

The following primary antibodies were used for immunoblotting or immunostaining: mouse anti-Actin (Proteintech, 66009-1-Ig), rabbit anti-ATP5A (Proteintech, 14676-1-AP), rabbit anti-COX5B (Proteintech, 11418-2-AP), rabbit anti-Cyto c (Proteintech, 10993-1-AP), rabbit anti-FASTKD2 (Proteintech, 17464-1-AP), mouse anti-Flag (Proteintech, 66008-4-Ig), rabbit anti-FIS1 (Proteintech, 10956-1-AP), rabbit anti-GADPH (Proteintech, 60004-1-Ig), rabbit anti-GM130 (Proteintech, 11308-1-AP), rabbit anti-HSP60 (Proteintech, 15282-1-AP), rabbit anti-MFF (Proteintech, 17090-1-AP), rabbit anti-MFN2 (Proteintech, 12186-1-AP), rabbit anti-MIC60 (Proteintech, 10179-1-AP), rabbit anti-MID49 (Proteintech, 16413-1-AP), rabbit anti-MID51 (Proteintech, 20164-1-AP), rabbit anti-MTCO2 (Proteintech, 55070-1-AP), rabbit anti-MTFP1 (Proteintech, 14257-1-AP), rabbit anti-NDUFS3 (Proteintech, 15066-1-AP), rabbit anti-OXPHOS Cocktail (Proteintech, PK30006), rabbit anti-SDHA (Proteintech, 14865-1-AP), rabbit anti-SSBP1 (Proteintech, 12212-1-AP), rabbit anti-STOML2 (Proteintech, 10348-1-AP), rabbit anti-TFAM (Proteintech, 22586-1-AP), rabbit anti-TIMM23 (Proteintech, 11123-1-AP), rabbit anti-TOMM20 (Proteintech, 11802-1-AP), rabbit anti-TOMM40 (Proteintech, 18409-1-AP), rabbit anti-TOMM70 (Proteintech, 14528-1-AP), rabbit anti-Tubulin (Proteintech, 11224-1-AP), rabbit anti-UQCRC2 (Proteintech, 14742-1-AP), rabbit anti-DRP1 (Cell Signaling Technology, 8570), rabbit anti-HA (Cell Signaling Technology, 3724), rabbit anti-LAMP1 (Cell Signaling Technology, 9091), rabbit anti-LC3B (Cell Signaling Technology, 2775), rabbit anti-MFN1 (Cell Signaling Technology, 14739), rabbit anti-OPA1 (Cell Signaling Technology, 80471), rabbit anti-PHB2 (Cell Signaling Technology, 14085), mouse anti-ATP5A (Abcam, ab14748), mouse anti-MTCO1 (Abcam, ab14705), rabbit anti-PHB (Abcam, ab28172), rabbit anti-SDHB (Abcam, ab14714), mouse anti-Delta (DSHB, C594.9B), mouse anti-LacZ (DSHB, 40-1a), mouse anti-Myc (Santa Cruz, sc-40), mouse anti-TOMM20 (Santa Cruz, sc-17764), rabbit anti-Myc (Sigma, C3956), mouse anti-Flag (Sigma, F3165), chicken anti-GFP (Aves Labs, GFP-1020), mouse anti-HA (Biolegend, 902301), rabbit anti-PMP70 (Invitrogen, PA1-650), mouse anti-DNA (Progen, 690014) and rabbit anti-C3orf33(MISO) (generated by Dia-An Biotechnology, Wuhan, China). We also used the following secondary antibodies: anti-mouse IgG (Proteintech, SA00001-1) or anti-rabbit IgG (Proteintech, SA00001-2) conjugated to HRP for immunoblotting, and anti-mouse AlexaFluor-488 (Invitrogen, A-11001), anti-mouse AlexaFluor-555 (Invitrogen, A-21422), anti-rabbit AlexaFluor 555 (Invitrogen, A-21428), anti-rabbit AlexaFluor-647 (Invitrogen, A-21245), anti-chicken AlexaFluor-488 (Invitrogen, A-11039), anti-Mouse IgG2b Cross-Adsorbed AlexaFluor-488 (Invitrogen, A-21141) or anti-Mouse IgM (Heavy chain) Cross-Adsorbed AlexaFluor-546 (Invitrogen, A-21045) for immunostaining.

**DNA Constructs**

The full-length cDNA of the human gene C3orf33 (MISO) was synthesized by Sangon Biotech and sub-cloned into the pcDNA3.1-3×Flag vector at the C-terminus using the Seamless Cloning Kit (Beyotime, D7010). Additionally, the pcDNA3.1-MISO-3×Myc plasmid was also constructed. For stably expressing in mammalian cells, MISO-3×Flag or MISO-3×Myc fragment was amplified by PCR and subsequently replaced with EcoRI-SalI fragment of the virus vector pLenti-CMV-GFP-Puro (658-5) (Addgene, 17448) To generate pLenti-TetOne-Puro-MISO-3×Flag, MISO-3×Flag fragment was amplified from pcDNA3.1- MISO-3×Flag and inserted into pLenti-TetOne-Puro via BamHI restriction site. To generate pCDNA3.1-PHB1/PHB2-3×Myc plasmid, PHB1 or PHB2 fragment was amplified from HEK293T cDNA and subsequently cloned into pcDNA3.1 with a C-terminal 3×Myc. The plasmid pLenti-TetOne-Blast-PHB1/PHB2-3×Myc-mNeonGreen was constructed with PHB1/PHB2-3×Myc fragment, mNeonGreen fragment and pLenti-TetOne-Blast vector (cut with BamHI restriction site) The cDNAs encoding SLC25A4, SLC25A5, SLC25A6, ATAD3A, ATAD3B, VDAC1, VDAC2, MFN1 and MFN2, as well as the plasmid mCherry-Parkin, were obtained from MiaoLing Bio (Wuhan, China). Then the SLC25A4, SLC25A5, SLC25A6, ATAD3A or ATAD3B fragment was cloned into pcDNA3.1 with a C-terminal 3×Myc and VDAC1 or VDAC2 fragment was inserted into pcDNA3.1 with a N-terminal 3×HA. The MFN1 and MFN2 was cloned into pLVX-mCherry-N1 vector at the EcoRI and BamHI restriction sites. The first 69 amino acids of ATP9 from Neurospora crassa were inserted into the PCDH-CMV-EF1-Hygro vector at the Xbal and BamHI restriction sites, fused with a C-terminal TagRFP or EGFP fragment, to generate a mitochondrial matrix-targeted protein. For the plasmids OMM-MISO or IMM-MISO, the first 58 amino acids of MISO were replaced with either a mitochondrial outer membrane targeting sequence (the first 33 amino acids of TOMM20) or a mitochondrial inner membrane targeting sequence (the first 49 amino acids of BCS1). These modified sequences were then sub-cloned into the pCDNA3.1 vector along with a C-terminal 3×Flag tag. All restriction enzymes used here were purchased from TAKARA. For RNAi in vitro, shRNA sequences were cloned into the pLKO.1 vector using AgeI and EcoRI restriction sites. The targeted sequences were as follows: Non-Targeting (NT) shRNA: CCTAAGGTTAAGTCGCCCTCG; shMISO-1: GGACGATTACGCCGAATAACT; shMISO-2: GGAAAGACAACATGAACAACT; shDRP1: GCTACTTTACTCCAACTTATT; shFIS1: CAAGAGCACGCAGTTTGAGTA; shMFN1: GCTCAAAGTTGTAAATGCTTT; shMFN2: GCAGGTTTACTGCGAGGAAAT; shOPA1: GACTACTGCTTGCTGCAAAGG; shMTFP1-1: GCCATTGACAAAGGCAAGAAG; shMTFP1-2: TGGCGGATGCCATTGACAAAG; shMIC60: GCCTGTACCAATACTTCCTTT; shATP5A: GCCAAGATGAACGATTCCTTT; shSSBP1: CATGGCACAGAATATCAGTAT; shPHB1: GAGTTCACAGAAGCGGTGGAA; shPHB2: AAGAACCCTGGCTACATCAAA. All plasmids were verified by Sanger sequencing.

***Drosophila* and Mice**

The fly strains *UAS-Luc RNAi* (BL35788)*, UAS-CG30159 RNAi* (BL61888), *UAS-myr::tdTomato* (BL32221) *UAS-NP15.6 RNAi* (BL36672), *UAS-ND-49 RNAi* (BL28573), *UAS-ND-23 RNAi* (BL30487), *UAS-SdhC RNAi* (BL53281), *UAS-UQCR-Q RNAi* (BL51357), *UAS-COX6C RNAi* (BL33878) and *UAS-ATPsynβ RNAi-2* (BL28062) were obtained from the Bloomington Drosophila Stock Center. *UAS-SdhC RNAi (NIG7361R-1); UAS-SdhC RNAi* (NIG14724R-3) and *UAS-ATPsynβ RNAi-1* (NIG11154R-1) were obtained from the Fly Stocks of National Institute of Genetics. *UAS-CG30159-HA* (F002359) was received from the FlyORF Stock Center., *esg-GAL4*, *Delta-GAL4*, *Delta-LacZ*, *Su(H)Gbe-lacZ*, *UAS-nlsGFP* and *hsFlp; AyGal4, UAS-GFP* were from lab stocks. The CG30159 deletion mutants, *CG30159-GAL4* and *CG30159-3×HA*[KI] transgenic strains were established by CRISPR-Cas9 mediated genome editing. In brief, sgRNAs (TTCCAGTTATCCAATGGAGG and GGAGCGGGATACGCGCGGCA for Gal4 knock-in; ATAGCATCGAGGCAGGAACG, CCTGTAAATAGCATCGAGGC and TAGCCCTGTAAATAGCATCG for 3×HA knock-in) were cloned into pCFD3 plasmid. For knock-in stocks, two homology arms amplified from fly genome were cloned into upstream and downstream of GAL4-SV40 or 3×Flag in pENTRY vector as donor construct. For CRISPR-Cas9 mediated homologous recombination, gRNA in pCFD3 (0.2 μg/μl) and donor DNA (0.5 μg/μl), were co-injected into the embryos of nos-Cas9/attP2 flies (BL78782) by UniHuaii Corporation (Guangzhou, China). Mutants or knock-in flies were verified by genomic PCR and sequencing. Fly strains were maintained with standard cornmeal-yeast-agar medium and were kept at 25 °C. For overexpression or RNAi knockdown, flies were reared at 32 °C after larval hatching to boost efficiency.

All mouse experiments were conducted in accordance with the China Animal Welfare Legislation guidelines and approved by the Animal Research Ethics Committee of the University of Science and Technology of China (USTCACUC25070120006) and the Animal Ethics Committee of Jinan University (IACUC2024011009). *mMISO*$^{flox/flox}$ mice (C57BL/6 background) were generated by GemPharmatech (Nanjing, China) using CRISPR/Cas9 technology. MISO knock-in mice expressing a triple Flag tag at the C-terminus of MISO (C57BL/6 background) were generated via CRISPR/Cas9 technology. Two sgRNAs (GTTATCACGACAGCCGGCGC and TTATCACGACAGCCGGCGCA) were used to target the insertion site for precise tagging. To generate mice with MISO deletion, *mMISO*$^{flox/flox}$ mice were crossed with *ZP3-Cre* transgenic mice. Offsprings with successful MISO deletion were verified by genomic PCR and sequencing. Genotyping was performed using the following primers: mF1: GCTATTGTAGCGCCACGTTAGAGC; mR1: CTGTATAGAGCTGGCTGTCCTGGTAC; mF2: CATCGCATTGTCTGAGTAGGTG; mR2: CTGGTCTCCTTTGGCTCCTTTC. All mice were maintained on a 12-hour light-dark cycle at a room temperature of 22 °C ± 2 °C, humidity of 50% ± 5%, with free access to water and standard rodent diet.

**Cell Culture and Treatments**

U2OS, HeLa, HEK293T and Huh7 cells were obtained from the Shanghai Cell Bank Type Culture Collection Committee (Shanghai, China). HCT116 and PLC/PRF/5 cells were provided by HuaFeng Zhang Lab (USTC). To generate MISO-knockout (MISO-KO) cell lines, U2OS cells were transfected with pSpCas9(BB)-2A-Puro (Addgene, 48139) containing specific sgRNAs (GAGGTGGGAGGTGCTTAATT and TTGATGTCTATTGATTTACA) targeting a common exon shared by both isoforms of MISO. Positive clones were selected using 1 μg/ml puromycin. Successful MISO knockout was confirmed by genomic PCR and Sanger sequencing. Genotyping was performed using the following four primers: F1: CCTGGTTGGTTAAGGTTCTAACT; R1: GCTTAGGTTCTAGACCATAATAGG; F2: GATTACGCCGAATAACTGAGAATG; R2: CTAAACCATTCTCAGTTATTCGGC. Cells were cultured at 37 °C in Dulbecco's Modified Eagle's Medium (DMEM) (Thermo-Fisher, C11965500BT) containing 10% fetal bovine serum (FBS) (Lonsera, S711-001S), supplemented with penicillin/streptomycin (Gbico, 15140122) and GlutaMAX Supplement (Gbico, 35050061) in 5% $CO_2$. To detach the cell monolayer, 0.25% Trypsin-EDTA with phenol red (Thermo Scientific, 25200072) was added for 1 min. The cells were then resuspended, and a dilution was made for maintenance. Plasmids were transfected into cells using the Lipo6000™ Transfection Reagent (Beyotime, C0526).

S2 cells were gifts from Linfeng Sun Lab (USTC) and cultured at 25 °C in Schneider's Medium (Sigma, S9895) with 10% FBS and 1% penicillin/streptomycin. Transfections were performed using the Effectene Transfection Kit (Qiagen, 301425). After transfection, the cultured cells were incubated for 1-2 days and seeded on 100 μg/ml Concanavalin A coated glasses before cell fixation.

To induce mitochondrial fragmentation, U2OS cells were incubated with 10 μM Carbonyl cyanide 3-chlorophenylhydrazone (CCCP) (SigmaAldrich, C2759) at 37 °C for 2 hours. To induce the formation of PHB2-enriched subdomains, ethidium bromide (EB) was applied at a final concentration of 50 ng/ml for 3 or 5 days. $H_2O$ served as the control vehicle. To measure the degradation rate of MISO, U2OS cells were treated with 50 μg/mL cycloheximide (CHX) for the indicated time points before harvesting. To investigate factors that regulate MISO protein levels or promote the formation of PHB2-enriched subdomains, U2OS cells were treated for 50 ng/mL EB, 1 μM Rotenone, 5 mM $NaN_3$, 2 μM Oligomycin or 2 μM CCCP for 24h before biochemical and imaging analyses. To induce mitophagy, U2OS cells were incubated with 10 μM CCCP for 12h.

**Primary Cell Isolation**

Mouse hepatocytes were isolated from male mice of the indicated genotypes at 8-10 weeks of age using previously published protocol[1, 2]. Briefly, the liver was perfused with 15 ml of perfusion buffer (Hanks' Balanced Salt Solution (HBSS) without $Ca^{2+}$ or $Mg^{2+}$, supplemented with 0.5 mM EGTA and 25 mM HEPES, pH 7.4) via the inferior vena cava. This was followed by perfusion with 15 ml of digestion buffer (HBSS containing $Ca^{2+}$ and $Mg^{2+}$, 25 mM HEPES, and 1 mg/ml Collagenase Type IV (Sigma, C5138), pH 7.4). After perfusion, the liver was excised, and hepatocytes were released into Williams E medium supplemented with 5% fetal bovine serum (FBS) and 1% penicillin/streptomycin. The cell suspension was then filtered through a 70 μm cell strainer. Following centrifugation, hepatocytes were purified using 50% Percoll (Yeasen, 40501ES60) to remove dead cells. The viability of the isolated hepatocytes, as determined by Trypan blue staining, was approximately 90%. Finally, the hepatocytes were plated onto collagen-coated wells in Williams E medium supplemented with 1% GlutaMAX, 1% penicillin-streptomycin, 0.1 nM insulin (Beyotime, P3376), 0.1 μM dexamethasone (Beyotime, ST1254), and 2 mM sodium pyruvate (Macklin, P6033).

Mouse embryonic fibroblasts (MEFs) were isolated from E13.5 embryos of mice with the indicated genotypes according to published protocol [3]. Each embryo was decapitated, and the internal organs were removed. The remaining tissue was minced and dissociated in 3 ml of 0.25% trypsin-EDTA at 37 °C for 30 minutes. The dissociated cells were then filtered through a 70 μm cell strainer, collected by centrifugation, and plated in DMEM supplemented with 10% FBS, 1% GlutaMAX, and 1% penicillin-streptomycin.

**Stable Cell Lines Construction**

Lentivirus packaging was conducted in HEK293T cells. One day before transfection, the cells were seeded on 6-well cell culture plates at a density of 60-70% confluence. The following day, a mixture containing 1 μg of lentiviral transfer plasmid encoding the gene of interest (with the target gene cloned into pLenti CMV GFP), 750 ng of psPAX2, and 250 ng of pMD2.G were co-transfected into the HEK293T cells using Lipo6000™ Transfection Reagent. 4-6 hours post-transfection, the cell culture medium was replaced with fresh medium. 48 hours after transfection, the supernatant was collected and filtered through a 0.45 μm pore size cellulose acetate filter. The lentiviruses were used immediately or stored at -80 °C for future use. To transduce target cells with lentivirus, an appropriate volume of lentivirus, along with polybrene, was added to the cell culture medium, and the cells were incubated for 12-16 hours. Subsequently, the medium was changed to fresh culture media supplemented with puromycin, blasticidin, or hygromycin to select for successfully infected cells.

**Live imaging**

For analysis of mitochondrial dynamics, U2OS cells expressing MISO-Flag, PHB2-mNeonGreen (PHB2-mNG), and mitochondrial matrix-targeted TagRFP (mito-RFP) were replated on glass-bottomed dishes. Live imaging was conducted using $CO_2$ Independent Medium (Gibco, 18045088) supplemented with 10% FBS and 100 U/ml Penicillin/Streptomycin.

For measurement of mitochondrial membrane potential, U2OS cells expressing MISO-Flag and PHB2-mNG were incubated with 100 nM tetramethylrhodamine (TMRM; Invitrogen, T668) at 37°C for 30 minutes.

For detection of oxidative stress, U2OS cells expressing MISO-Flag and PHB2-mNG were incubated with 1 μM MitoSOX Red (Invitrogen, M36007) at 37 °C for 30 minutes.

For the visualization of cardiolipin, U2OS cells expressing MISO-Flag and PHB1-mCherry were incubated with 1 nM Nonyl Acridine Orange (NAO) (Invitrogen, A1372) at 37 °C for 14 hours.

For tracking mtDNA, U2OS cells expressing MISO-Flag and PHB1-mCherry were incubated with PicoGreen (Invitrogen, P7581) diluted 1:500 at 37 °C for 30 minutes. Following incubation, cells were washed three times with pre-warmed PBS and then imaged. All the images were acquired with a Leica THUNDER imager equipped with a ×63 oil-immersion lens. Time-lapse images were acquired every 5 seconds.

**Transmission Electron Microscopy**

For transmission electron microscopy (TEM), cells grown on glass coverslips were fixed 3% glutaraldehyde (Ted Pella, EM grade, 18427) and 2% paraformaldehyde solution (Ted Pella, 18505-100) at room temperature for 2 hours. After fixation, cells were rinsed three times with 0.1 M sodium cacodylate buffer and then stained with sodium cacodylate buffer containing 1.5% potassium ferrocyanide and 1% osmium tetroxide (Electron Microscopy Sciences, 19190). After washing, the cells were incubated with 1% osmium tetroxide for another 40 minutes. Incubate the cells with 2% uranyl acetate solution at 4 °C in the dark for 1 hour, rinsed with ice-cold distilled water, and dehydrated in a graded ethanol series (30%-100%) for 15 minutes at each step. The sample was then mixed 1:1 with SPI-Pon 812 (SPI Supplies, 1260804) and anhydrous ethanol for 1 hour twice. The mixture was incubated overnight in 1:1 resin and then transferred to fresh resin, followed by polymerization at 60 °C for 48 hours. The sample was cut with a diamond knife into 70 nm sections and imaged on an FEI-Tecnai T12 120KV TEM instrument (TECNAI T12 120KV FEI, USA) at the Cryo-EM Center in USTC.

**APEX2 Staining for TEM**

Staining for APEX2 was performed as previously published protocol [4], with minor modifications. Briefly, cells were transiently transfected with either MISO-APEX2 or APEX2-MISO constructs for 20 hours. They were then fixed in 2% glutaraldehyde (EM grade; Sigma-Aldrich) in 0.1 M sodium cacodylate buffer at 4 °C for 16 hours. Post-fixation, the cells were rinsed three times with 0.1 M sodium cacodylate buffer and quenched with 20 mM glycine for 15 minutes to neutralize residual aldehydes. After additional washes, the cells were incubated on ice in a staining solution consisting of 0.1 M sodium cacodylate buffer containing 0.5 mg/ml DAB (Sigma, D12384), 10 mM hydrogen peroxide ($H_2O_2$), and 2 mM calcium chloride ($CaCl_2$). This incubation lasted between 30 to 45 minutes. Following this incubation, specimens were stained with osmium tetroxide and uranyl acetate, dehydrated through a graded series of ethanol solutions followed by acetone, infiltrated with Epon resin, embedded, sectioned, and analyzed by TEM as previously detailed.

**Immunoblotting and Chemical Cross-linking**

Cell or tissue samples were lysed in RIPA buffer (50 mM Tris-HCl, pH 8.0, 150 mM NaCl, 5 mM EDTA, 0.1% SDS, 1% NP-40) supplemented with a protease inhibitor cocktail (Beyotime, P1005) on ice for 30 minutes. The lysates were then centrifuged at 12,000×g for 20 minutes at 4 °C to pellet debris. The supernatant was collected, and the protein concentration of each sample was determined using BCA assay kit (Biosharp, BL521A). Next, 2×SDS loading buffer (125 mM Tris-HCl, pH 6.8, 4% SDS, 10% β-mercaptoethanol, 20% glycerol, 0.004% bromophenol blue) was added to the samples, which were then mixed thoroughly and boiled at 100 °C for 5 minutes. The proteins were separated by SDS-PAGE and transferred onto NC membranes. The membranes were blocked with 5% non-fat powdered milk (Sangon, A600669) in TBST (20 mM Tris-HCl, 150 mM NaCl, 0.05% Tween-20) at room temperature for 30 minutes, followed by three 10-minute washes with TBST. The membranes were then incubated overnight at 4 °C with the indicated primary antibodies. After washing with TBST, the membranes were incubated with HRP-conjugated secondary antibodies at room temperature for 1 hour. Finally, after three additional washes with TBST, the protein bands were visualized using SuperSignal West Pico PLUS Chemiluminescent Substrate (Thermo Scientific, 34580).
To analyze MISO oligomerization, U2OS cells expressing MISO-Flag were treated with 1 mM disuccinimidyl glutarate (DSG; Invitrogen, 20593) at 4 °C for 2 hours. The cells were then washed with PBS containing 20 mM glycine to quench the reaction. Subsequently, the samples were collected for immunoblot analysis.

**Immunofluorescence**

Drosophila tissues were fixed in 4% paraformaldehyde (PFA) at room temperature for 30 minutes. Mammalian cells were fixed with 4% PFA at 37 °C for 15 minutes, while S2 cells were fixed at room temperature for the same duration. Fixed samples were washed three times with PBST (PBS containing 0.05% Triton X-100), each washing lasting 10 minutes.

Subsequently, the samples were incubated in blocking solution (PBSTB: PBS containing 0.05% Triton X-100 and 2% BSA) at room temperature for 1 hour. After washing samples with PBST three times, the samples were then incubated with primary antibody at 4 °C overnight. Following an additional washing step in PBST, samples were incubated with matching secondary antibodies at room temperature for 2 hours. The cell nuclei were stained with DAPI. Finally, the samples were mounted with Antifade Mounting Medium (Beyotime, P0126) or ProLong Gold antifade reagent (Invitrogen, P36965) and imaged using a Leica DMi8 equipped with a THUNDER imager, Leica STELLARIS STED, or Zeiss LSM 980 with Airyscan 2.

To visualize replicating mtDNA, the BeyoClick™ EdU-555 labeling kit (Beyotime, C0075S) was used according to the manufacturer's instructions. Cells were incubated with 10 μM EdU at 37 °C for 4 hours to allow incorporation into newly synthesized mtDNA. Following the incubation, cells were fixed and processed for subsequent immunofluorescence analysis.

**Sample preparation of expansion microscopy**

Expansion microscopy experiment was carried out according to the published protocol[5]. Samples were immunostained as described above, but with all antibody concentrations doubled to enhance signal retention during expansion. Following staining, samples were anchored with 0.1 mg/ml methacryloyl-activated NHS ester (MA-NHS) in PBS for 4-5 hours at room temperature, then immersed in a pre-gel solution containing 4% *N, N*-dimethylacrylamide, 34% sodium acrylate, 10% acrylamide, 0.01% *N, N*-methylenebisacrylamide, and 1% sodium chloride in PBS, supplemented with 0.1% ammonium persulfate (APS) and 0.1% tetramethylethylenediamine (TEMED) to initiate polymerization. Gelation was carried out at 37 °C for 5-6 hours. After gel formation, the hydrogels were transferred to denaturation buffer (10% SDS, 8 M urea, 25 mM EDTA in 2× PBS, pH 7.4) and incubated at 80 °C for 6 hours. Subsequently, the gels were washed three times with PBST and mounted on glass-bottom dishes for imaging.

**Mitochondrial Isolation**

Cells were harvested after being washed twice with pre-chilled PBS. They were then resuspended in Lysis Buffer (Mitochondrial Extraction Kit, Solarbio, SM0020) and transferred to a glass homogenizer for grinding on ice, performing 40-50 cycles. Following this, the homogenate was centrifuged at 1000×g for 5 minutes. The supernatant was carefully transferred to a new centrifuge tube and subjected to another centrifugation at the same speed and duration. The resulting supernatant, designated as the whole cell lysate (WCL), was then transferred to a fresh tube. To isolate the cytoplasmic fraction (Cyto) and crude mitochondrial pellets, the WCL was centrifuged at 12,000×g for 10 minutes. The crude mitochondrial pellets were resuspended in Wash Buffer provided by the Mitochondrial Extraction Kit. To purify the mitochondria, the resuspended pellets underwent two rounds of centrifugation at 1000×g for 5 minutes each, followed by a final centrifugation at 12,000×g for 10 minutes. The purified mitochondrial pellets were collected and resuspended in Mito buffer (comprising 250 mM mannitol, 5 mM HEPES, pH 7.4, and 0.5 mM EGTA). All steps of mitochondrial isolation were performed at 4 °C to maintain mitochondrial integrity.

**Proteolysis of Mitochondria**

The phosphate swelling-shrinking assay was carried out as previously described[6]. Mitochondria were isolated as detailed in the mitochondrial isolation method, and the purified mitochondrial pellets were resuspended in swelling buffer (10 mM $KH_2PO_4$, pH 7.4) and incubated on ice for 20 minutes. An equal volume of shrinking buffer (10 mM $KH_2PO_4$, pH 7.4, 32% sucrose, 30% glycerol, 10 mM $MgCl_2$) was then added to the mitochondrial suspension to stabilize mitoplasts, followed by an additional 20-minute incubation. Mitochondria and mitoplasts were subsequently incubated in homogenization buffer (10 mM Tris-MOPS, 1 mM EGTA, 200 mM sucrose, pH 7.4) containing 0.2 mg/ml proteinase K, either with or without 1% NP-40, for 30 minutes. After incubation, protease inhibitor cocktail was added to terminate proteolytic activity. The samples were then mixed with 2× SDS loading buffer and boiled at 100 °C for 5 minutes prior to immunoblot analysis.

**Alkaline extraction of mitochondrial proteins**

Alkaline extraction was performed as described previously[7]. Briefly, purified mitochondria were resuspended in one of the following buffers: 0.1 M $Na_2CO_3$ (pH 11.0, 11.5, or 12.0) or PBS (pH 7.4). Samples were incubated on ice for 30 min with brief vortex mixing at 10-min intervals. Following incubation, samples were subjected to ultra-centrifugation at 51,000 rpm for 30 min at 4 ℃in a TLA 120 rotor (Beckman Coulter). The resulting membrane pellets were solubilized directly in Laemmli sample buffer (2% SDS, 10% glycerol, 60 mM Tris-HCl pH 6.8, 0.005% bromophenol blue). Supernatants were subjected to trichloroacetic acid (TCA) precipitation and dissolved in Laemmli loading buffer. Equal proportions of the initial total mitochondria (T), mitochondrial membrane pellet (P), and soluble supernatant (S) fractions were analyzed by SDS-PAGE and immunoblotting to assess protein distribution.

**Blue-native PAGE and in-gel activity assays**

Blue native polyacrylamide gel electrophoresis (BN-PAGE) was performed to isolate mitochondrial proteins from tissues as described previously[8]. In-gel activity assays for oxidative phosphorylation (OXPHOS) complexes were conducted according to established protocols[8, 9]. Briefly, 30 μg of total mitochondrial protein was separated on a 3 – 12% Bis-Tris Native PAGE gel. Following electrophoresis, gels were pre-washed in cold water and then incubated with specific substrates for each complex at room temperature for 2 hours: Complex I : 0.1 mg/mL NADH, 2.5 mg/mL nitroblue tetrazolium (NBT), 2 mM Tris-HCl (pH 7.4); Complex II: 20 mM sodium succinate, 2.5 mg/mL NBT, 0.2 mM phenazine methosulfate (PMS), 5 mM Tris-HCl (pH 7.4) and Complex IV : 0.5 mg/mL DAB, 1 mg/mL cytochrome c, 45 mM phosphate buffer (pH 7.4). Reactions were stopped by immersion in 10% acetic acid, followed by washing with distilled water. Gels were scanned to visualize the formation of insoluble formazan (Complexes I and II) or oxidized DAB precipitates (Complex IV), indicating enzymatic activity.

To estimate the in-gel ATP hydrolysis activity of complex V, gel strips were preincubated in 50 mm glycine (adjusted to pH 8.0 with triethanolamine) at 37 °C for 2 hours. The equilibration solution was discarded, and the gel strips were transferred to assay buffer containing 50 mM glycine (pH 8.0, adjusted with triethanolamine), 10 mM $MgCl_2$, 0.2% $Pb(NO_3)_2$, and 8 mM ATP.

ATP hydrolysis correlated with the formation of white lead phosphate precipitates. The reaction was terminated by immersion in 50% methanol, followed by transfer to distilled water. Gels were then scanned against a dark background to enhance contrast and visualize precipitate formation.

**Real-time Quantitative PCR (RT-qPCR)**

Total RNA was extracted from cells or tissues using the RNA-easy Isolation Reagent (Vazyme, R701), followed by cDNA synthesis using the ABScript III RT Master Mix for qPCR (ABclonal, RK20429). Gene expression was analyzed by Universal SYBR Green Fast qPCR Mix (ABclonal, RK21203). The relative gene expression levels were calculated using the $2^{-\Delta\Delta Ct}$ method. The primers used were as follows: hMISO-F: TTGAGAAGCATTCGACTGACA, hMISO-R: GGCGTAATCGTCCACGTAGT; hActin-F: CTCCATCCTGGCCTCGCTGT, hActin-R: GCTGTCACCTTCACCGTTCC; hND2-F: CTATCACCCTATTAACCACTCA, hND2-R: TTCGCCTGTAATATTGAACGTA; hB2MG-F: CTATGGGACGCTTGATGT, hB2MG-R: GCAATCATTCGTCTGTTT; mND1-F: CTAGCAGAAACAAACCGGGC, mND1-R: CCGGCTGCGTATTCTACGTT; mHK2-F: GCCAGCCTCTCCTGATTTTAGTGT, mHK2-R: GGGAACACAAAAGACCTCTTCTGG; dMISO-F: TTTAAGCAGCCCTCGCACAT dMISO-R: GGATCATCAGCAGGGTGTCC dTubulin-F: TGTCGCGTGTGAAACACTTC; dTubulin-R: AGCAGGCGTTTCCAATCTG

**Measurement of Oxygen Consumption Rate**

To measure the oxygen consumption rates (OCR) of cells, an XFe96 extracellular flux analyzer (Seahorse Bioscience; Agilent Technologies) was utilized. Briefly, 10,000 cells per well were seeded in a XF96 cell culture microplate. After allowing the cells to adhere and recover, the medium was aspirated, and the cells were washed twice with XF assay medium (XF base medium supplemented with 10 mM glucose, 5 mM pyruvate, and 2 mM L-glutamine). The cells were then incubated in this assay medium within a non-$CO_2$ incubator at 37 °C for 1 hour. The OCR values were recorded under basal conditions and following sequential injections of drugs at the following final concentrations: 1 μM oligomycin, 1 μM FCCP, and a combination of 1 μM rotenone and 1 μM antimycin A. Total protein amount was used as normalization for OCR.

**Flow Cytometry Analysis of Mitochondrial Membrane Potential**

Mitochondrial membrane potential was determined by TMRM according to the manufacturer's manual. In brief, cells were incubated with 50 nM TMRM at 37 °C for 30 minutes. After washing three times with PBS, cells were trypsin-digested and suspended in HBSS. The TMRM fluorescence intensity was detected by flow cytometry using a CytoFLEX flow cytometer (Beckman Coulter). Data analysis was performed using FlowJo software.

**Imunoprecipitation and Mass Spectrometry Analysis**

Cells were lysed in a lysis buffer composed of 150 mM NaCl, 50 mM Tris-HCl at pH 8.0, 1 mM EDTA, and 0.5% Triton X-100, enhanced with both protease inhibitor cocktail and phosphatase inhibitor cocktail (Beyotime, P1081). After lysis, samples underwent centrifugation at 12,000×g for 20 minutes at 4 °C, after which the supernatant was collected. The collected supernatant was then incubated with Anti-FLAG M2 Magnetic Beads (Sigma-Aldrich, M8823) for a period of 2 to 4 hours at 4 °C under gentle rotation. Upon completion of the incubation, the beads were isolated using a magnetic stand and subjected to three washes with lysis buffer to eliminate non-specific protein binding. Finally, the SDS loading buffer was applied to the beads in preparation for subsequent Western blot analysis.

For MS sample preparation, the FLAG magnetic beads were washed twice with TBS (137 mM NaCl, 20 mM Tris, pH 7.6), and the Flag-tagged protein was eluted with TBS supplemented with 100 μg/ml 3×FLAG peptide (Beyotime, P9801). The eluted proteins were denatured, reduced and alkylated, and then digested by trypsin (1/50 w/w) at 37 °C for 20 hours. Subsequently, the peptides were desalted, concentrated with C18 tips, and resuspended in 0.1% FA (formic acid) for mass spectrometry analysis. Samples were analyzed by liquid chromatography/tandem mass spectrometry (Easy-nLC 1000+Q Exactive Orbitrap mass spectrometer, Thermo Fisher) at Shanghai Applied Protein Technology Co. Ltd. The MA raw data were analyzed using MASCOT v2.2 and protein database with the following parameters:

Database: uniport_Homo_sapiens_194324_20210106; Fixed modifications: Carbamidomethyl (C); Variable modifications: Oxidation (M); Missed Cleavage: 2; Peptide Mass Tolerance: 20 ppm; Fragment Mass Tolerance: 0.1 Da; Filter by score ≥ 20.

**Polyethylene Glycol (PEG)-Mediated Mitochondrial Fusion**

HeLa cells stably expressing Mito-RFP were transfected with either an empty vector (pcDNA3.1) or MISO-Flag. After 20 hours, the cells were harvested using trypsin, counted, and mixed in equal proportions with cells stably expressing Mito-GFP. The mixed cells were co-seeded onto glass coverslips and allowed to adhere for 24 hours before the experiment. For the fusion assay, cells on the coverslips were incubated with pre-warmed 50% (w/v) PEG1500 solution (Zeye Biology, ZY2261a) at 37 °C for 90 seconds. Following this incubation, the cells were gently washed and then cultured in complete medium supplemented with 50 μg/ml cycloheximide (MedChemExpress, HY-12320) to maintain inhibition of protein synthesis. Finally, cells were fixed with 4% PFA and prepared for microscopy analysis.

**Image analysis**

**Mitochondrial morphology analysis**

Mitochondrial morphology was analyzed as previously described[10]. Z-stack images were acquired in a randomized manner, then compiled into maximum intensity projections. Mitochondrial morphologies were categorized as fragmented, intermediate, hyperfused, or aggregated. Mitochondria were classified as fragmented when the majority appeared short and spherical, intermediate when predominantly tubular with no extensive interconnections or spherical forms, hyperfused when mitochondria exhibited highly elongated networks, and aggregated when mitochondria were densely clustered in the perinuclear region. For mitochondrial morphology analysis in *Drosophila* intestinal stem cells, fragmented mitochondria were defined as those appearing predominantly as short and spherical structures, while hyperfused mitochondria were characterized by extensive fusion, forming one or two highly interconnected networks.

Quantification of mitochondrial morphological parameters was performed according to established protocols[11, 12]. A defined region of interest (ROI) of 225 $\mu m^2$ was selected at the cell periphery for analysis using ImageJ software. Briefly, maximum intensity projections were generated from Z-stacks, followed by automated thresholding using the *Otsu* method. Thresholded images were then processed using the "Analyze Particles" function to determine the number of mitochondrial fragments, the area of each fragment per ROI and the perimeter of each fragment per ROI. To assess the mitochondrial junctions, thresholded images were skeletonized using the *Analyze Skeleton* plugin, and further counted by *Analyze Particles*. At least 15-20 cells per condition were analyzed from at least three independent experiments.

To Quantification of the number of MISO-enriched subdomains and mitochondrial length, fluorescence channels were separated, and images were thresholded using the *Advanced Weka Segmentation* plugin. The mitochondrial channel was skeletonized, and analyzed via *Analyze Particles* to determine average mitochondrial length per ROI. The MISO signal was processed using the *Watershed* algorithm to resolve closely apposed subdomains, followed by particle analysis to count the number of MISO-enriched puncta per ROI.

**Quantification of mtDNA foci**

The mtDNA foci number was quantified using maximum intensity projection images. Images were first processed to reduce background noise and enhance signal clarity by applying a *Despeckle* filter and background subtraction (Rolling Ball radius = 50 pixels). Following manual thresholding to preserve nucleoid-specific signal, particle analysis was performed using the *Analyze Particles*. At least 20 cells per condition were analyzed from at least three independent experiments.

**Colocalization analysis**

Colocalization between MISO-enriched mitochondrial subdomains and specific mitochondrial proteins or cellular organelles was quantified using FIJI. Pearson's correlation coefficient and Mander's overlap coefficient were calculated using the *Colocalization Finder* plugin to assess the degree of spatial overlap between MISO and target signals.

To further validate colocalization at suborganellar resolution, line-scan intensity profiles were generated across regions of interest using *Plot Profile*. Fluorescence intensities for all relevant channels were plotted along the defined line to visualize spatial correspondence of signals. Mitochondrial-associated DRP1 punctate were identified by the *Colocalization Finder* plugin, then counted using the *Analyze Particles* function in ImageJ.

**Number of samples for microscopy analysis**

**Fig. 1:** (**i**): *n* = 460 cells from 16 fly midguts for *Luc-i*, *n* = 350 cells from 22 fly midguts for *dMISO-i*. (**l**): *n* = 500 cells from 15 fly midguts for Control, *n* = 511 cells from 18 fly midguts for *dMISO-OE*.

**Fig 2:** (**c**): *n* = 411, 531 cells for WT and MISO-KO, respectively. (**d**): Mitochondrial number/area/perimeter: *n* = 60, 60 for WT and KO, respectively. Mitochondrial junctions: *n* = 75 cells per group. (**f**): *n* = 408, 366, 348 cells for pcDNA3.1, MISO iso1-Flag and MISO iso2-Flag, respectively. (**g**): Mitochondrial number/area/perimeter: *n* = 60, 69, 61 cells for pcDNA3.1, MISO iso1-Flag and MISO iso2-Flag, respectively. Mitochondrial junctions: *n* = 75 cells per group. (**k**): *n* = 60, 81 for WT and KO, respectively.

**Fig 3:** (**c**): Cells examined at each post-induction time point were as follows: n = 36 at 4 h, *n* = 46 at 6 h, *n* = 37 at 8 h, *n* = 26 at 10 h, and *n* = 27 at 12 h. HSP60 signal intensity was quantified in n = 13 uninduced cells and in n = 26 (4 h), n = 25 (6 h), n = 21 (8 h), n = 18 (10 h), and n = 18 (12 h) induced cells. (**f**): *n* = 20, 20 for TOMM20 and COX5B staining, respectively. (**j**): A total of *n* = 332 from at least 140 cells were analyzed. (**l**): *n* = 100 cells were examined. (**o**): A total of *n* = 259 mitochondrial fission events from at least 45 cells were examined. (**q**): *n* = 187, 220, 173 for Type I, Type II, Type III mitochondrial fusion events, respectively, from at least 45 cells. (**r**): For each time point, 21 individual hybrid cells for pcDNA3.1 and MISO-Flag, respectively.

**Fig 4:** (**b**): *n* = 30 cells for mtDNA/Mito, mtDNA/MISO, mtRNA/Mito and mtDNA/MISO. (**c**): *n* = 10 MISO-enriched subdomians from 5 cells were examined. (**e**): n = 50, 50 for pcDNA3.1 and MISO-Flag, respectively. (**o**) and (**p**): *n* = 73, 100 for WT and KO, respectively.

(**t**) and (**u**): *n* = 38, 35, 36, 36, 36 cells for shNT, shDRP1, shFIS1, shMTFP1-1 and shMTFP1-2, respectively.

**Fig 5:** (**a**): *n* = 347, 318, 313 and 320 cells for WT (Veh), WT (EB 3d), KO (Veh) and KO (EB 3d), respectively. (**d**): *n* = 145, 139, 158 and 110 cells for WT (Veh), WT (EB 5d), KO (Veh) and KO (EB 5d), respectively. (**e**): *n* = 60, 72, 65 and 66 mitochondria for WT (Veh), WT (EB 5d), KO (Veh) and KO (EB 5d), respectively.

**Fig 6:** (**b**): *n* = 122, 150 for pLenti and MISO-Flag, respectively. (**d**): *n* = 228, 220 cells for pLenti and MISO-Flag, respectively. (**h**): *n* = 397, 445 cells for APEX2-MISO and MISO-APEX2, respectively. (**n**): At least 300 cells for OMM-GFP, *n* = 488 cells for OMM-MISO. (**p**): *n* = 387, 368 cells for IMM-GFP and IMM-MISO, respectively.

**Extended Data Fig. 3:** (**d**): *n* = 44, 57 mitochondria for MISO-WT and MISO-KO, respectively. (**g**): *n* = 45 cells for WT (pLenti), WT (MISO-Flag iso1), WT (MISO-Flag iso2), KO (pLenti), KO (MISO-Flag iso1) and KO (MISO-Flag iso2).

**Extended Data Fig. 5:** (**b**): *n* = 426, 484, 424 cells for shNT, shMISO-1 and shMISO-2, respectively. (**e**): *n* = 395, 431, 350 cells for shNT, shMISO-1 and shMISO-2, respectively. (**g**): *n* = 376, 495, 361 cells for shNT, shMISO-1 and shMISO-2, respectively.

**Extended Data Fig. 6:** (**d**). *n* = 401, 610 cells for $mMISO^{flox/flox}$, and $mMISO^{-/-}$, respectively. (**f**): *n* = 422, 412 cells for $mMISO^{flox/flox}$, and $mMISO^{-/-}$, respectively. (**h**): *n* = 32, 34 mitochondria for $mMISO^{flox/flox}$, and $mMISO^{-/-}$, respectively.

**Extended Data Fig. 7:** (**b**): *n* = 356, 404 cells for MISO-WT and MISO-KO, respectively. (**d**): *n* = 368, 323 cells for MISO-WT and MISO-KO, respectively. (**e**): Mitochondrial number/area/perimeter: *n* = 71, 65, 60 cells for MISO-Flag + shNT, MISO-Flag + shDRP1, MISO-Flag + shFIS1, respectively. Mitochondrial junctions: *n* = 60 cells per group.

**Extended Data Fig. 11:** (**c**): *n* = 20 for each staining.

**Extended Data Fig. 12:** (**b**): *n* = 20, 17, 21, 21, 21, 17, 19, 19, 21, 16, 22, 20, 20, 22, 18, 19 subdomains for SLC25A4-Myc, SLC25A5-Myc, SLC25A6-Myc, PHB1-Myc, PHB2-Myc, TIMM23, VDAC1-HA, VDAC2-HA, COX5B, TOMM20, TOMM40, TOMM70, SDHB, STOML2, ATAD3A-Myc and ATAD3B-Myc, respectively.

**Extended Data Fig. 16:** (**d**): *n* = 321, 303, 294 cells for MISO-Flag, MISO-Flag + MFN1-mCherry and MISO-Flag + MFN2-mCherry, respectively. (**f**): *n* = 45 cells for WT (shNT), WT (shMFN1), WT (shMFN2), WT (shOPA1), KO (shNT), KO (shMFN1), KO (shMFN2) and KO (shOPA1).

**Extended Data Fig. 17:** (**b**): *n* = 26, 27, 26, 28, 30, 28 cells for Mito, ER, Golgi, Lyso, Perox and LD, respectively.

**Extended Data Fig. 18:** (**c**): *n* = 21, 21cells for Vehicle and EB (3d), respectively.

**Extended Data Fig. 23:** (**f**): *n* = 45 cells for pLenti (shNT), pLenti (shMTFP1-1), pLenti (shMTFP1-2), MISO-Flag (shNT), MISO-Flag (shMTFP1-1), MISO-Flag (shMTFP1-2), **(f):** *n* = 329, 359 cells for WT and KO, respectively. (**g**): *n* = 45 cells for WT (pcDNA3.1), WT (MTFP1-Myc), KO (pcDNA3.1) and KO (MTFP1-Myc).

**Extended Data Fig. 26:** (**b**): *n* = 20, 25, 22, 23, 25, 20, 20, 20, 21, 23, 28 cell clones for *Luc-i*, *NP15.6-i*, *ND-49-i*, *ND-23-i*, *SdhC-i*, *UQCR-Q-i*, *RFeSP-i*, *COX6C-i*, *COX5A-i*, *ATPsyn-beta-i-1* and *ATPsyn-beta-i-2*, respectively. (**d**): *n* = 20, 20 for Luc-i and dMISO-i, respectively.

**Extended Data Fig. 27:** (**f**): *n* = At least 300 cells for WT (DMSO) and KO (DMSO), *n* = 406, 365 cells for WT (CCCP) and KO (CCCP), respectively.

**Extended Data Fig. 28:** (**b**): *n* = 427, 426, 429, 434, 413, 382 cells for WT (shNT), WT (shMIC60), WT (shATP5A), KO (shNT), KO (shMIC60) and KO (shATP5A), respectively.

**Extended Data Fig. 30:** (**e**): At least 300 cells for OMM-GFP, *n* = 201 cells for OMM-MISO. (**g**): At least 300 cells for OMM-GFP, OMM-MISO-M and OMM-MISO-C, *n* = 341 cells for OMM-MISO. (**j**): *n* = 350, 329, 400 and 338 cells for IMM-GFP, IMM-MISO, IMM-MISO-M and IMM-MISO-C, respectively.

**Statistical Analysis**

All experiments in this paper were repeated at least three times. Data analyses were performed with Prism 9 (GraphPad software, San Diego, CA, USA) and presented as mean ± SD unless specified. Data in Mitochondrial membrane potential assay was analyzed with FlowJo software. Multiple statistical methods were employed to analyze differences, as indicated in the figure legends. $p < 0.05$ was considered statistically significant.

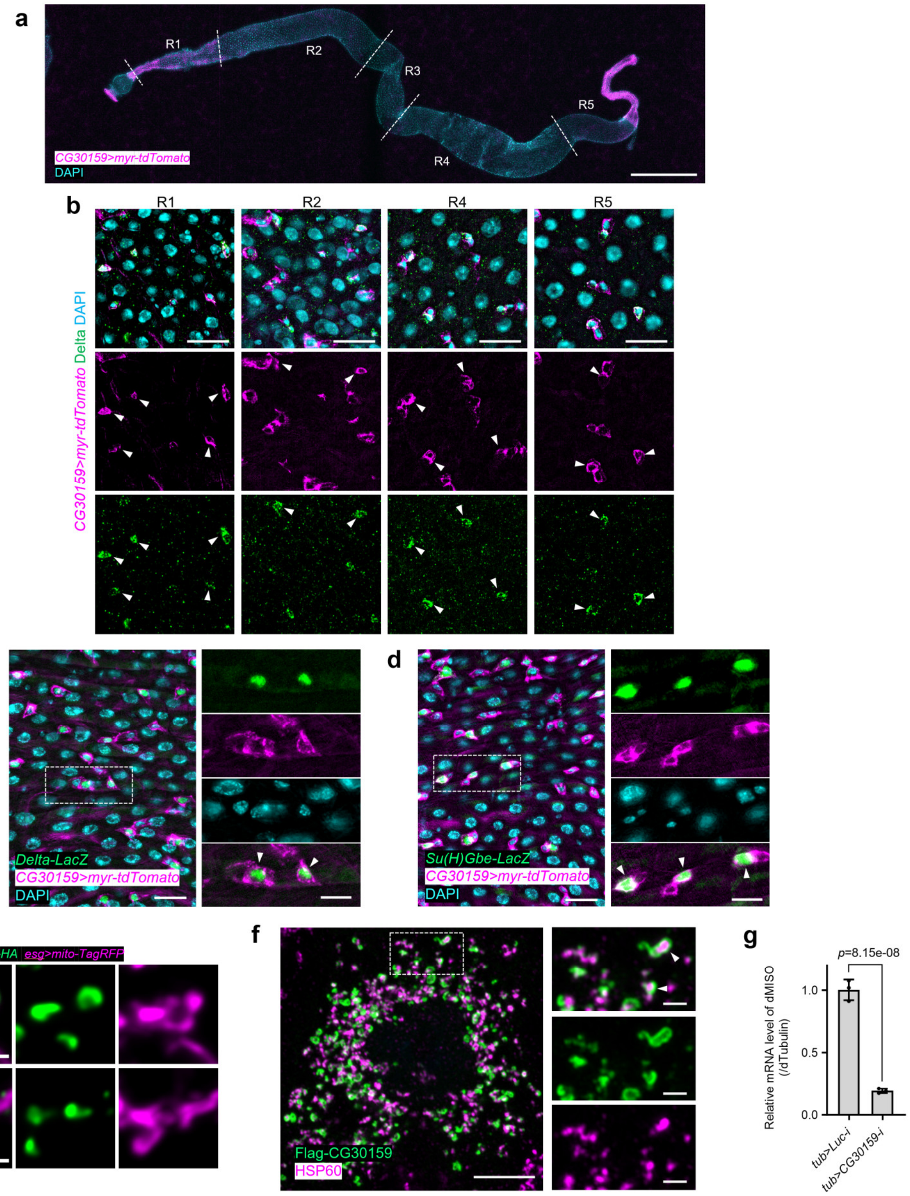


**Extended Data Fig. 1. *CG30159* is expressed in *Drosophila* intestinal stem cells (ISCs) and forms mitochondrial subdomains in cultured insect cells.**

**a.** Expression pattern of *CG30159-Gal4* in fly intestine visualized with myr-tdTomato. DAPI staining indicates the cell nuclei throughout the figure. Scale bars: 500 µm.

**b.** *CG30159* is expressed in ISCs identified by Delta staining (green) in regions R1, R2, R4 and R5 of the intestine. White arrowheads point to ISCs. Scale bars: 20 µm.

**c.** CG30159-Gal4 driven tdTomato co-staining with *Dl-LacZ* (green). The white arrowheads point to *Dl* positive stem cells. Scale bars: main panels 20 µm, magnified insets 10 µm.

**d.** CG30159-Gal4 driven tdTomato co-staining with *Su(H)Gbe-LacZ* (green). The white arrowheads point to *Su(H)Gbe* positive enteroblasts. Scale bars: main panels 20 μm, magnified insets 10 μm.

**e.** Representative images of CG30159 immunofluorescence staining in *esg*+ stem cells. Yellow arrowheads point to membrane subdomains. Scale bar: 1 μm.

**f.** Representative images of immunofluorescence staining for Flag-tagged CG30159 in S2 cells. Scale bars: main panels 5 μm, magnified insets 2 μm.

**g.** qPCR analysis of *CG30159* knockdown efficiency in adult flies. *n* = three experiments.

All data are presented as mean ± SD. (**f**): two-tailed unpaired t test.

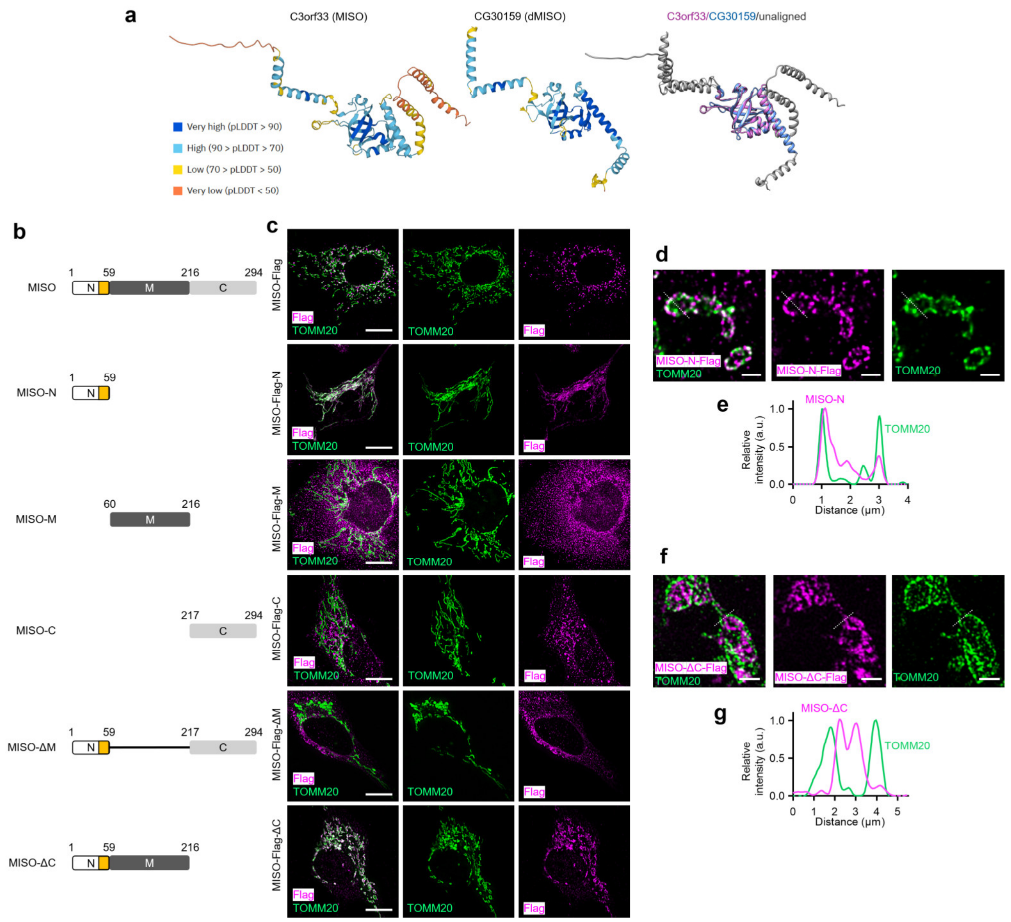


**Extended Data Fig. 2. Structural and truncation analysis of MISO.**

**a.** AlphaFold structure prediction of CG30159 (dMISO) and its human orthologue C3orf33 (MISO). The prediction confidence scores (0-100) are visually represented using a color scale. The alignment of the structures of the two is displayed on the right, with aligned regions superimposed and highlighted in color.

**b.** Schematic representation of full-length MISO and corresponding truncation mutants. transmembrane domains are highlighted in yellow.

**c.** Representative images of U2OS cells expressing Flag-tagged full-length MISO and corresponding truncation mutants. Scale bars: 10 μm.

**d-f.** Representative expansion microscopy images (**d**) and line profiles (**f**) showing the localization of MISO-N-Flag and TOMM20 in U2OS cells. Scale bars: 2 μm (post-expansion).

**g-h.** Representative expansion microscopy images (**g**) and line profiles (**h**) showing the localization of MISO-ΔC-Flag and TOMM20 in U2OS cells. Scale bars: 5 μm (post-expansion).

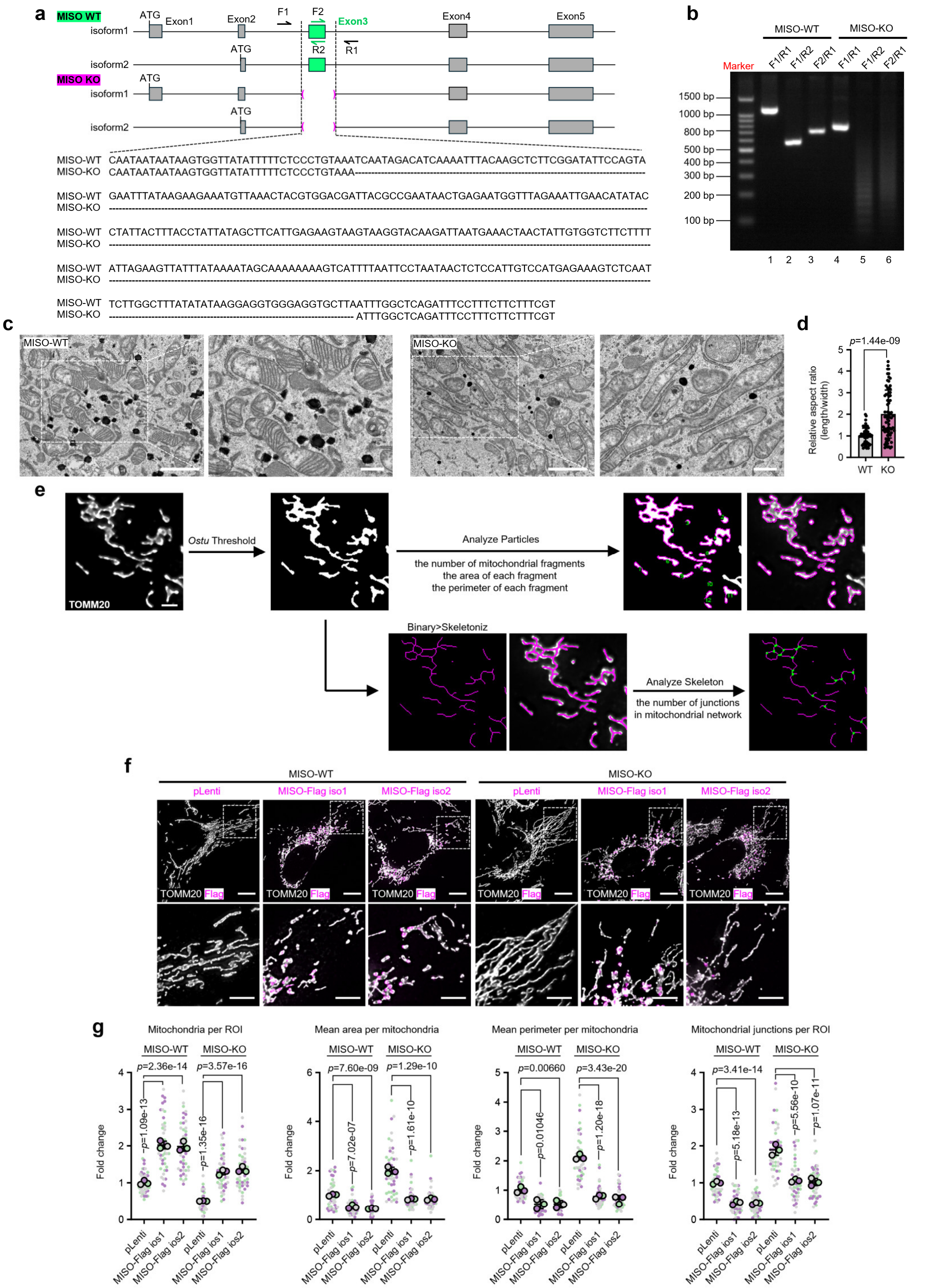


**Extended Data Fig. 3. Characterization of mitochondrial morphology in MISO-KO cells.**

**a.** Sequence analysis of *MISO* alleles in wild-type (WT) and CRISPR/Cas9-generated MISO-knockout (MISO-KO) U2OS cells.

**b.** PCR-based genotyping of *MISO* locus in WT and MISO-KO U2OS cells.

**c-d.** Representative TEM images (**c**) and corresponding quantification (**d**) of mitochondrial morphology in WT and MISO-KO U2OS cells. Scale bars: main panels 2 μm, magnified insets 500 nm.

**e.** Representative images illustrating the methodology for the quantitative analysis of mitochondrial morphology using Fiji. Scale bars: 2 μm.

**f-g.** Representative images (**f**) and corresponding quantification (**g**) of mitochondrial morphology in WT and MISO-KO U2OS cells expressing an empty vector (pLenti), MISO-Flag isoform 1 (MISO-Flag iso2) or MISO-Flag isoform 2 (MISO-Flag iso2). Scale bars: main panels 10 μm, magnified insets 5 μm. *n* = three experiments.

All data are presented as mean ± SD. (**d**): non-parametric Mann-Whitney test; (**g**): two-tailed nested t test.

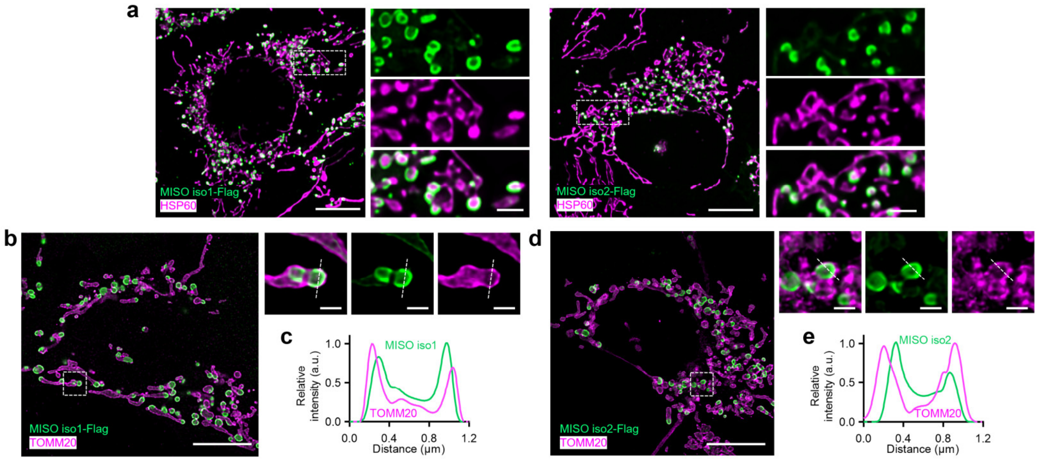


**Extended Data Fig. 4. Both MISO isoforms induce the formation of similar IMM subdomains.**

**a.** Representative images show that both MISO isoform 1 and isoform 2 are capable of establishing mitochondrial subdomains. Scale bars: main panels 10 µm, magnified insets 2 µm.

**b-c.** Representative images (**b**) and line profiles (**c**) showing the localization of MISO-iso1 and TOMM20 in U2OS cells. Scale bars: main panels 10 µm, magnified insets 1 µm.

**d-e.** Representative images (**d**) and line profiles (**e**) showing the localization of MISO-iso2 and TOMM20 in U2OS cells. Scale bars: main panels 10 µm, magnified insets 1 µm.

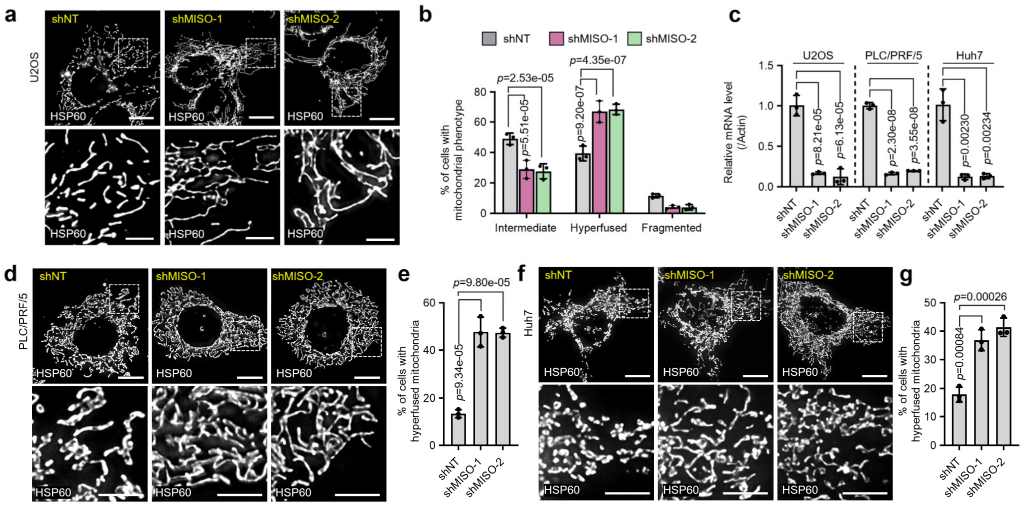


**Extended Data Fig. 5. MISO knock-down phenotypes in different cell lines.**

**a-b.** Representative images (**a**) and corresponding quantification (**b**) of mitochondrial morphology in U2OS cells treated with the indicated shRNAs. NT: non-targeted. Scale bars: main panels 10 μm, magnified insets 5 μm. *n* = three experiments.

**c.** qPCR analysis of MISO knockdown efficiency in various cell lines treated with the indicated shRNAs. *n* = three experiments.

**d-e.** Representative images (**d**) and corresponding quantification (**e**) of mitochondrial morphology in PLC/PRF/5 cells treated with the indicated shRNAs. NT: non-targeted. Scale bars: main panels 10 μm, magnified insets 5 μm. *n* = three experiments.

**f-g.** Representative images (**f**) and corresponding quantification (**g**) of mitochondrial morphology in Huh7 cells treated with the indicated shRNAs. Scale bars: main panels 10 μm, magnified insets 5 μm. *n* = three experiments.

All data are presented as mean ± SD. (**b**): two-way ANOVA with Tukey's multiple comparisons test; (**c**), (**e**) and (**g**): ordinary one-way ANOVA with Tukey's multiple comparisons test.

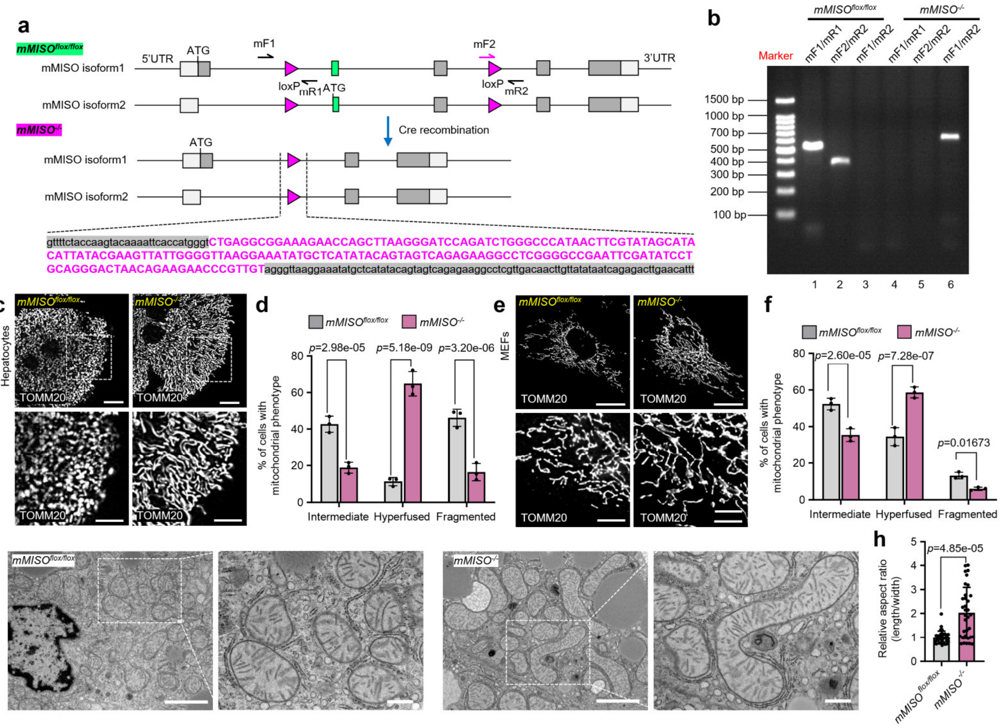


**Extended Data Fig. 6. Analysis of mitochondrial morphology in primary cells and liver tissue from MISO-knockout mice.**

**a.** Schematic of *mMISO/E130311K13Rik* flox allele and KO allele generated by cre recombination.

**b.** PCR-based genotyping of *mMISO* locus in *mMISO*$^{flox/flox}$ and *mMISO*$^{-/-}$ mice.

**c-d.** Representative images (**c**) and corresponding quantification (**d**) of mitochondrial morphology in *mMISO*$^{flox/flox}$ and *mMISO*$^{-/-}$ primary mouse hepatocytes. Scale bars: main panels 10 µm, magnified insets 5 µm. n = three experiments.

**e-f.** Representative images (**e**) and corresponding quantification (**f**) of mitochondrial morphology in *mMISO*$^{flox/flox}$ and *mMISO*$^{-/-}$ mouse embryonic fibroblasts (MEFs). Scale bars: main panels 20 µm, magnified insets 5 µm. n = three experiments.

**g-h.** Representative TEM images (**g**) and corresponding quantification (**h**) of mitochondrial morphology in liver cells from *mMISO*$^{flox/flox}$ and *mMISO*$^{-/-}$ mice. Scale bars: main panels 2 µm, magnified insets 500 nm.

All data are presented as mean ± SD. (**d**) and (**f**): two-way ANOVA with Tukey's multiple comparisons test; (**h**): non-parametric Mann-Whitney test.

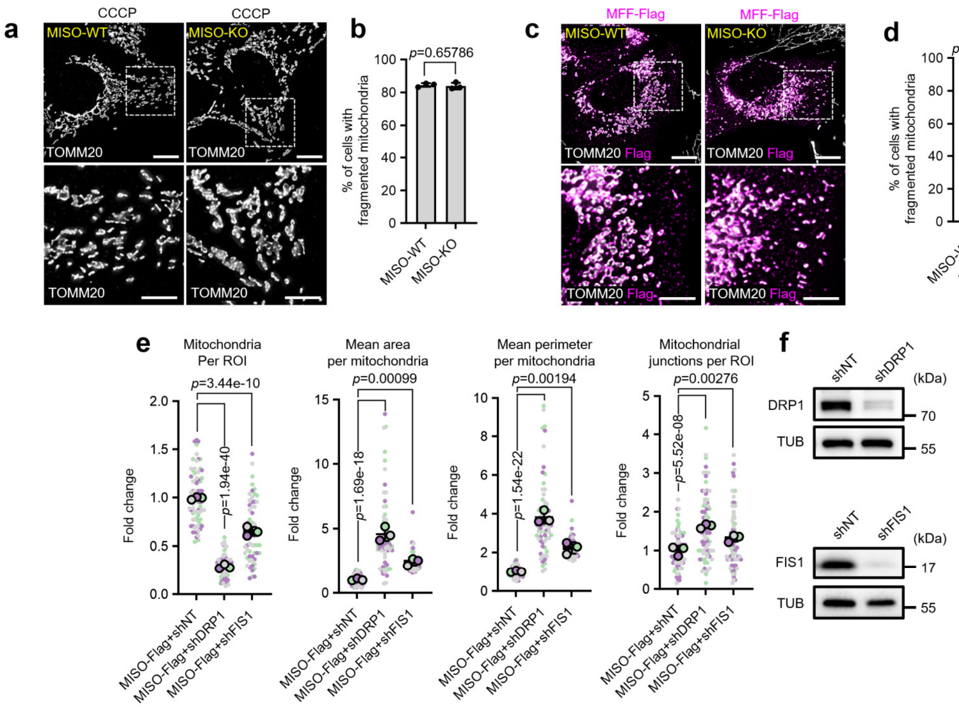


**Extended Data Fig. 7. Molecular mechanisms underlying MISO-mediated regulation of mitochondrial morphology.**

**a-b.** Representative images (**a**) and corresponding quantification (**b**) of mitochondrial morphology in WT and MISO-KO U2OS cells treated with CCCP. Scale bars: main panels 10 µm, magnified insets 5 µm. *n* = three experiments.

**c-d.** Representative images (**c**) and corresponding quantification (**d**) of mitochondrial morphology in WT and MISO-KO U2OS cells expressing MFF-Flag. Scale bars: main panels 10 µm, magnified insets 5 µm. *n* = three experiments.

**e.** Quantification of mitochondrial parameters from **Fig. 2** (**l**). *n* = three experiments.

**f.** Immunoblots showing DRP1 and FIS1 knockdown efficiency in U2OS cells.

All data are presented as mean ± SD. (**b**) and (**d**): two-tailed unpaired t test; (**e**): two-tailed nested t test.

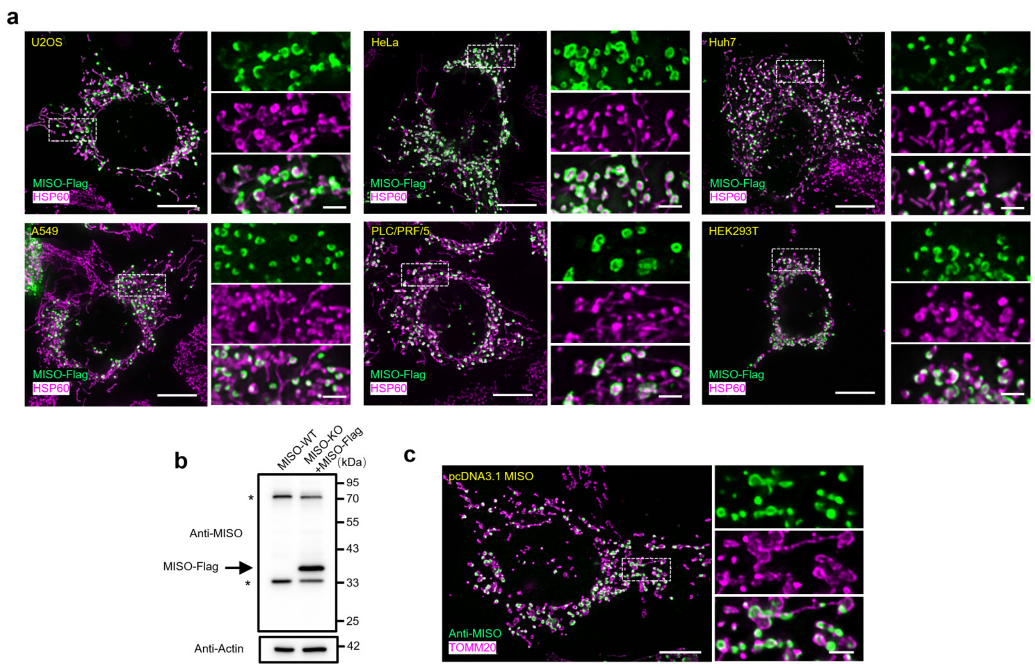


**Extended Data Fig. 8. MISO forms mitochondrial subdomains across multiple cell types independently of tag-fusion.**

**a.** Representative images showing the mitochondrial localization of MISO-Flag expressed in different cell types, including U2OS, HeLa, PLC/PRF/5, HEK293T, Huh7, and A549 cells. Mitochondria were marked with HSP60 (magenta). Scale bars: main panels 10 μm, magnified insets 2 μm.

**b.** Immunoblot analysis of lysates from MISO-WT and Miso-KO expressing MISO-Flag cells by Anti-MISO antibody. Arrowheads indicate the expected band, and asterisks denote nonspecific bands. Actin was used as a loading control.

**c.** Representative images of MISO immunofluorescence staining. U2OS cells transiently expressing untagged MISO were stained with an anti-MISO antibody. Scale bars: main panels 10 μm, magnified insets 5 μm.

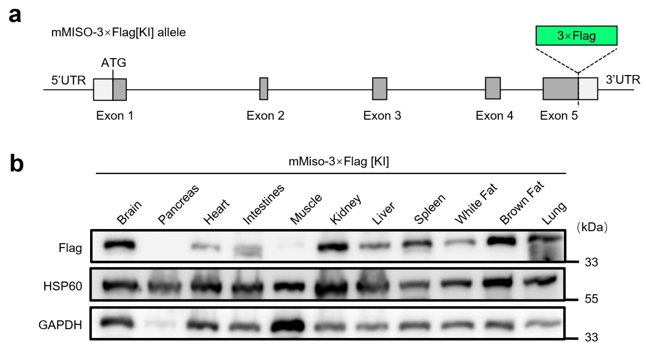


**Extended Data Fig. 9. Generation of mMISO-knockin mice and tissue-specific analysis of mMISO protein expression.**

**a.** A diagram of mMISO gene structure with 3×Flag tag knock-in element.

**b.** Immunoblot analysis was performed to evaluate the expression levels of endogenous mMISO in various mouse tissues. HSP60 and GAPDH were used as loading controls.

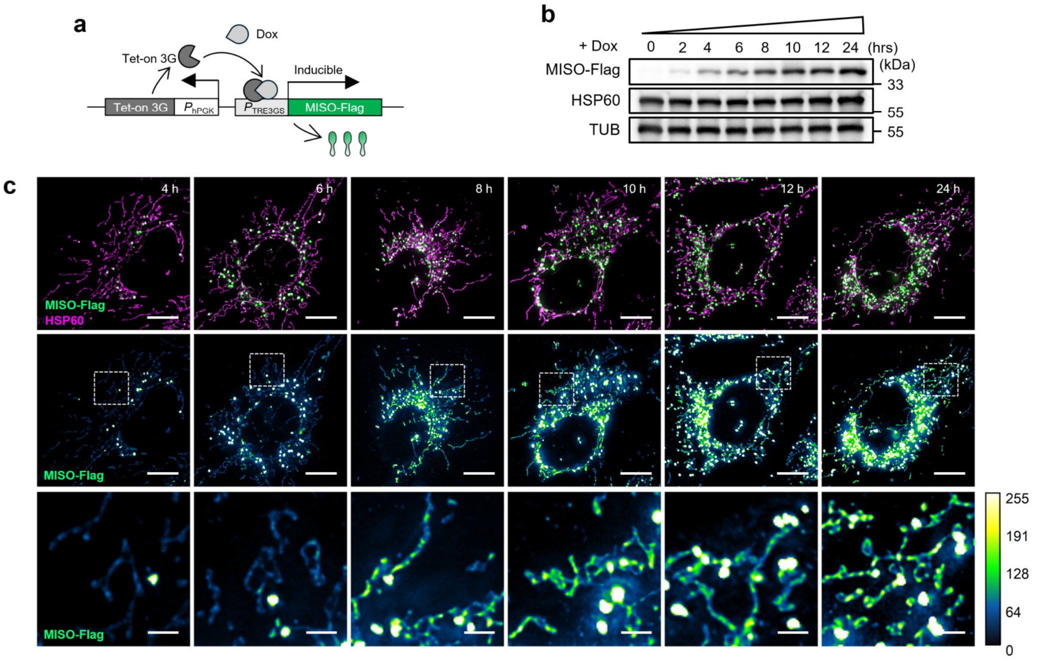


**Extended Data Fig. 10. MISO-enriched subdomains form across a range of MISO expression levels.**

**a.** A Scheme of the lentiviral cassette for the inducible expression of MISO-Flag.

**b.** Time course for doxycycline induction of MISO-Flag. U2OS cells were treated with 10 ng/ml doxycycline for various time intervals and cell lysates were analyzed by Western blot using the M2 antibody. Tubulin was used as a loading control.

**c.** Representative images showing MISO-Flag intensity in mitochondrial filaments and subdomains across a gradient of expression levels. Scale bars: main panels 10 µm, magnified insets 2 µm.

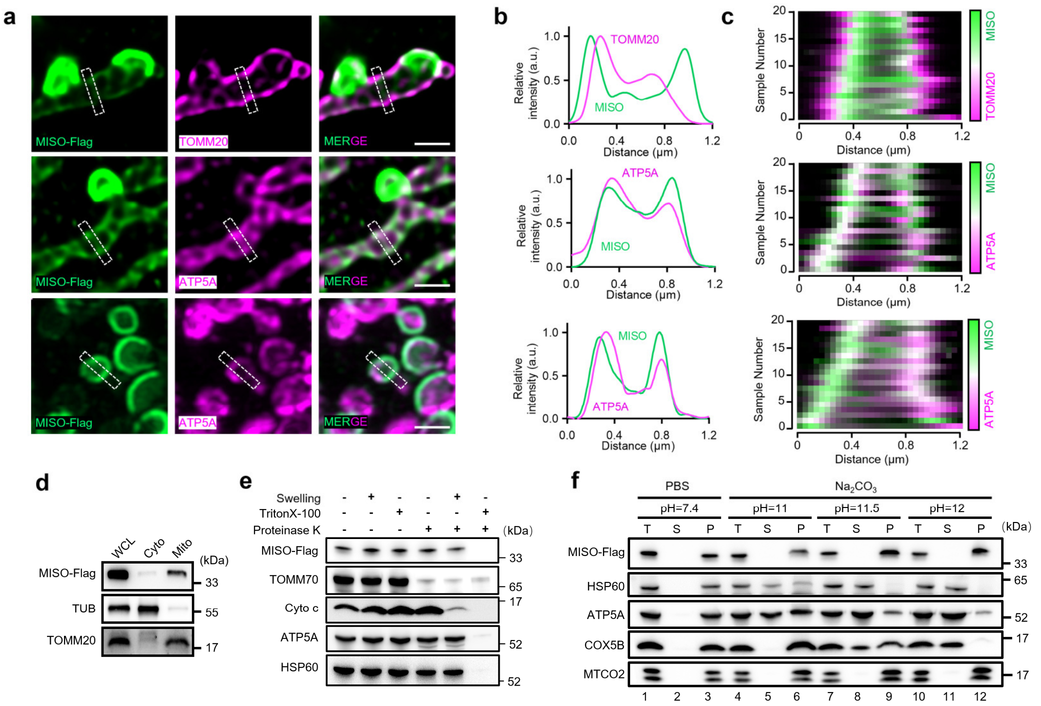


**Extended Data Fig. 11. MISO localizes to the inner mitochondrial membrane as an integral membrane protein.**

**a.** Super-resolution microscopy validating the localization of MISO to the inner mitochondrial membrane. U2OS cells expressing MISO-Flag were subjected to immunofluorescence staining to detect the FLAG epitope (green) and the outer mitochondrial membrane marker TOMM20 (upper panel, magenta), or the inner mitochondrial membrane marker ATP5A (lower panel, magenta), followed by imaging using STED microscopy. Scale bar: 1 μm.

**b.** Line profiles showing fluorescent signals of each channel across mitochondria, marked by the dotted rectangles in (**a**).

**c.** Fluorescence co-localization analysis of MISO (green) in mitochondria with TOMM20 (upper panel, magenta) or ATP5A (lower panel, magenta).

**d.** Cell fractionation assay of MISO-Flag stable expressing cells shows that MISO is localized in the mitochondria fraction, which is enriched with TOMM20. WCL: whole cell lysate; Cyto: cytoplasm; Mito: mitochondria.

**e.** Protease protection assay of mitochondria isolated from U2OS cells stably expressing MISO-Flag.

**f.** Mitochondrial protein extraction by sodium carbonate. Mitochondrial protein extraction by sodium carbonate treatment. Total mitochondria (T) isolated from U2OS cells stably expressing MISO-Flag were subjected to sodium carbonate extraction to separate peripheral membrane-bound proteins into the supernatant fraction (S), whereas integral membrane proteins remained in the pellet fraction (P).

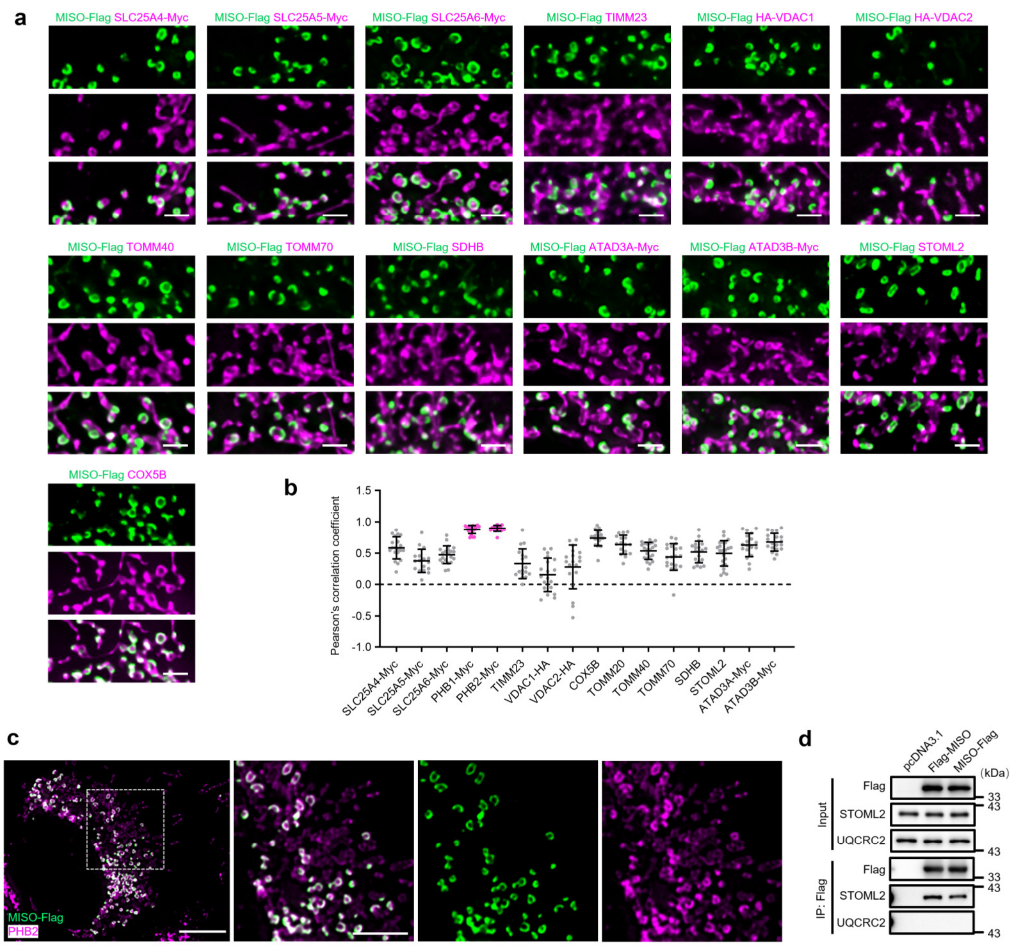


**Extended Data Fig. 12. Co-localization of MISO with different candidate mitochondrial proteins.**

**a-b.** Representative images (**a**) and corresponding co-localization quantification (**b**) of MISO with the indicated mitochondrial proteins in U2OS cells. Co-localization was assessed using Pearson's correlation coefficient. Scale bars: 2 µm.

**c.** Representative images demonstrating the colocalization of MISO-Flag (green) with endogenous PHB2 (magenta). Scale bars: main panels 10 µm, magnified insets 5 µm.

**d.** Immunoblots of the indicated proteins from anti-Flag immunoprecipitation of HEK293T cells overexpressing either an empty vector (pcDNA3.1), Flag-MISO or MISO-Flag.

All data are presented as mean ± SD.

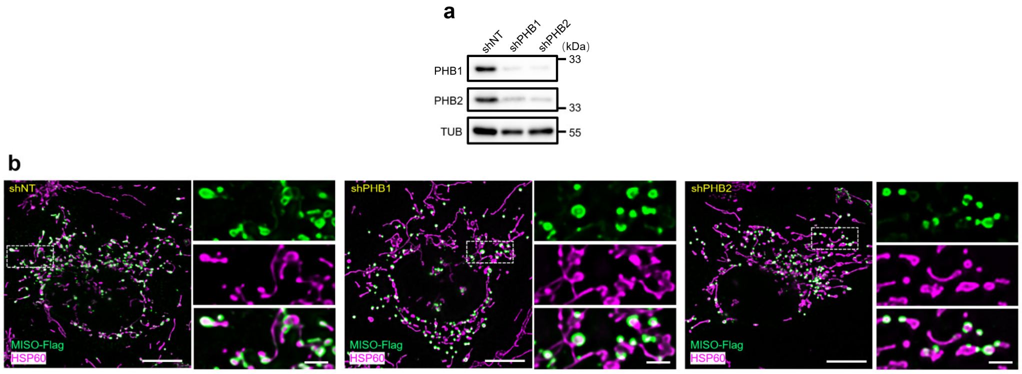


**Extended Data Fig. 13. PHBs are dispensable for subdomain formation.**

**a.** Western blot indicating the effective knockdown of PHB1 or PHB2 in U2OS cells. Tubulin as a loading control.

**b.** Representative images of mitochondrial subdomains in U2OS cells expressing MISO-Flag and treated with the indicated shRNAs. Scale bars: main panels 10 μm, magnified insets 2 μm.

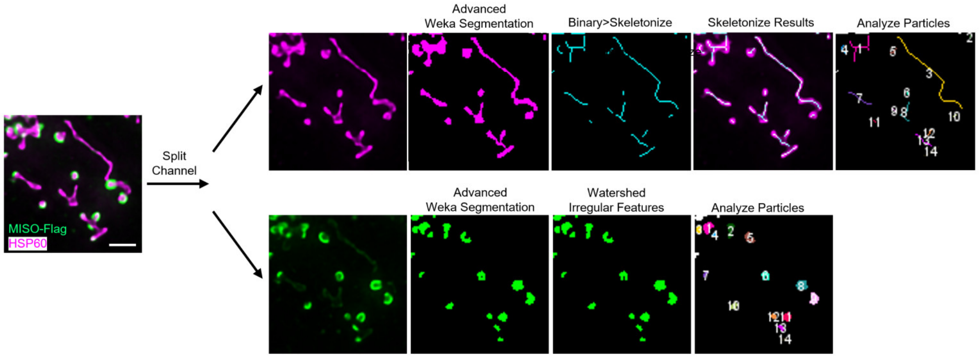


**Extended Data Fig. 14. Quantification of MISO-enriched subdomains and mitochondrial length.**

Representative images demonstrating the methodology for quantitatively analyzing the length of mitochondria and the number of subdomains using Fiji. Scale bars: 2 µm.

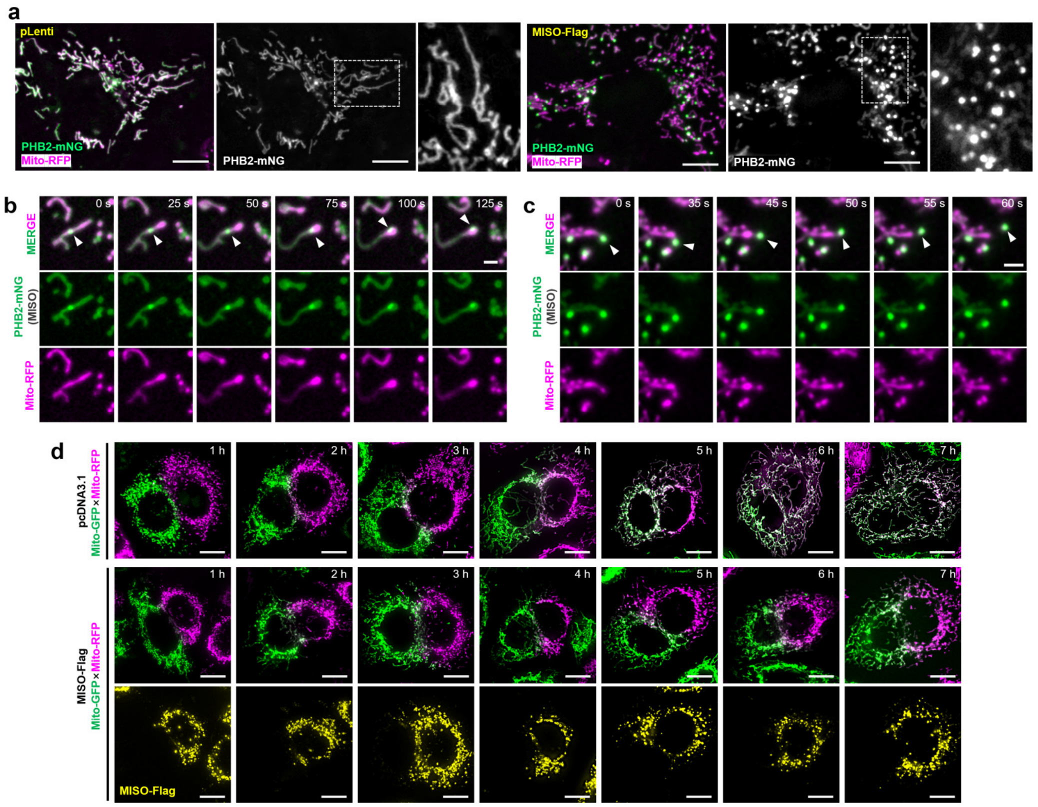


**Extended Data Fig. 15. MISO-enriched subdomains drive mitochondrial fission while suppressing fusion.**

**a.** Representative images showing the localization of PHB2-mNeonGreen in live U2OS cells, with or without MISO-Flag expression. Scale bars: 10 µm.

**b.** Representative time-lapse images showing the movement of subdomains towards the mitochondrial periphery in cells expressing MISO-Flag and PHB2-mNeonGreen (green) along with Mito-RFP (magenta). Scale bar: 2 µm.

**c.** Representative time-lapse images showing the dissociation of subdomain from the mitochondrial periphery. Scale bars: 2 µm.

**d.** Representative images of mitochondrial fusion in HeLa cells following PEG-mediated fusion assays. HeLa cells expressing Mito-GFP (green) and Mito-RFP (magenta) were transfected with the respective plasmids, co-cultured, and fused using PEG treatment. Left panel: Cells expressing Mito-RFP transfected with an empty vector; right panel: Cells expressing Mito-RFP transfected with Flag-tagged MISO. Scale bar: 10 µm.

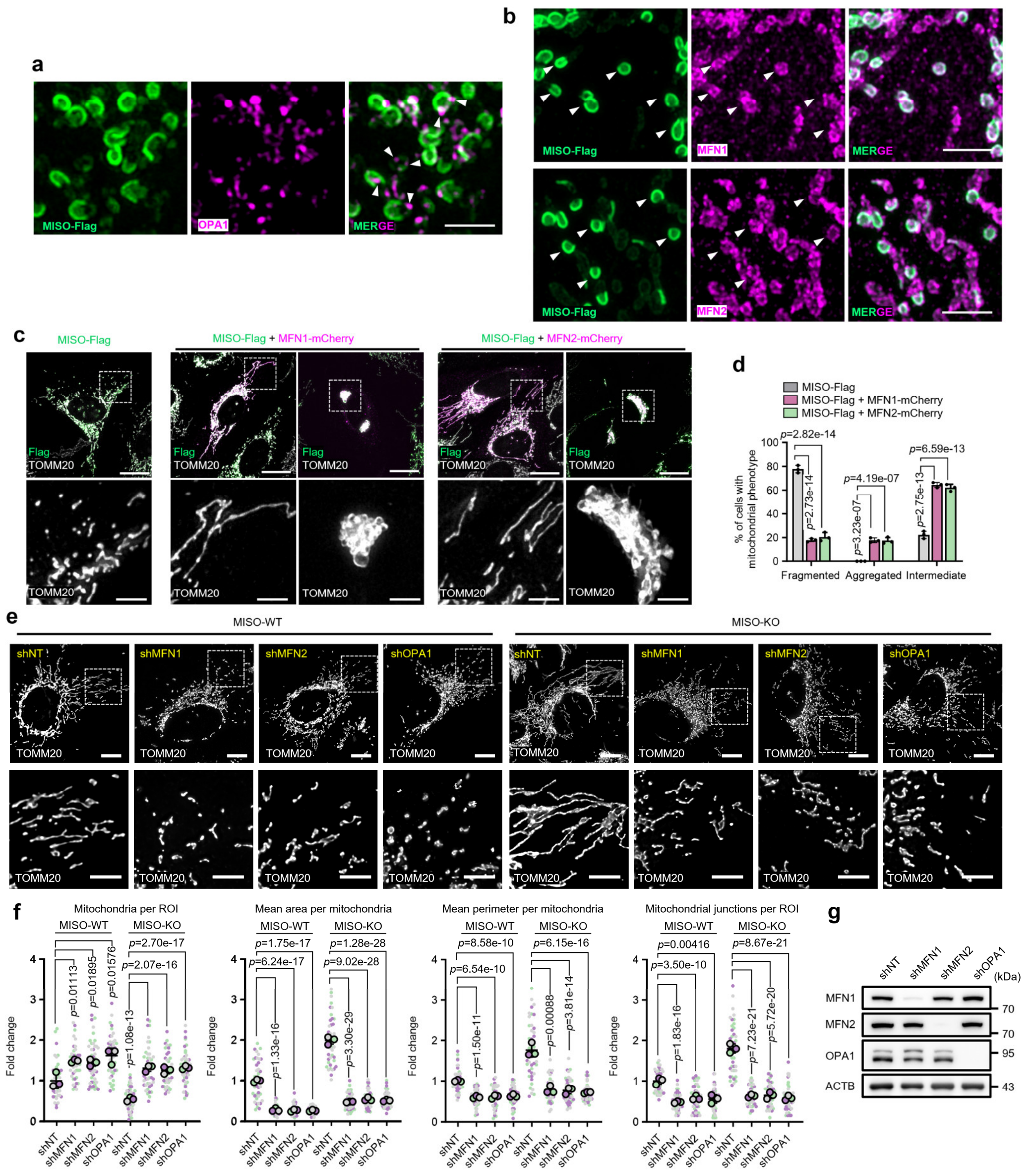


**Extended Data Fig. 16. The molecular mechanism by which MISO regulates mitochondrial fusion.**

**a.** Representative images showing the absence of localization of OPA1 with mitochondrial subdomains in U2OS cells expressing MISO-Flag. Scale bars: 2 μm.

**b.** Representative images showing the presence of localization of MFN1 and MFN2 with mitochondrial subdomains in U2OS cells expressing MISO-Flag. Scale bars: 2 μm.

**c-d.** Representative images (**c**) and corresponding quantification (**d**) of mitochondrial morphology in U2OS cells overexpressing MISO-Flag, with or without co-expression of

MFN1-mCherry or MFN2-mCherry. Scale bars: main panels 20 μm, magnified insets 5 μm. *n* = three experiments.

**e-f.** Representative images (**e**) and corresponding quantification (**f**) of mitochondrial morphology in WT and MISO-KO U2OS cells treated with the indicated shRNAs. Scale bars: main panels 10 μm, magnified insets 5 μm. *n* = three experiments.

**g.** Immunoblots showing the knockdown efficiency of MFN1, MFN2, and OPA1 in U2OS cells.

All data are presented as mean ± SD. (**d**): two-way ANOVA with Tukey's multiple comparisons test; (**f**): two-tailed nested t test.

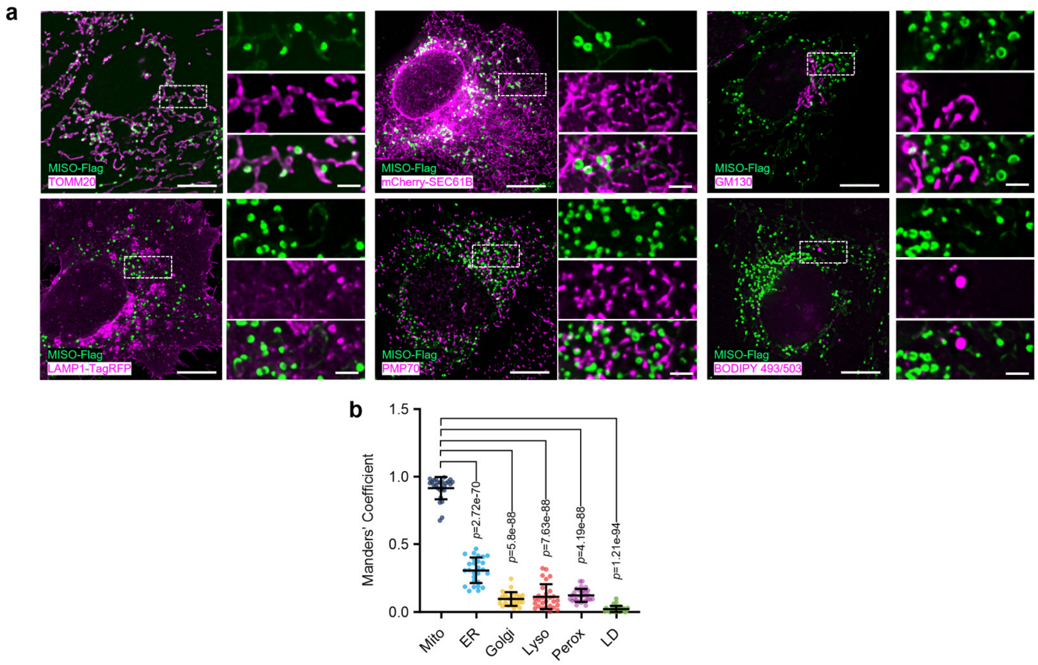


**Extended Data Fig. 17. Co-localization analysis of MISO-enriched subdomains with major organelles.**

**a.** Immunolocalization analysis was performed to determine the subcellular distribution of Flag-tagged human MISO alongside various cellular markers in U2OS cells, including TOMM20 (mitochondria), mCherry-SEC61B (endoplasmic reticulum), GM130 (Golgi apparatus), LAMP1-RFP (lysosome), PMP70 (peroxisome), and BIODIPY 493/503 (lipid droplets). Scale bars: main panels 10 μm, magnified insets 2 μm.

**b.** Co-localization analysis with quantification of Mander's coefficient between MISO-Flag and various cellular markers.

All data are presented as mean ± SD. (**b**): ordinary one-way ANOVA with Sidak's multiple comparisons test.

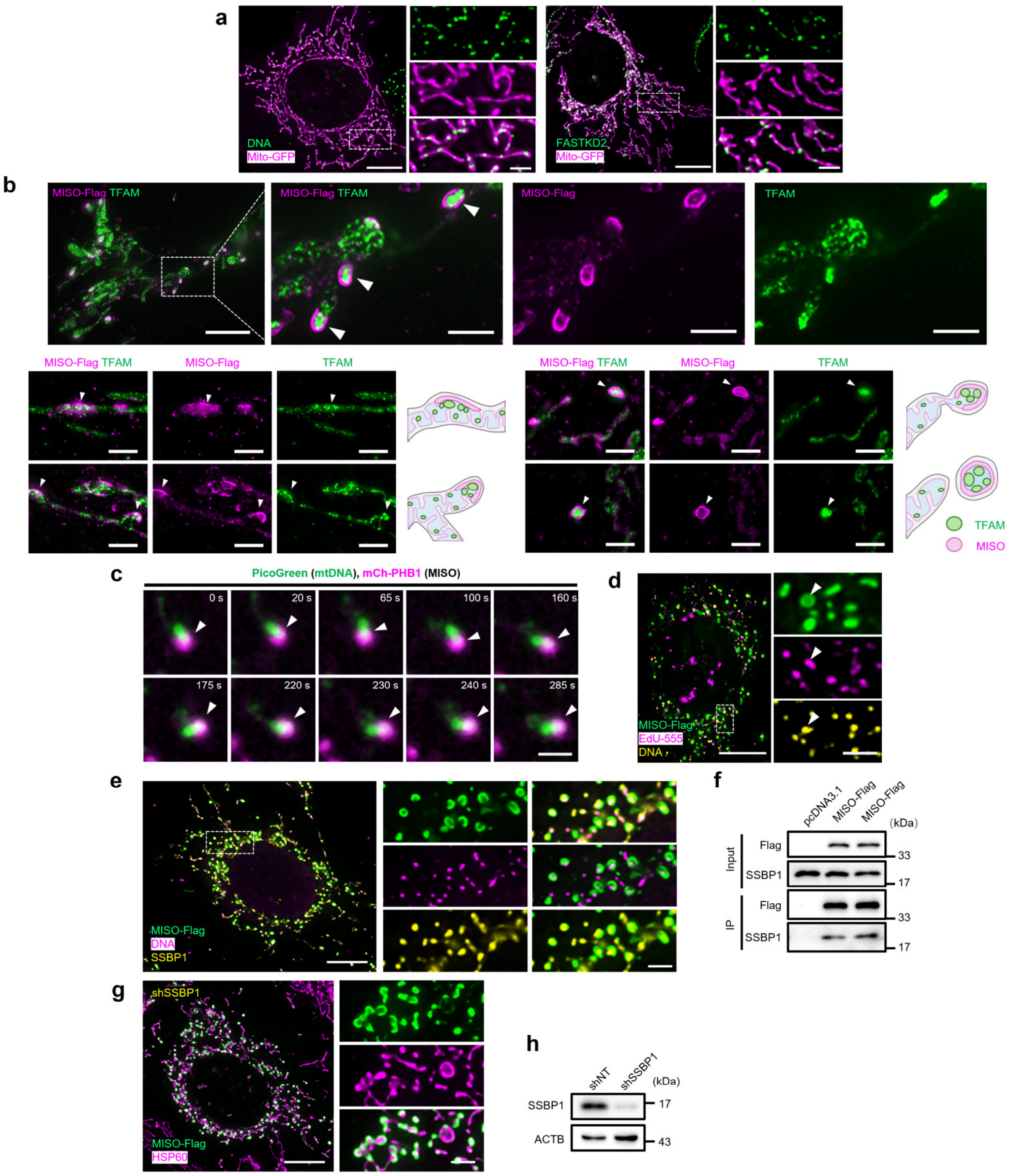


**Extended Data Fig. 18. MISO-enriched subdomains recruit mitochondrial nucleoids.**

**a.** Co-staining of the mitochondrial matrix-localized proteins Mito-GFP and mtDNA or mtRNA (indicating with FASTKD2). Scale bars: main panels 10 µm, magnified insets 2 µm.

**b.** Representative images showing the spatial association between mtDNA and MISO-enriched subdomains in U2OS cells with or without EB treatment. Scale bars: main panels 10 µm, magnified insets 2 µm.

**c.** Quantification of DNA enrichment in subdomains from (**b**).

**d.** Representative expansion microscopy images depicting the association between MISO with TFAM in U2OS cells. Scale bars: Top panels, main panels 20 μm (post-expansion), magnified insets 5 μm (post-expansion); bottom panels, 5 μm (post-expansion).

**e.** Representative time-lapse images of mitochondrial DNA (indicated by PicoGreen) and PHB1-mCherry (indicating MISO) in U2OS cells. White arrowheads points to the subdomains. Scale bar: 2 μm.

**f.** Representative images of MISO-enriched subdomains, mtDNA and replicating DNA (EdU) in U2OS cells expressing MISO-Flag. Scale bars: main panels 10 μm, magnified insets 2 μm.

**g.** Representative images of MISO-enriched subdomains, mtDNA and SSBP1 in U2OS cells expressing MISO-Flag. Scale bars: main panels 10 μm, magnified insets 2 μm.

**h.** Immunoprecipitation with anti-Flag validates the interaction between MISO and SSBP1.

**i.** Representative images of mitochondrial subdomains in U2OS cells expressing MISO-Flag with SSBP1 knockdown.

**i.** Immunoblots showing the knockdown efficiency of SSBP1 in U2OS cells.

All data are presented as mean ± SD. (**b**): non-parametric Mann-Whitney test.

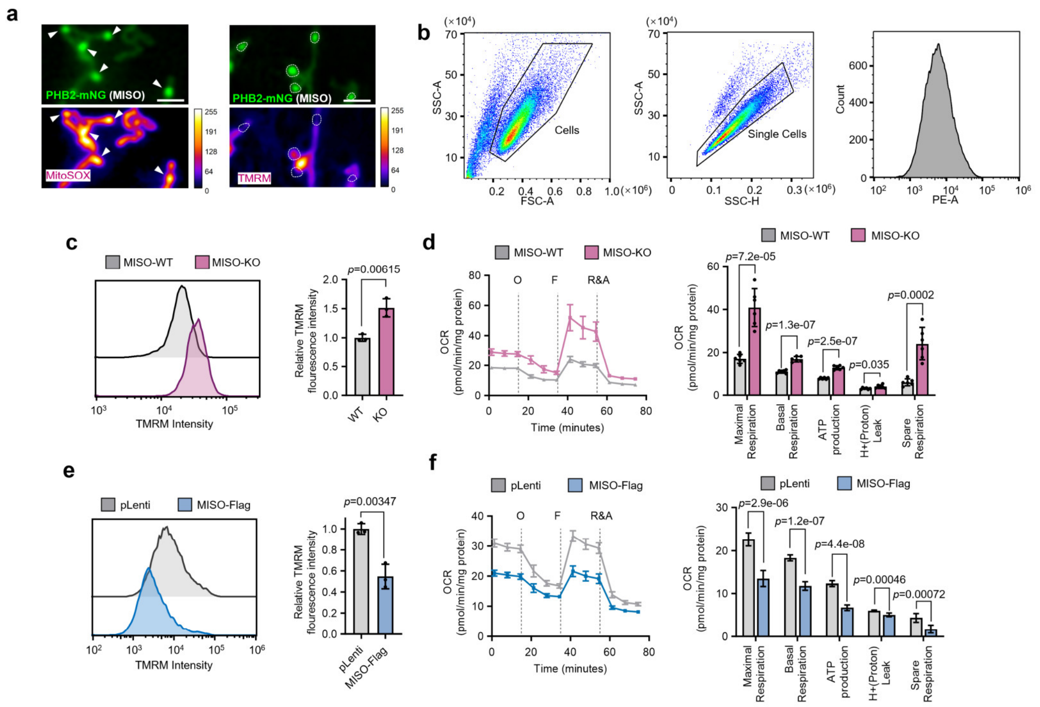


**Extended Data Fig. 19. MISO modulates both mitochondrial OXPHOS activity and membrane potential.**

**a.** Representative images of U2OS cells expressing PHB2-mNeonGreen and MISO-Flag incubated with MitoSOX or TMRM. MitoSOX or TMRM intensities were represented by calibrated color scale. Scale bars, 2 μm.

**b.** The gating strategy for determining TMRM fluorescence intensity.

**c.** Flow analysis and quantification of TMRM fluorescence intensity in WT and MISO-KO U2OS cells. *n* = three experiments.

**d.** Seahorse analyses of the mitochondrial respiratory capacity of WT and MISO-KO U2OS cells. O: Oligomycin; F: FCCP; R&A: Rotenone and Antimycin. *n* = six experiments.

**e.** Flow cytometry analysis and quantification of TMRM fluorescence intensity in U2OS cells expressing an empty vector (pLenti) or MISO-Flag. *n* = three experiments.

**f.** Seahorse analyses of the mitochondrial respiratory capacity of U2OS cells expressing an empty vector (pLenti) or MISO-Flag. O: Oligomycin; F: FCCP; R&A: Rotenone and Antimycin. *n* = six experiments.

All data are presented as mean ± SD. (**c-f**): two-tailed unpaired t test.

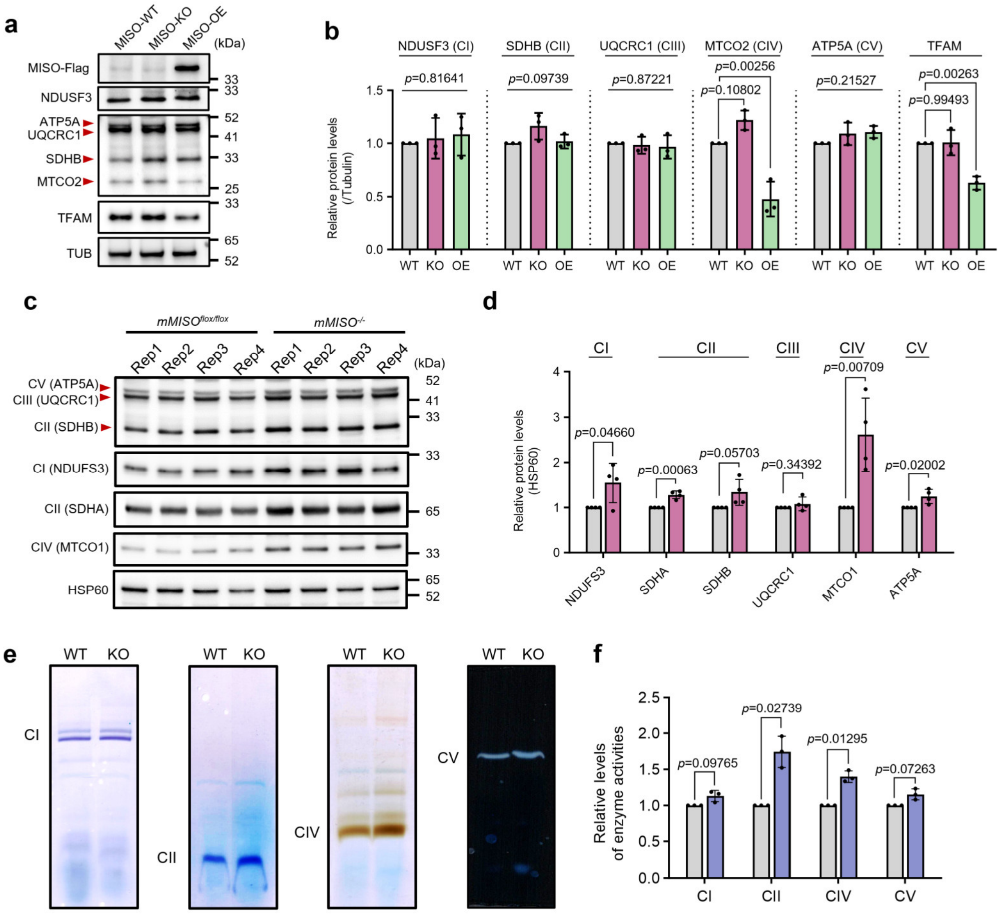


**Extended Data Fig. 20. MISO modulates the level and activity of OXPHOS complexes.**

**a-b.** Immunoblots (**a**) and corresponding quantifications (**b**) of indicated proteins in MISO-WT, MISO-KO and MISO-OE U2OS cells. *n* = three experiments.

**c-d.** Immunoblots (**c**) and corresponding quantifications (**d**) of mitochondrial OXPHOS complexes in liver tissues from *mMISO*$^{flox/flox}$ and *mMISO*$^{-/-}$ mice. Each lane represents an individual mouse.

**e-f.** Blue native PAGE-based in-gel activity assays (**e**) and corresponding quantification (**f**) of mitochondrial respiratory complexes I, II, IV, and V in WT and MISO-KO U2OS cells. *n* = three experiments.

All data are presented as mean ± SD. (**b**): ordinary one-way ANOVA with Tukey's multiple comparisons test; (**d**) and (**f**): two-tailed unpaired t test.

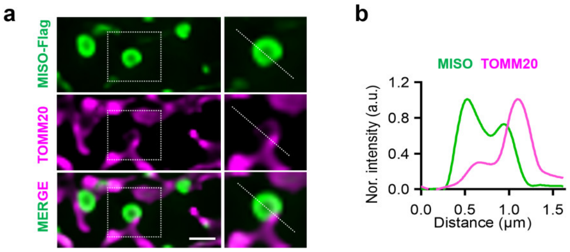


**Extended Data Fig. 21. MISO-enriched subdomains localize to the ends of mitochondria lack outer membrane coverage.**

**a.** Representative image MISO-enriched subdomains with loss of TOMM20. Scale bar: 1 μm.

**b.** A line profiles of MISO-enriched subdomain with loss of TOMM20.

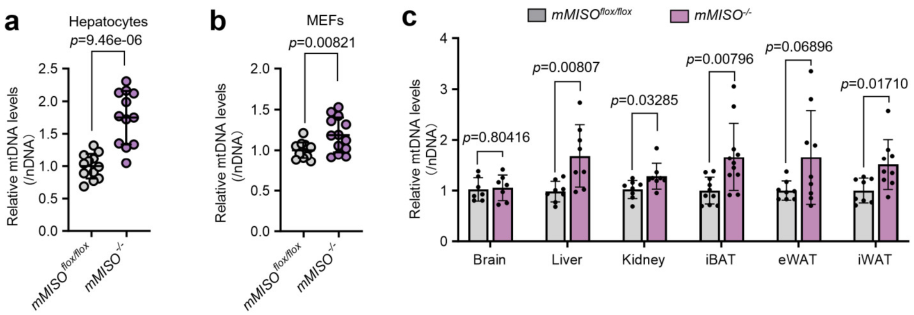


**Extended Data Fig. 22. MISO knockout increases mtDNA copy number.**

**a.** qPCR analysis of mtDNA copy number in *mMISO*$^{flox/flox}$ and *mMISO*$^{-/-}$ Hepatocytes. *n* = 12 experiments.

**b.** qPCR analysis of mtDNA copy number in *mMISO*$^{flox/flox}$ and *mMISO*$^{-/-}$ MEFs. *n* = four experiments.

**c.** qPCR analysis of mtDNA copy number in tissues from wild-type and mMISO knockout (KO) mice. *n* = three experiments.

All data are presented as mean ± SD. (**a**), (**b**) and (**c**): two-tailed unpaired t test.

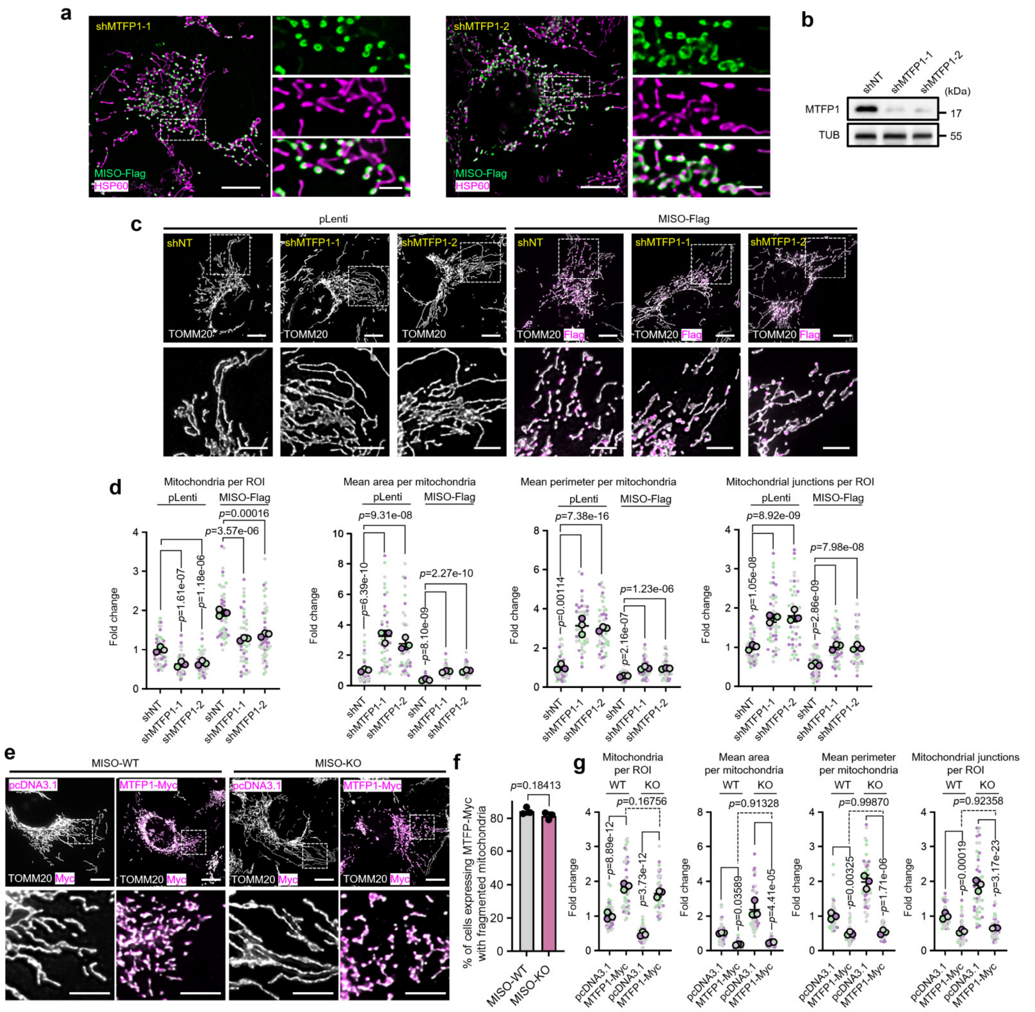


**Extended Data Fig. 23. Mitochondrial fission, but not SMEM formation, requires MTFP1.**

**a.** Representative images depicting mitochondrial subdomains in U2OS cells expressing MISO-Flag with MTFP1 knockdown. Scale bars: main panels 10 μm, magnified insets 2 μm.

**b.** Immunoblots showing the knockdown efficiency of two distinct shRNA constructs targeting MTFP1 in U2OS cells.

**c.** Representative images of mitochondrial morphology in U2OS cells treated with the indicated shRNAs and overexpressing pLenti or MISO-Flag. Scale bars: main panels 10 μm, magnified insets 5 μm.

**d.** Quantification of mitochondrial parameters from (**c**). *n* = three experiments.

**e-f.** Representative images (**e**) and corresponding quantification (**f**) of mitochondrial morphology in WT and MISO-KO U2OS cells expressing an empty vector (pcDNA3.1) or MTFP1-Myc. Scale bars: main panels 10 μm, magnified insets 5 μm. *n* = three experiments.

**g.** Quantification of mitochondrial parameters from (**e**). *n* = three experiments.

All data are presented as mean ± SD. (**d**) and (**g**): two-tailed nested t test; (**f**): two-tailed unpaired t test.

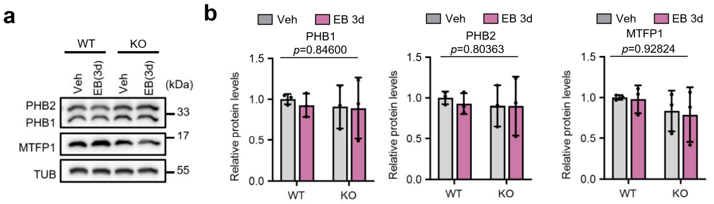


**Extended Data Fig. 24. EB treatment and MISO knockout do not alter the expression levels of other SMEM proteins.**

**a-b.** Representative immunoblots (**a**) and corresponding quantifications (**b**) of indicated proteins from WT and MISO-KO U2OS cells incubated in the presence or absence of EB. *n* = three experiments.

All data are presented as mean ± SD. (**b**): two-way ANOVA with Tukey's multiple comparisons test.

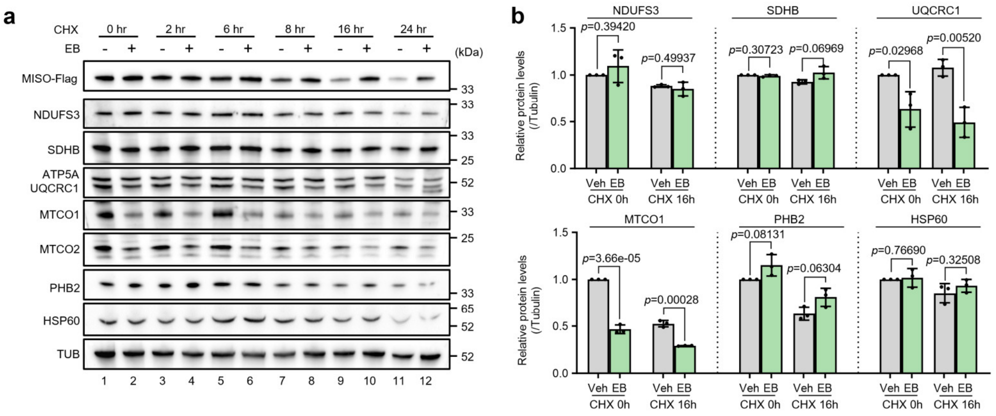


**Extended Data Fig. 25. EB treatment stabilizes MISO protein levels while reducing OXPHOS complex levels.**

**a.** CHX chase assay assessing MISO protein stability in U2OS cells treated in the presence or absence of EB.

**b.** Quantification of indicated protein levels from (**a**). $n$ = three experiments.

All data are presented as mean ± SD. (**b**): two-tailed unpaired t test.

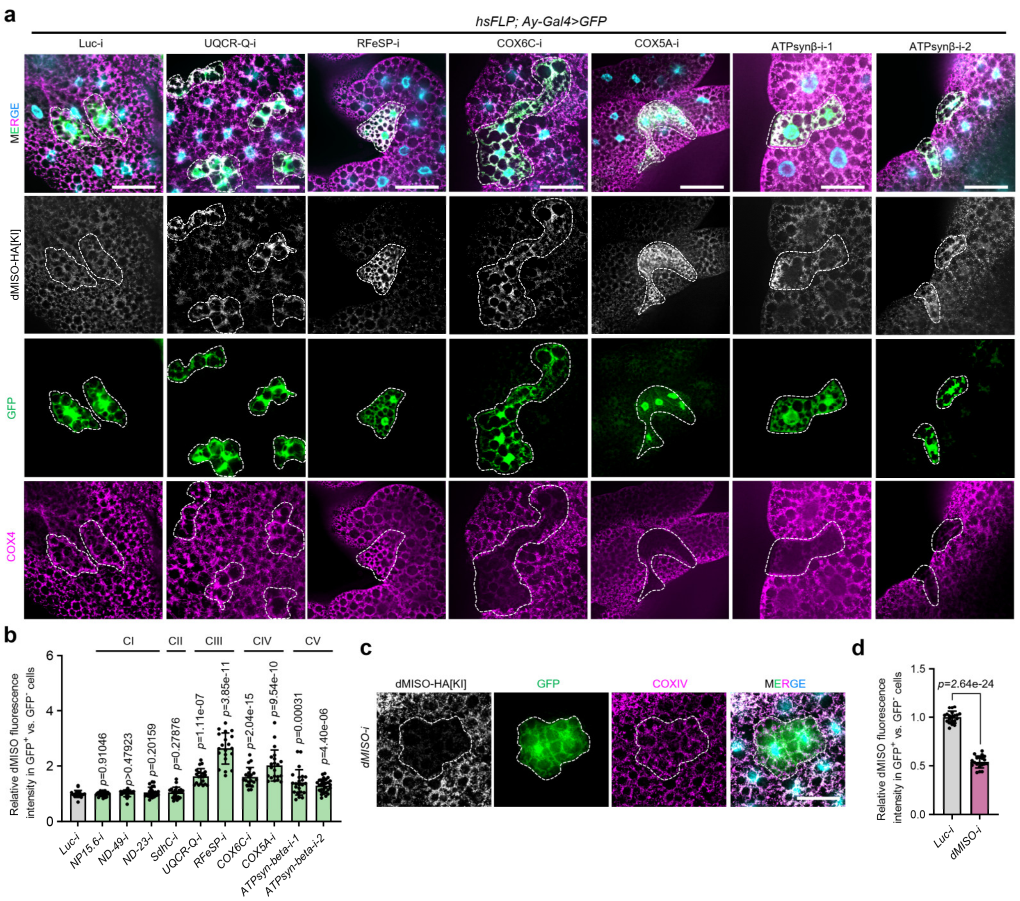


**Extended Data Fig. 26. OXPHOS complex knockdown induces MISO upregulation in *Drosophila* adipose tissue.**

**a.** Representative images of the *Drosophila* larval fat body harboring GFP-marked somatic clones of the indicated genotypes, immunostained for HA-tagged endogenous dMISO. Scale bars: 50 μm.

**b.** Quantification of relative MISO fluorescence intensity from (**a**).

**c.** Representative confocal micrographs of the *Drosophila* larval fat body harboring GFP-marked somatic clones of *dMISO* knockdown. Scale bars: 50 μm.

**d.** Quantification of relative MISO fluorescence intensity from (**c**).

All data are presented as mean ± SD. (**b**) and (**d**): two-tailed nested t test.

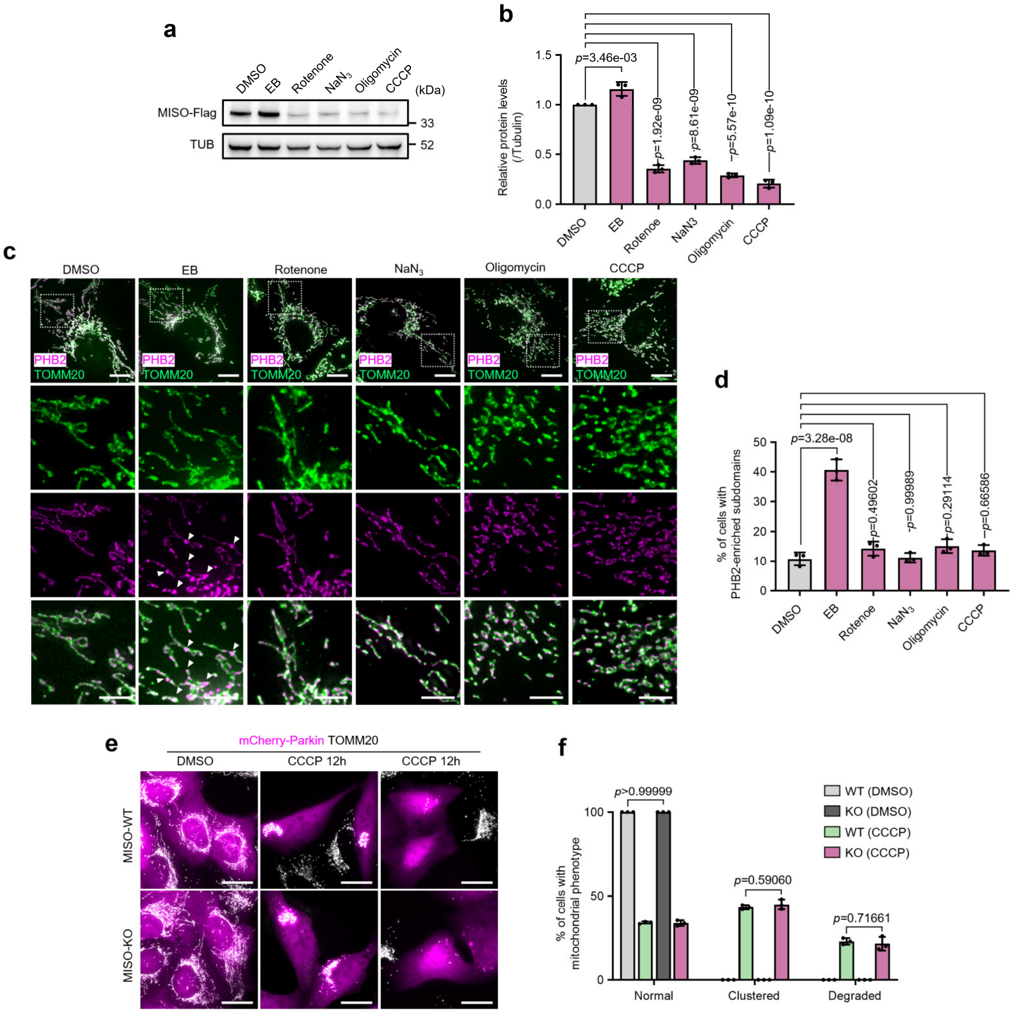


**Extended Data Fig. 27. MISO and SMEM are not induced by common mitochondrial stressors and does not participate in classic Parkin-mediated mitophagy.**

**a-b.** Immunoblots (**a**) and corresponding quantifications (**b**) of MISO protein levels in U2OS cells treated for 24 hours with the indicated compounds (1 μM Rotenone, 5 mM $NaN_3$, 2 μM Oligomycin and 2 μM CCCP), except for EB, which was applied at 50 ng/mL for 3 days. Relatively low dosages of the compounds were used to avoid triggering significant mitophagy. *n* = three experiments.

**c-d.** Representative images (**c**) and corresponding quantifications (**d**) of PHB2-enriched microdomains (white arrows) in WT and MISO-KO U2OS cells treated with the indicated compounds as described in (**a-b**). Scale bars: main panels 10 μm, magnified insets 5 μm. *n* = three experiments.

**e-f.** Representative images (**e**) and corresponding quantification (**f**) of mitochondrial phenotype in WT and MISO-KO U2OS cells expressing mCherry-Parkin, treated with or without 10 μM CCCP for 12 hours. Scale bars: main panels 20 μm, magnified insets 5 μm. *n* = three experiments.

All data are presented as mean ± SD. (**b**) and (**d**): ordinary one-way ANOVA with Tukey's multiple comparisons test; (**f**): two-way ANOVA with Tukey's multiple comparisons test.

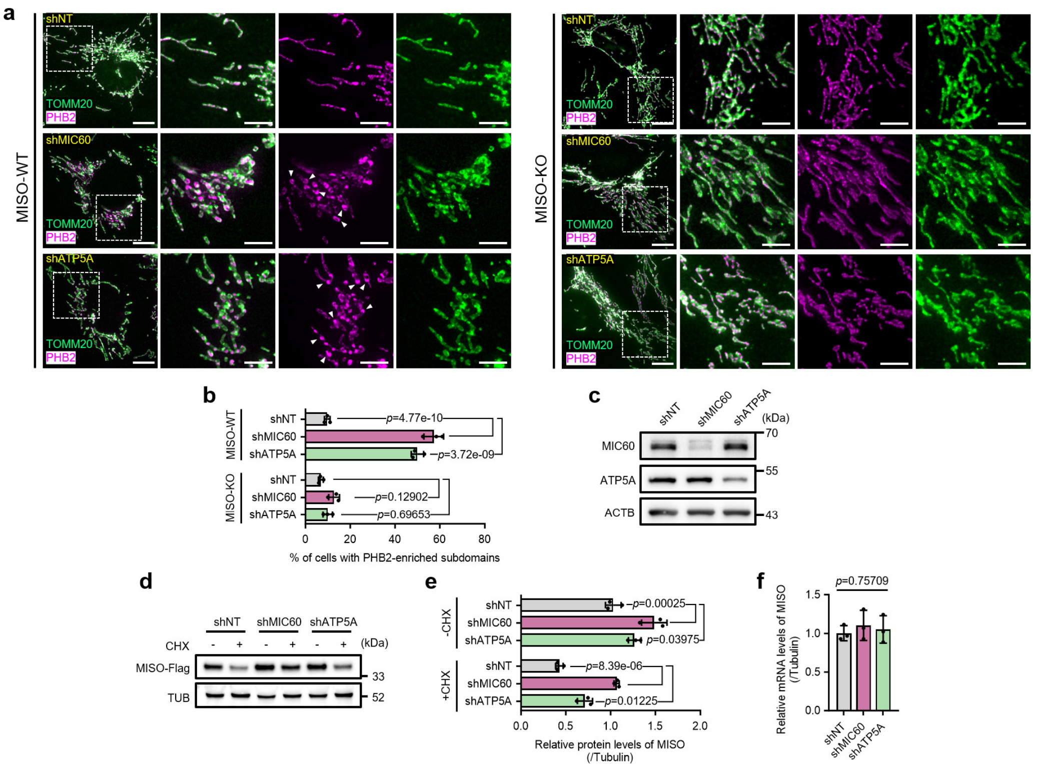


**Extended Data Fig. 28. MIC60 and ATP5A depletion promotes SMEM formation through a MISO-dependent pathway.**

**a-b.** Representative images (**a**) and corresponding quantifications (**b**) of PHB2-enriched microdomains (white arrows) in WT and MISO-KO U2OS cells treated with indicated shRNAs. Scale bars: main panels 10 μm, magnified insets 5 μm. *n* = three experiments.

**c.** Immunoblots showing the knockdown efficiency of MIC60 and ATP5A in U2OS cells.

**d-e.** Immunoblots (**d**) and corresponding quantifications (**e**) from CHX chase assay assessing MISO protein stability in U2OS cells stably expressing MISO and transduced with the indicated shRNAs. *n* = three experiments.

**f.** qPCR analysis of MISO mRNA levels in U2OS cells transduced with the indicated shRNAs. *n* = three experiments.

All data are presented as mean ± SD. (**b**): two-way ANOVA with Tukey's multiple comparisons test; (**e**) and (**f**): ordinary one-way ANOVA with Tukey's multiple comparisons test.

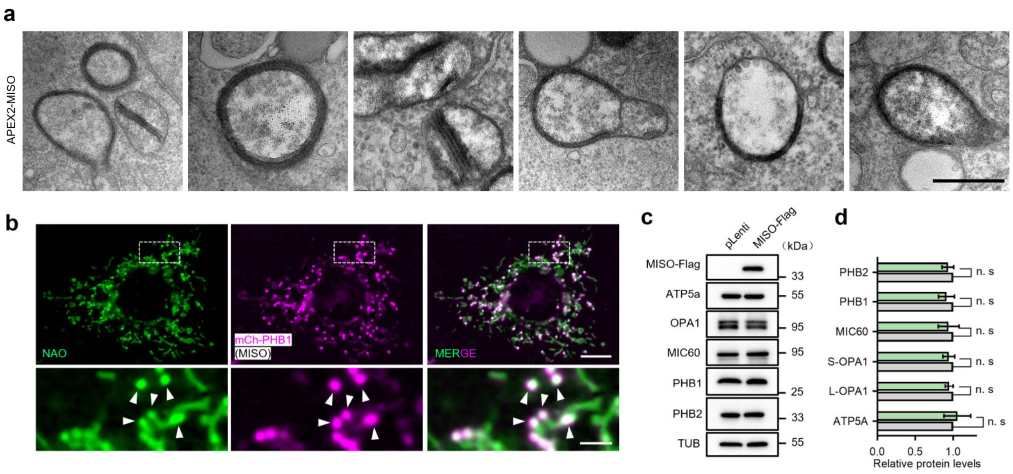


**Extended Data Fig. 29. SMEM are formed from collapsed and condensed inner membranes without altering the protein levels of major cristae-organizing proteins.**

**a.** Representative TEM images of mitochondria in U2OS cells expressing APEX2-MISO. Scale bar: 500 nm.

**b.** Representative images of U2OS cells expressing PHB1-mCherry and MISO-Flag, incubated with nonyl acridine orange (NAO). White arrows indicate subdomains. Scale bars: main panels 10 µm, magnified insets 2 µm.

**c-d.** Representative immunoblots (**c**) and corresponding quantification (**d**) of proteins involved in the maintenance of mitochondrial cristae structure and morphology from U2OS cells expressing an empty vector (pLenti) or MISO-Flag. Tubulin was used as a loading control. *n* = three experiments.

All data are presented as mean ± SD. (**d**): two-tailed unpaired t test. n. s: not significant.

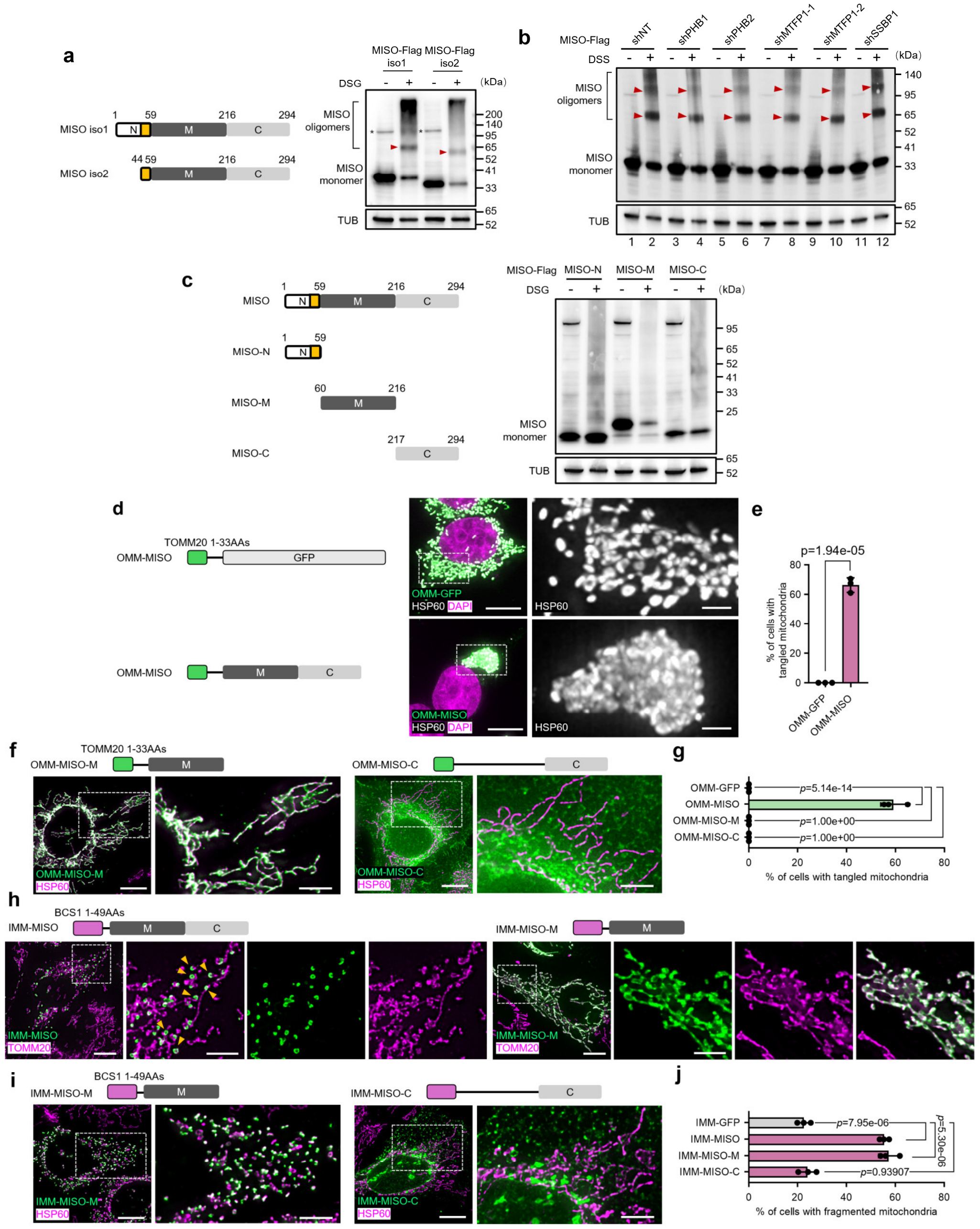


**Extended Data Fig. 30. The oligomerization and subcellular localization of different MISO variants.**

**a.** Crosslinking analysis of the oligomerization of two MISO isoforms in U2OS cells using disuccinimidyl glutarate (DSG). Red arrowheads point to the MISO dimers. Asterisks denote nonspecific bands.

**b.** Crosslinking analysis of MISO oligomerization using disuccinimidyl glutarate (DSG) in U2OS cells expressing MISO-Flag and treated with the indicated shRNAs. Red arrowheads point to the MISO dimers and trimers.

**c.** Crosslinking analysis of oligomerization of MISO and corresponding truncation variants in U2OS cells.

**d-e.** Representative images and corresponding quantification (**e**) of mitochondrial morphology in HEK293T cells expressing OMM-GFP or OMM-MISO. Scale bars: main panels 10 μm, magnified insets 2 μm. *n* = three experiments.

**f-g.** Representative images (**f**) and corresponding quantification (**g**) of mitochondrial morphology in U2OS cells expressing OMM-MISO truncation mutants. Scale bars: main panels 10 μm, magnified insets 5 μm. *n* = three experiments.

**h.** Representative images showing the localization of IMM-MISO and IMM-MISO-M in U2OS cells.

**i-j.** Representative images (**h**) and corresponding quantification (**e**) of mitochondrial morphology in U2OS cells expressing IMM-MISO truncation mutants. Scale bars: main panels 10 μm, magnified insets 5 μm. *n* = three experiments.

All data are presented as mean ± SD. (**e**): two-tailed unpaired t test; (**g**) and (**j**): ordinary one-way ANOVA with Tukey's multiple comparisons test.

**Extended Data Table 1：List of Fly Genotypes in the Figures**

| Figure | Panels | Genotype |
|---|---|---|
| Fig. 1 | Fig. 1b | *w; esg::GFP, CG30159-Gal4; UAS-myr::tdTomato* |
| | Fig. 1c | *w; esg::GFP, CG30159-Gal4; UAS-CG30159-3×HA* |
| | Fig. 1e | *w; esg-Gal4, tub-Gal80ts, UAS-GFP; UAS-CG30159-3×HA* |
| | Fig. 1g | *w, Su(H)Gbe-lacZ; esg::GFP/CG30159-3×HA[KI]* |
| | Fig. 1h | *w; esg-Gal4, tub-Gal80ts, UAS-GFP/CG30159-3×HA[KI]; UAS-mito-APEX-Flag* |
| | Fig. 1i | *w; esg-Gal4, tub-Gal80ts, UAS-GFP; UAS-mito-APEX-Flag/UAS-Luc RNAi* |
| | | *w; esg-Gal4, tub-Gal80ts, UAS-GFP; UAS-mito-APEX-Flag/UAS-CG30159 RNAi* |
| | Fig. 1k | *w; esg-Gal4, tub-Gal80ts, UAS-GFP; UAS-mito-APEX-Flag* |
| | | *w; esg-Gal4, tub-Gal80ts, UAS-GFP; UAS-mito-APEX-Flag/UAS-CG30159-3×HA* |
| Extended Data Fig. 1 | Ext Data Fig. 1a | *w; CG30159-Gal4; UAS-myr::tdTomato* |
| | Ext Data Fig. 1b | *w; CG30159-Gal4; UAS-myr::tdTomato* |
| | Ext Data Fig. 1c | *w; CG30159-Gal4; UAS-myr::tdTomato/Dl-LacZ* |
| | Ext Data Fig. 1d | *w, Su(H)Gbe-lacZ; CG30159-Gal4; UAS-myr::tdTomato* |
| Extended Data Fig. 26 | Ext Data Fig. 26a | *hsFlp; AyGal4, UAS-GFP/CG30159-3×HA[KI]; UAS-Luc RNAi* |
| | | *hsFlp; AyGal4, UAS-GFP/CG30159-3×HA[KI]; UAS-UQCR-Q RNAi* |
| | | *hsFlp; AyGal4, UAS-GFP/CG30159-3×HA[KI]; UAS-RFeSP RNAi* |
| | | *hsFlp; AyGal4, UAS-GFP/CG30159-3×HA[KI]; UAS-COX5A RNAi* |
| | | *hsFlp; AyGal4, UAS-GFP/CG30159-3×HA[KI]; UAS-COX6C RNAi* |
| | | *hsFlp;AyGal4, UAS-GFP/CG30159-3×HA[KI]; UAS-ATPsynβ RNAi-1* |
| | | *hsFlp;AyGal4, UAS-GFP/CG30159-3×HA[KI]; UAS-ATPsynβ RNAi-2* |
| | Ext Data Fig. 26b | *hsFlp; AyGal4, UAS-GFP/CG30159-3×HA[KI]; UAS-Luc RNAi* |
| | | *hsFlp; AyGal4, UAS-GFP/CG30159-3×HA[KI]; UAS-NP15.6 RNAi* |
| | | *hsFlp; AyGal4, UAS-GFP/CG30159-3×HA[KI]; UAS-ND-49 RNAi* |
| | | *hsFlp; AyGal4, UAS-GFP/CG30159-3×HA[KI]; UAS-ND-23 RNAi* |
| | | *hsFlp; AyGal4, UAS-GFP/CG30159-3×HA[KI]; UAS-SdhC RNAi* |
| | | *hsFlp; AyGal4, UAS-GFP/CG30159-3×HA[KI]; UAS-UQCR-Q RNAi* |
| | | *hsFlp; AyGal4, UAS-GFP/CG30159-3×HA[KI]; UAS-RFeSP RNAi* |
| | | *hsFlp; AyGal4, UAS-GFP/CG30159-3×HA[KI]; UAS-COX6C RNAi* |
| | | *hsFlp; AyGal4, UAS-GFP/CG30159-3×HA[KI]; UAS-COX5A RNAi* |
| | | *hsFlp;AyGal4, UAS-GFP/CG30159-3×HA[KI]; UAS-ATPsynβ RNAi-1* |
| | | *hsFlp;AyGal4, UAS-GFP/CG30159-3×HA[KI]; UAS-ATPsynβ RNAi-2* |
| | Ext Data Fig. 26c | *hsFlp; AyGal4, UAS-GFP/CG30159-3×HA[KI]; UAS-CG30159 RNAi* |
| | Ext Data Fig. 26d | *hsFlp; AyGal4, UAS-GFP/CG30159-3×HA[KI]; UAS-Luc RNAi* |
| | | *hsFlp; AyGal4, UAS-GFP/CG30159-3×HA[KI]; UAS-CG30159 RNAi* |

**Extended Data Table 2： List of fly RNAi lines tested for OXPHOS complexes.**

| | Stock ID | Reported phenotypes caused by the indicated RNAi line |
|---|---|---|
| Complex I | BL36672 | Knockdown of NP15.6 upregulates the expression of components of the tyrosine degradation pathway, which can decrease fly lifespan[1]; knockdown of NP15.6 in in wings causes larval lethality[2]; knockdown of NP15.6 in neurons activated glial senescence in the antennal lobes[3]; knockdown of ND49 in fat body tissues leads to enlarged mitochondria with reduced cristae density[4]. |
| | BL28573 | NDI1-induced NAD+/NADH ratio and cell viability changes were rescued by ND-49 RNAi[5]; knockdown of ND-49 in muscle drastically reduced the synthesis of complex I[6]. |
| | BL30487 | NDI1-induced NAD+/NADH ratio and cell viability changes were rescued by ND-49 RNAi[5]; knockdown of ND-49 in muscle drastically reduced the synthesis of complex I[6]. |
| Complex II | BL53281 | The expression of SdhC-RNAi blocked Hipk-induced tumor-like growth to a certain extent in lifespan[7]. |
| Complex III | BL51357 | The reduction of the expression of UQCR-Q in fat body led to severe swollen mitochondria[4]. |
| | NIG7361R-1 | The disruption of Rieske iron sulfur protein increases lysosomal and autophagic activity[8]. |
| Complex IV | BL33878 | Knockdown of COX6C in the eyes induces a glossy-eye phenotype through disruption of the electron transport chain and ATP synthase function[9]. |
| | NIG14724R-3 | Loss-of-function mutations in Complex IV subunit 5A resulted in severely impaired mitochondrial membrane potential, as demonstrated by mosaic analysis[10]. |
| Complex V | NIG11154R-1 | The ATP synthase β subunit RNAi flies showed accumulation of total α/β-tubulin and abnormal muscle structure, which similar to the phenotypes found in parkin mutants[11]. |
| | BL28062 | Reduced expression of ATPsynB in the eyes leads to a glossy eye phenotype[12]; knockdown of ATPsynB leads to swollen mitochondria in the third instar larval fat body[4]. |

**Supplementary Video 1**

**Translocation of SMEM to the mitochondrial periphery.** Time-lapse imaging of U2OS cells stably expressing PHB2-mNeonGreen, MISO-3×Flag, and mito-RFP. Scale Bar: 2 μm.

**Supplementary Video 2**

**Representative live imaging of type I and type II fission events relative to SMEM.** Time-lapse imaging of U2OS cells stably expressing PHB2-mNeonGreen, MISO-3×Flag, and mito-RFP. Type I fission: Fission event occurring within a 0-1 μm range from the SMEM spot. Type II fission: Fission event occurring within a 1-2 μm range from the SMEM spot. Scale bar: 2 μm.

**Supplementary Video 3**

**Detachment of SMEM from mitochondria through peripheral fission.** Time-lapse imaging of U2OS cells stably expressing PHB2-mNeonGreen, MISO-3×Flag, and mito-RFP. Scale Bar: 2 μm.

**Supplementary Video 4**

**Representative live imaging of type I, II, and III mitochondrial fusion events.** Time-lapse imaging of U2OS cells stably expressing PHB2-mNeonGreen, MISO-3×Flag, and mito-RFP. Type I fusion: Fusion between two mitochondrial termini without SMEM. Type II fusion: Fusion between two mitochondrial termini with SMEM present on one terminus. Type III fusion: Fusion between two mitochondrial termini with SMEM present on both termini. Scale Bar: 2 μm.

**Supplementary Video 5**

**Live imaging of mtDNA and SMEM. Description:** U2OS cells stably expressing PHB1-mCherry and MISO-3×Flag were stained with PicoGreen to visualize mtDNA. Scale Bar: 2 μm.
